# European admixture in Chinchorro DNA

**DOI:** 10.1101/132555

**Authors:** Robert Smith

## Abstract

It is widely held that, except for migrations from Beringia or Siberia, there was no contact between the Old World and the New World prior to the colonization of North America by the Norse in the late 10^th^ century AD. Analyses of 23 ancient American DNA samples reveal, however, the presence of European admixture in a sample taken from a Chinchorro mummy of northern Chile dated to 3972–3806 BC. This discovery implies a more complex history of the peopling of the Americas than previously accepted.

## Introduction

Mainstream accounts of the peopling of the Americas stipulate that there was a large migration of people from Beringia during the late Pleistocene, and that modern Amerindians derive most of their ancestry from these migrants. It is commonly believed that these migrants were the originators of the Clovis material culture, and it was long asserted that the appearance of these migrants south of Beringia did not predate the earliest datings of Clovis artifacts, around 13,200 years ago. The accumulation of evidence for earlier American settlements has forced mainstream scholars to abandon the latter position, however. Mainstream accounts also acknowledge two later migrations from Siberia, responsible for bringing speakers of Na-Dene and Eskimo-Aleut languages to the Americas.

A small minority of researchers have advanced theories of other migrations to the Americas. Dennis Stanford and Bruce Bradley, observing the many similarities between Clovis technology and that of the Upper Paleolithic Solutrean culture of Western Europe, have theorized that the ancestors of the Clovis people came not from Siberia, but from Europe, by traversing the Atlantic Ocean during the Last Glacial Maximum [1, 2]. In his 1947 crossing of the Pacific on the balsa raft *Kon-Tiki* [3], and his 1970 crossing of the Atlantic on the reed boat *Ra II* [4], Thor Heyerdahl demonstrated the feasibility of early transoceanic contacts, and in his written works he produced an abundance of historical, archeological, and anthropological evidence that such contacts had indeed taken place [5, 6, 7].

Mainstream scholars have continually rejected these other proposed migrations to the Americas, in spite of the evidence from a multitude of scientific disciplines in support of them. Ancient DNA evidence, however, can prove incontrovertibly that these other migrations took place, and such evidence is presented here for the first time.

## Methods

### Data

The 23 ancient American genomes analyzed are those published in [8]. Genotype calls generated from aligned reads were intersected with a set of 110,817 transversion SNPs (set 1) for principal component analysis, and a set of 228,841 transversion SNPs (set 2) for *qpAdm* and ADMIXTURE analyses. The resultant numbers of SNPs for each of the ancient American samples are shown in Table 1.

**Table 1.**
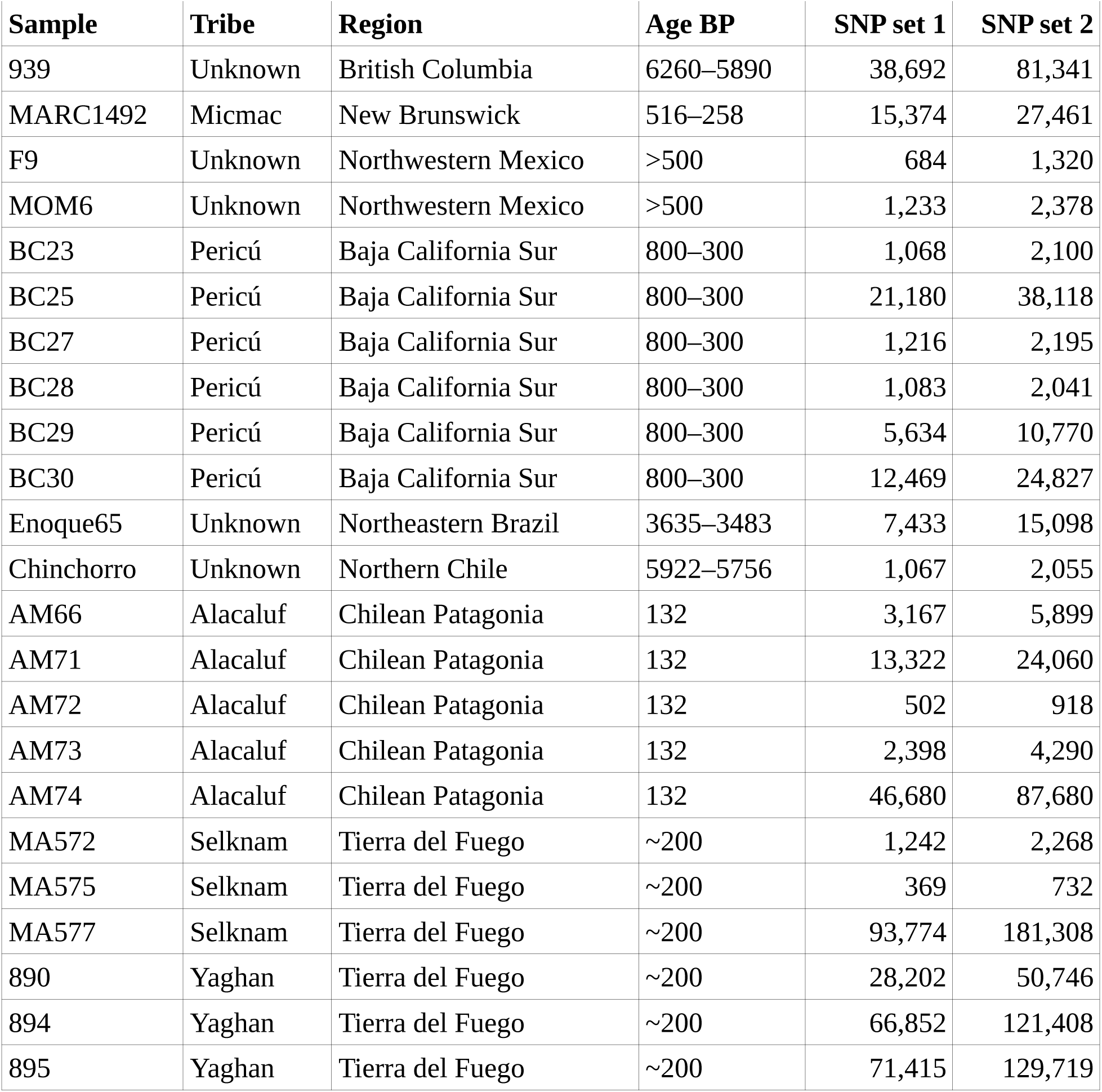
Description and numbers of SNPs used for 23 ancient American samples.

### *qpAdm* analysis

The program *qpAdm* [9] was used to estimate mixture coefficients for the ancient American samples, using the following populations:
- Target population: One of the 23 ancient American samples.
- Source population 1: A collection of populations shown to be purely Amerindian in admixture analyses: Mixe, Piapoco, Wichi, Chané, Karitiana, and Surui.
- Source population 2: One of 11 ancient or modern European, Middle Eastern, or South Asian populations.
- Outgroup populations: Mbuti Pygmies, Han Chinese, Nganasans, Eskimos, and Papuans.

### Principal component analysis

Principal component analysis of the ancient American samples was performed using the *smartpca* program of EIGENSOFT [10], using default parameters and the lsqproject: YES and numoutlieriter: 0 options.

### ADMIXTURE analysis

Model-based clustering analysis of the ancient American samples was performed with the ADMIXTURE program [11], with the number of assumed ancestral populations ranging from *K* = 4 to *K* = 13.

## Results

### *qpAdm* analysis

The results of the of the 253 *qpAdm* analyses are shown in Table 2, which gives the mean source population 2 mixture coefficient, and the coefficients plus or minus one standard error, for each analysis. The analyses are listed in decreasing order of the mixture coefficient lower bounds.

**Table 2.**
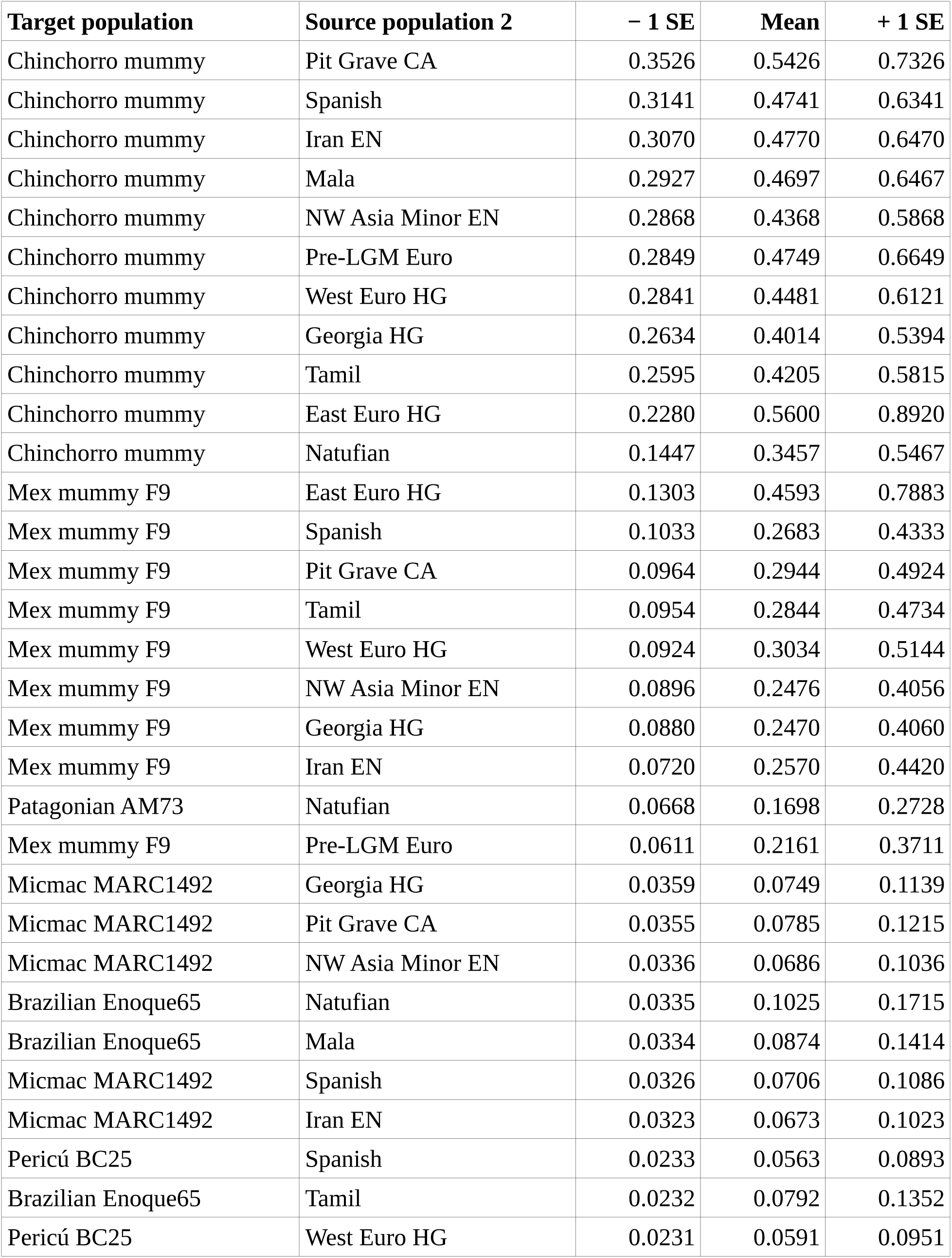
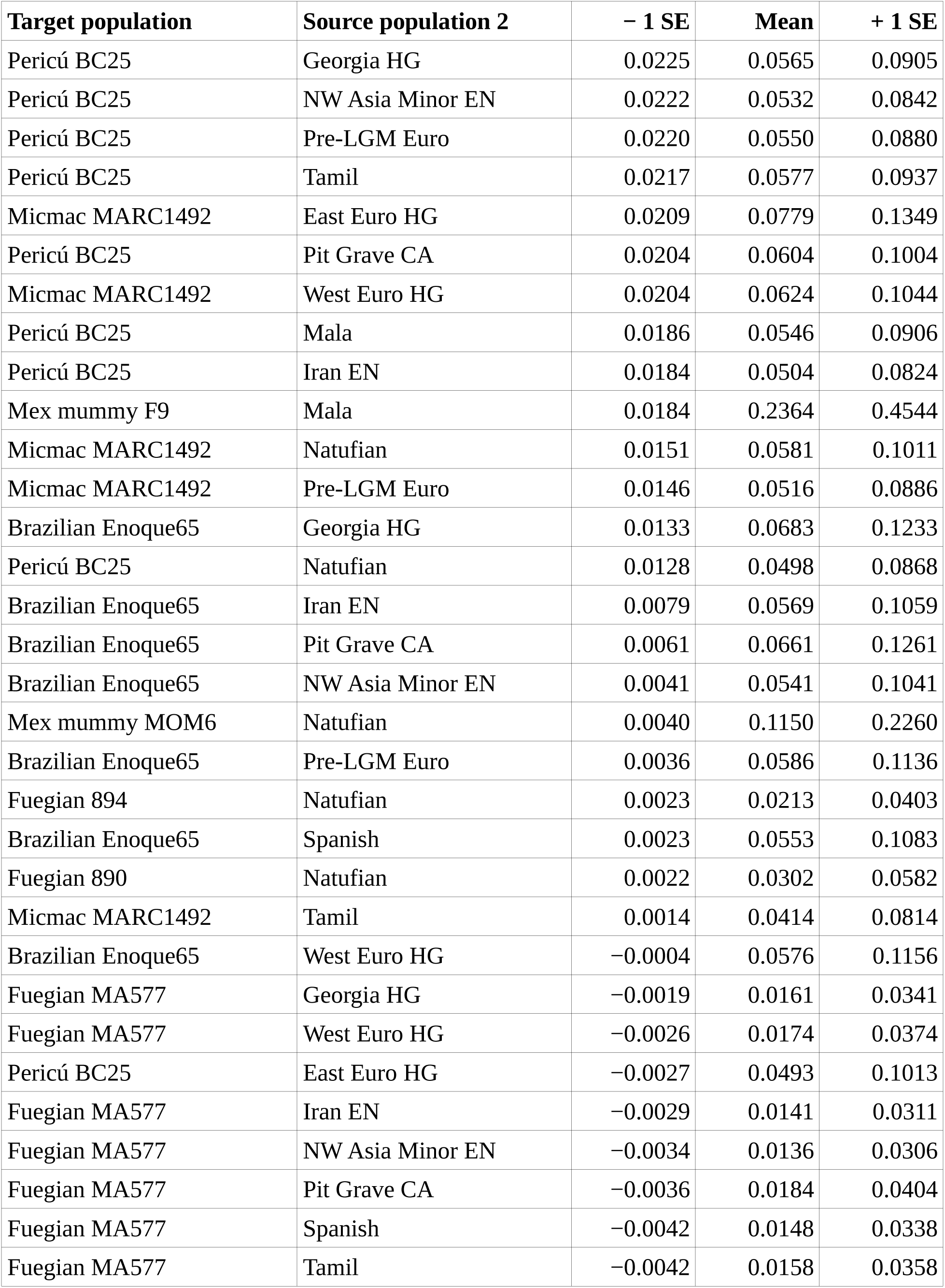
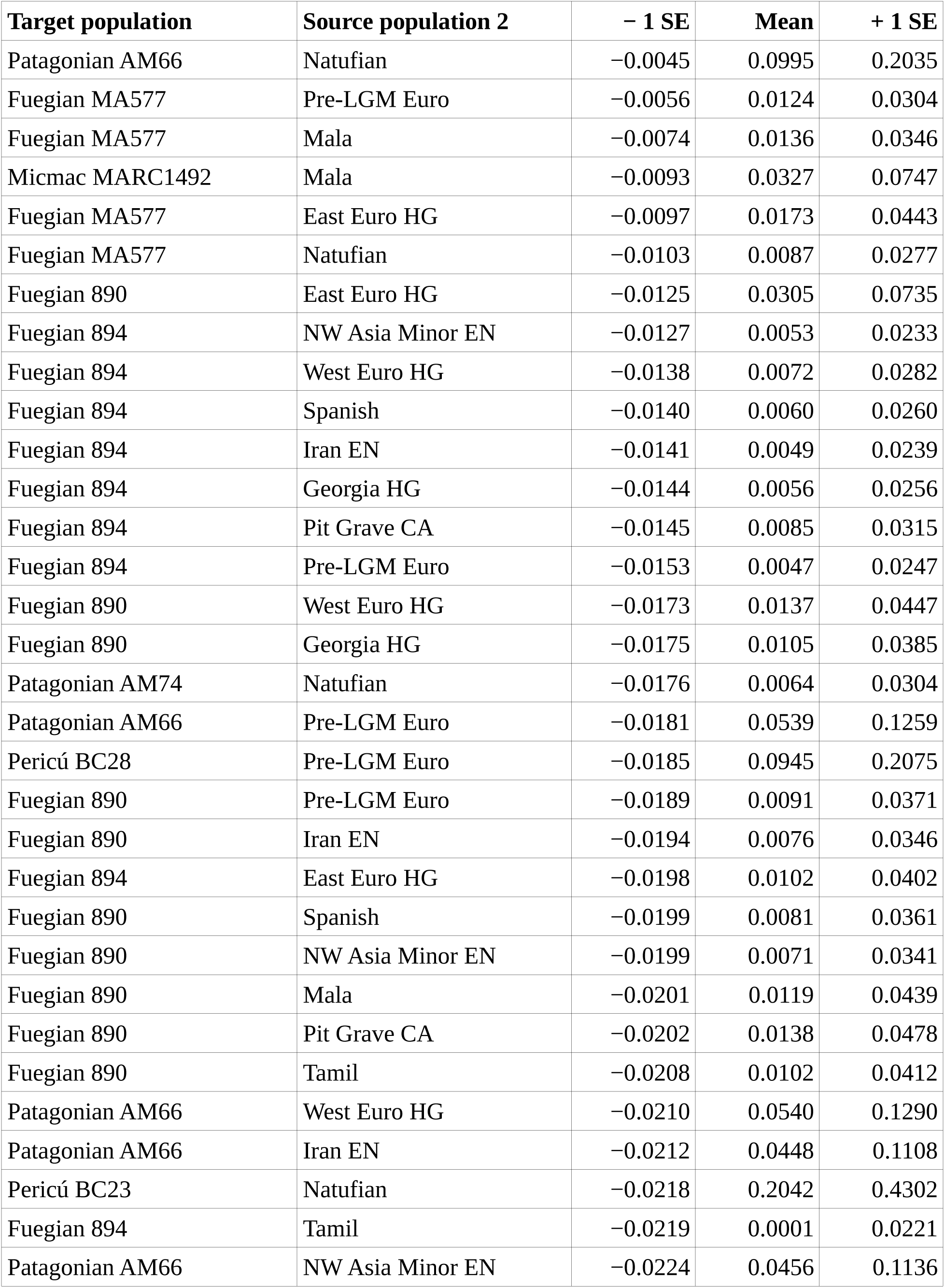
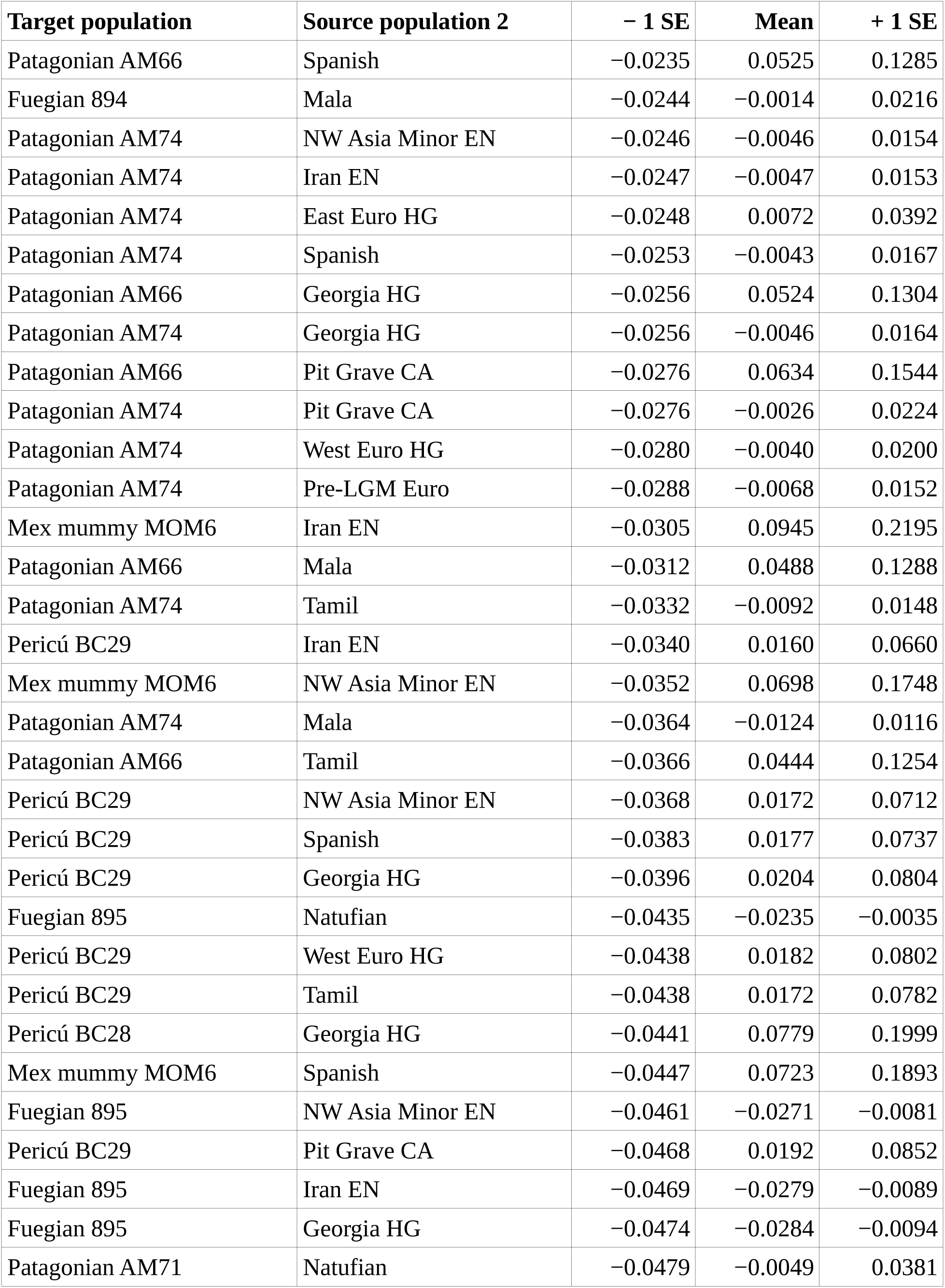
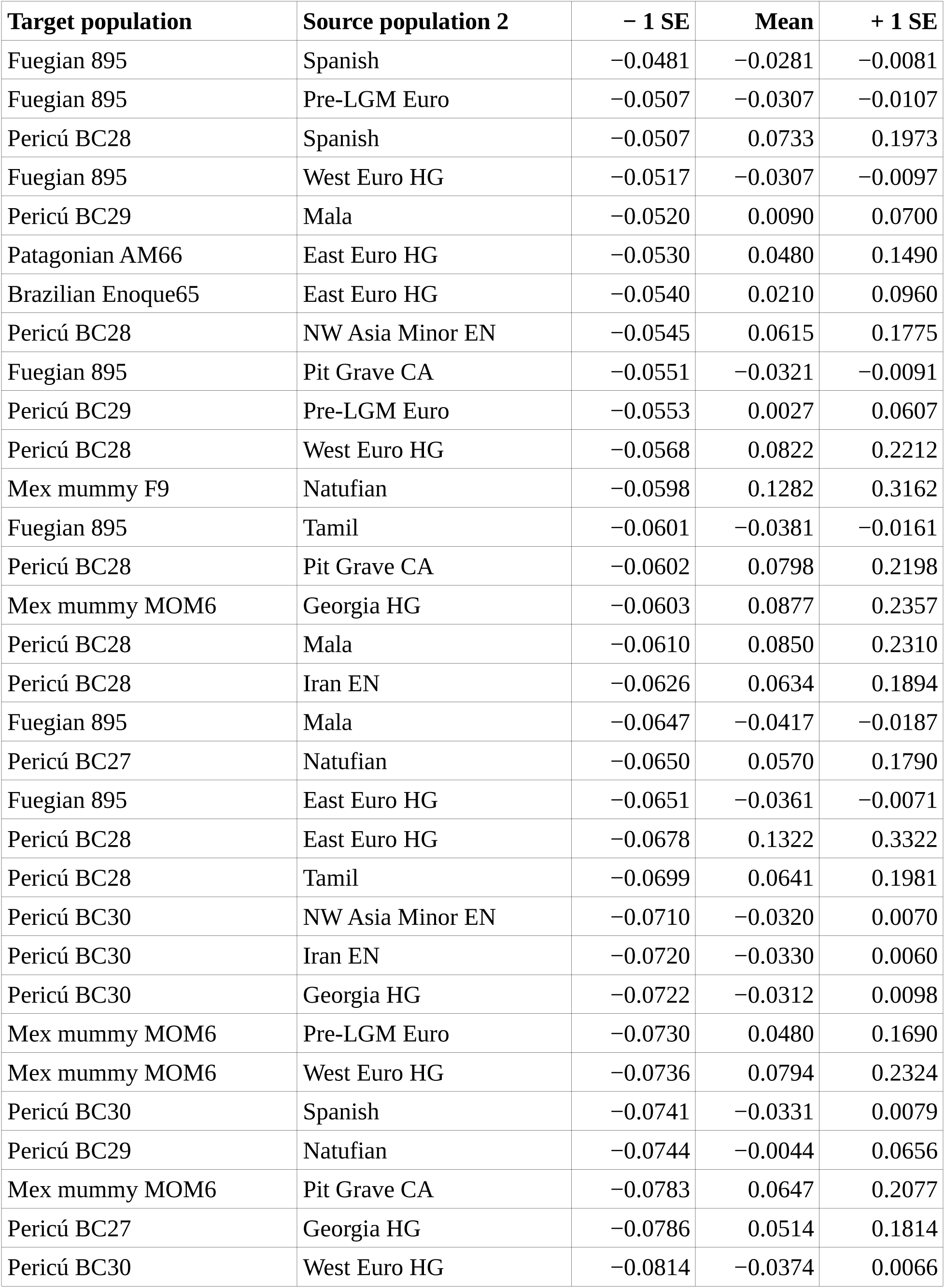
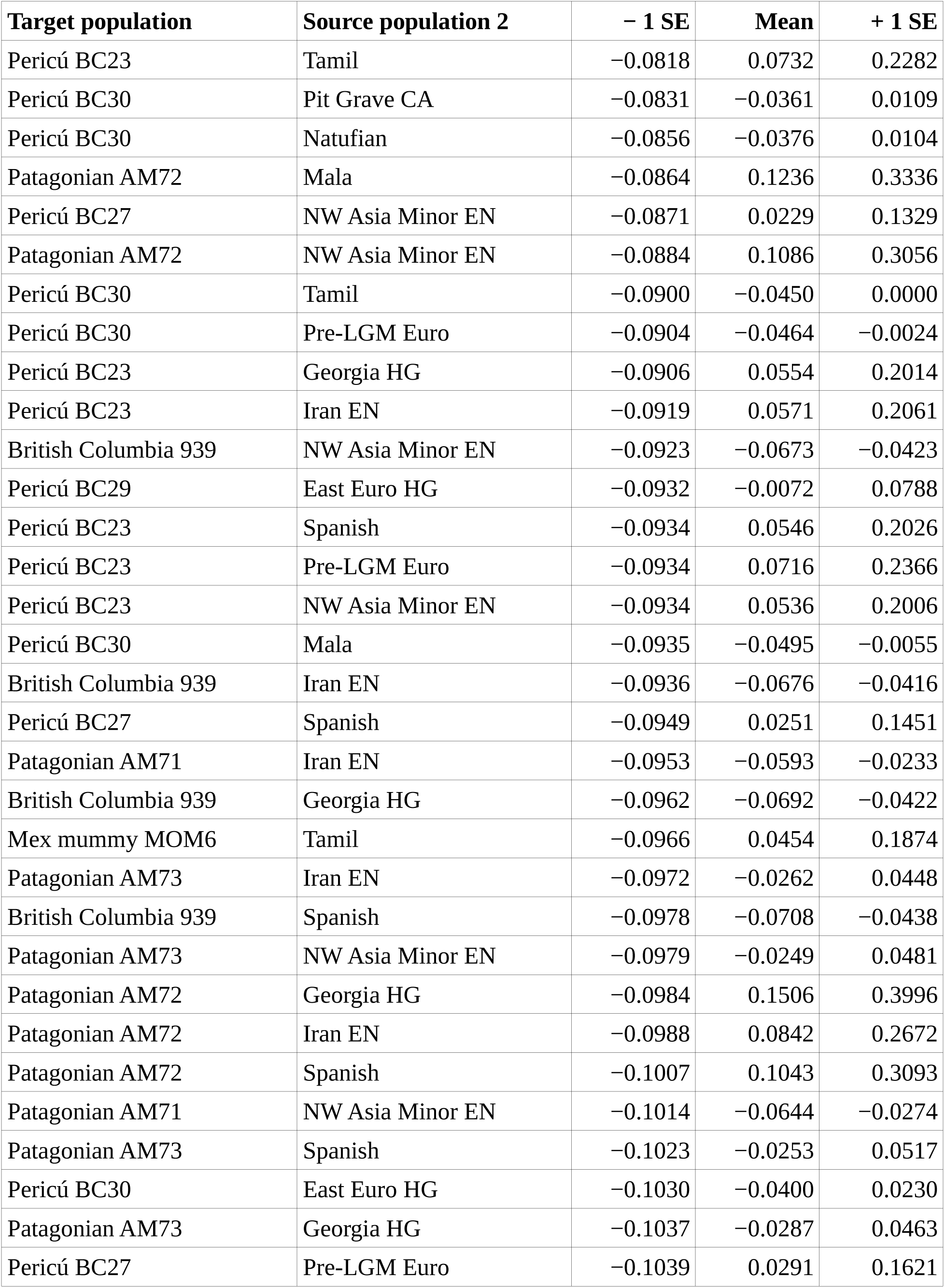
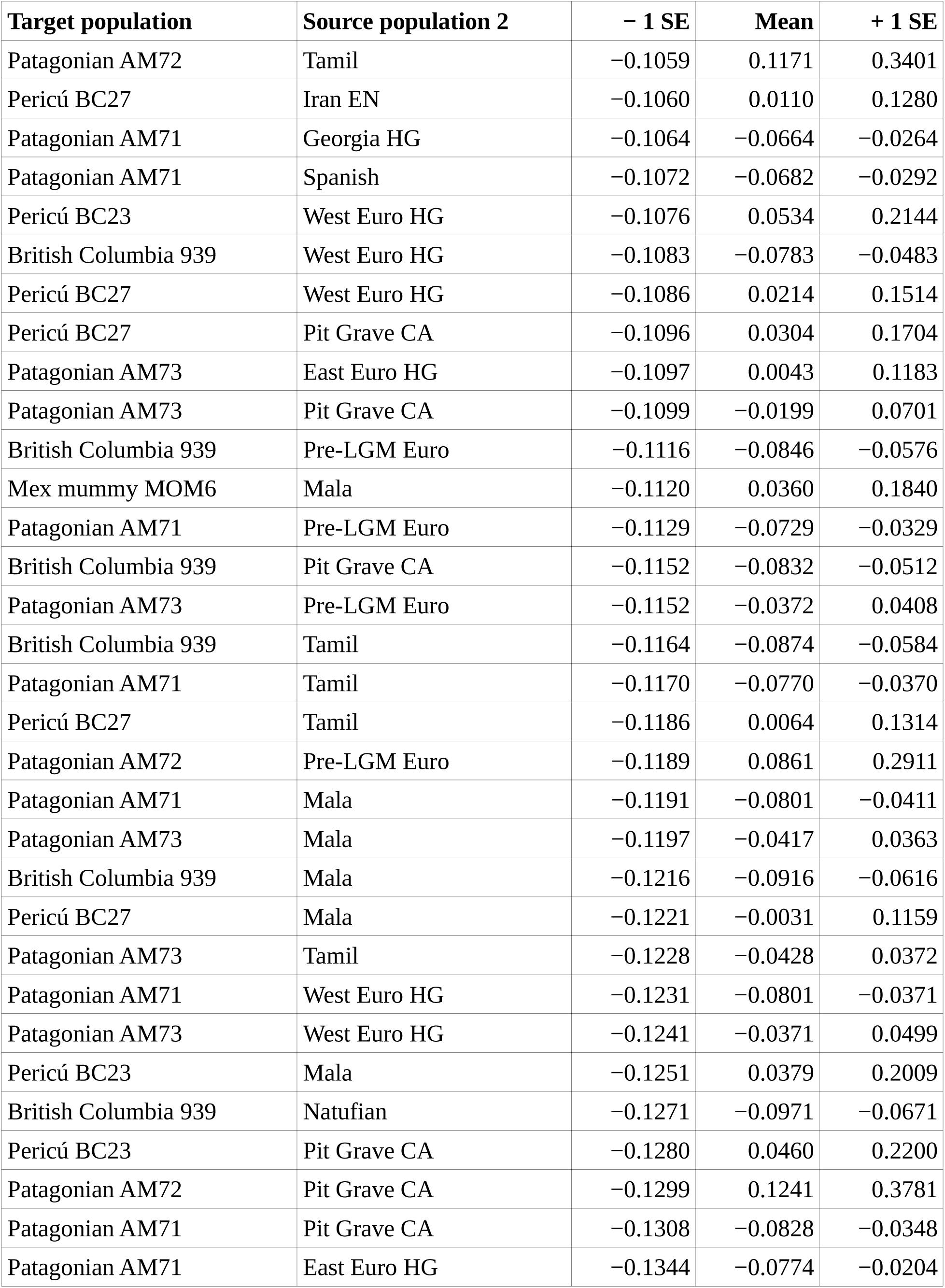
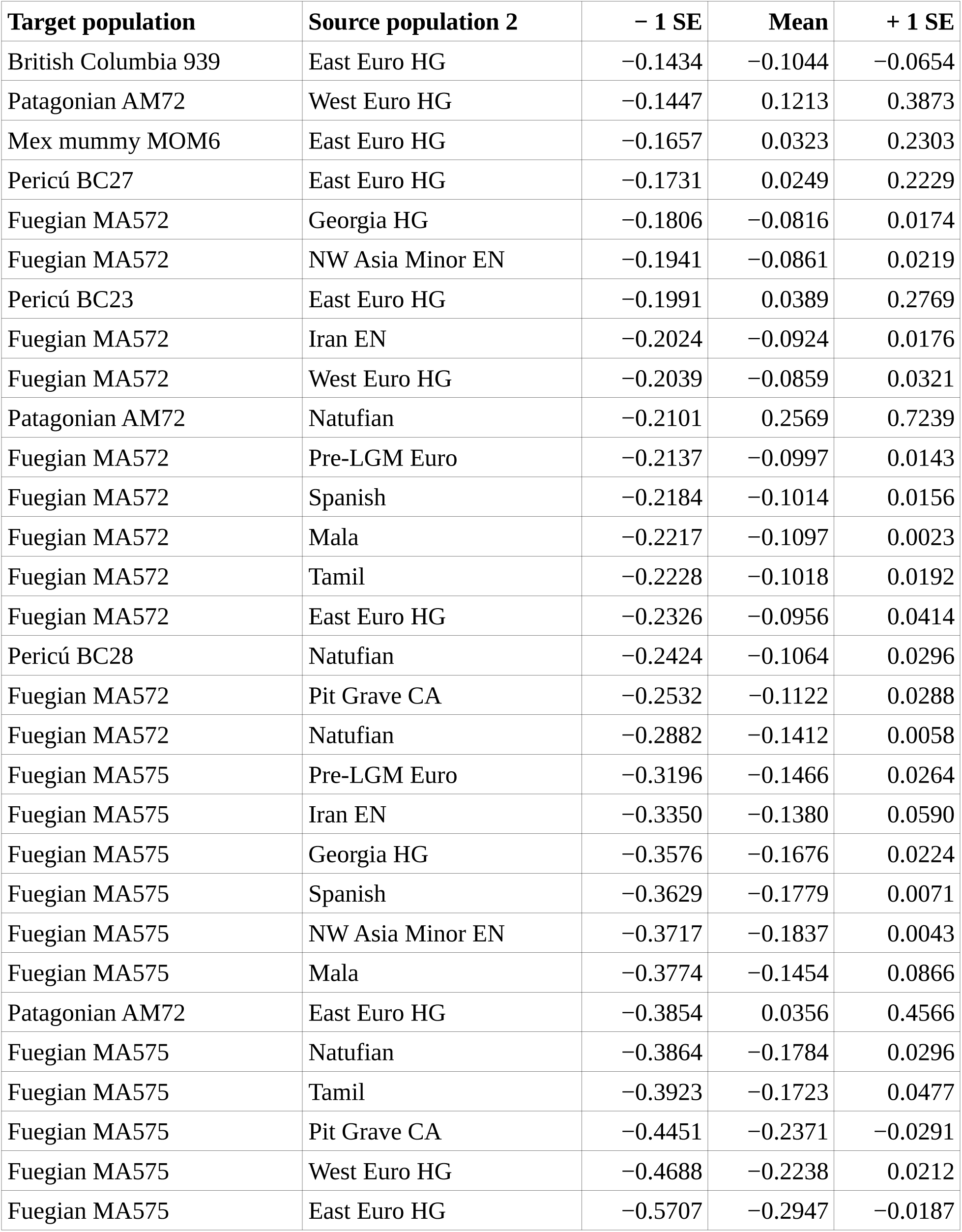
Estimated source population 2 mixture coefficients produced by 253 *qpAdm* analyses.

### Principal component analysis

A plot of the results of a principal component analysis of ancient and modern American, European, and Middle Eastern samples is shown in Figure 1. Figure 2 is a plot of the results of a second principal component analysis, in which the Middle Eastern samples were excluded, leaving only the American and European samples.

**Figure 1:**
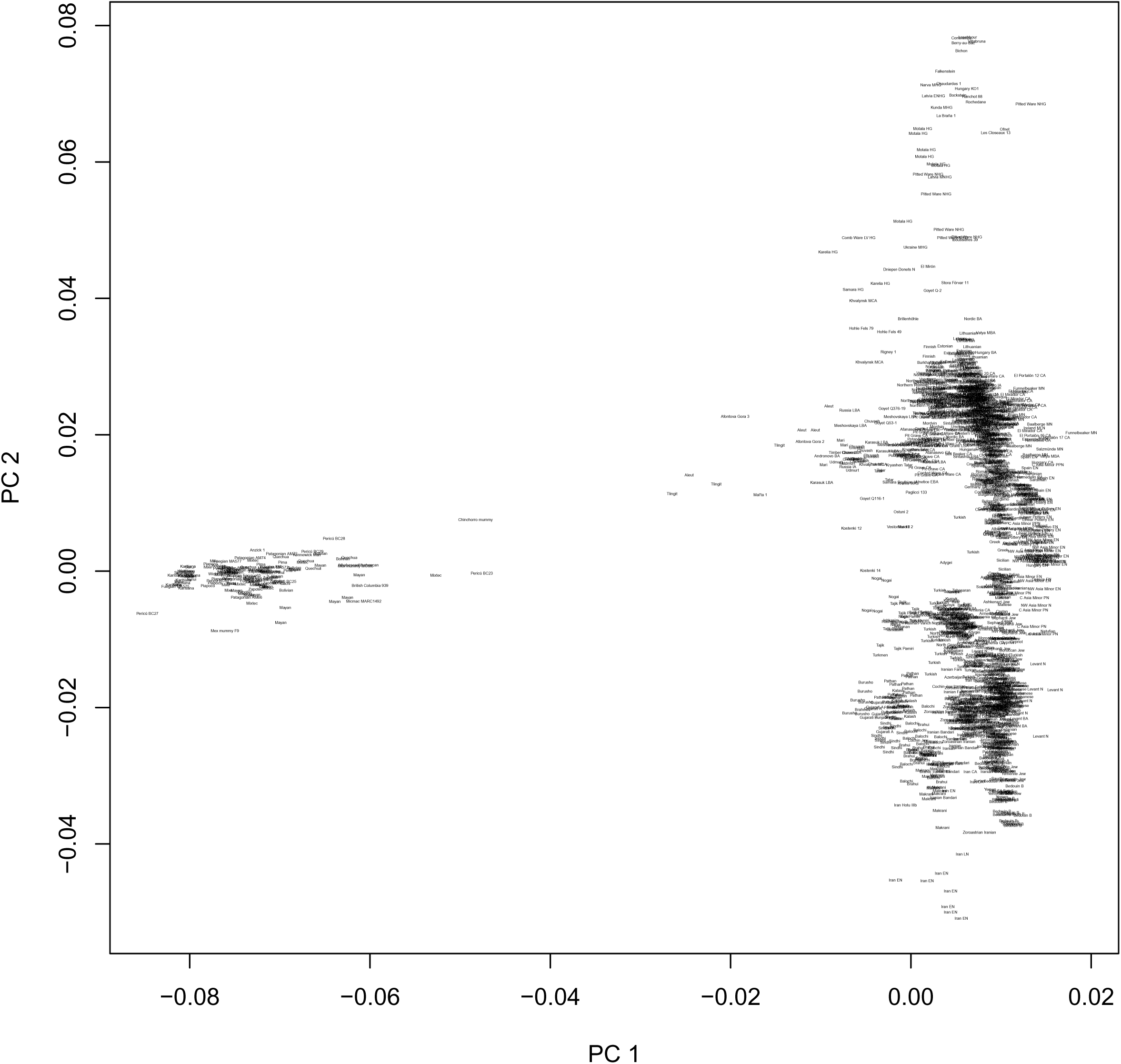
**PCA of Amerindians, Europeans, and Middle Easterners**

**Figure 2:**
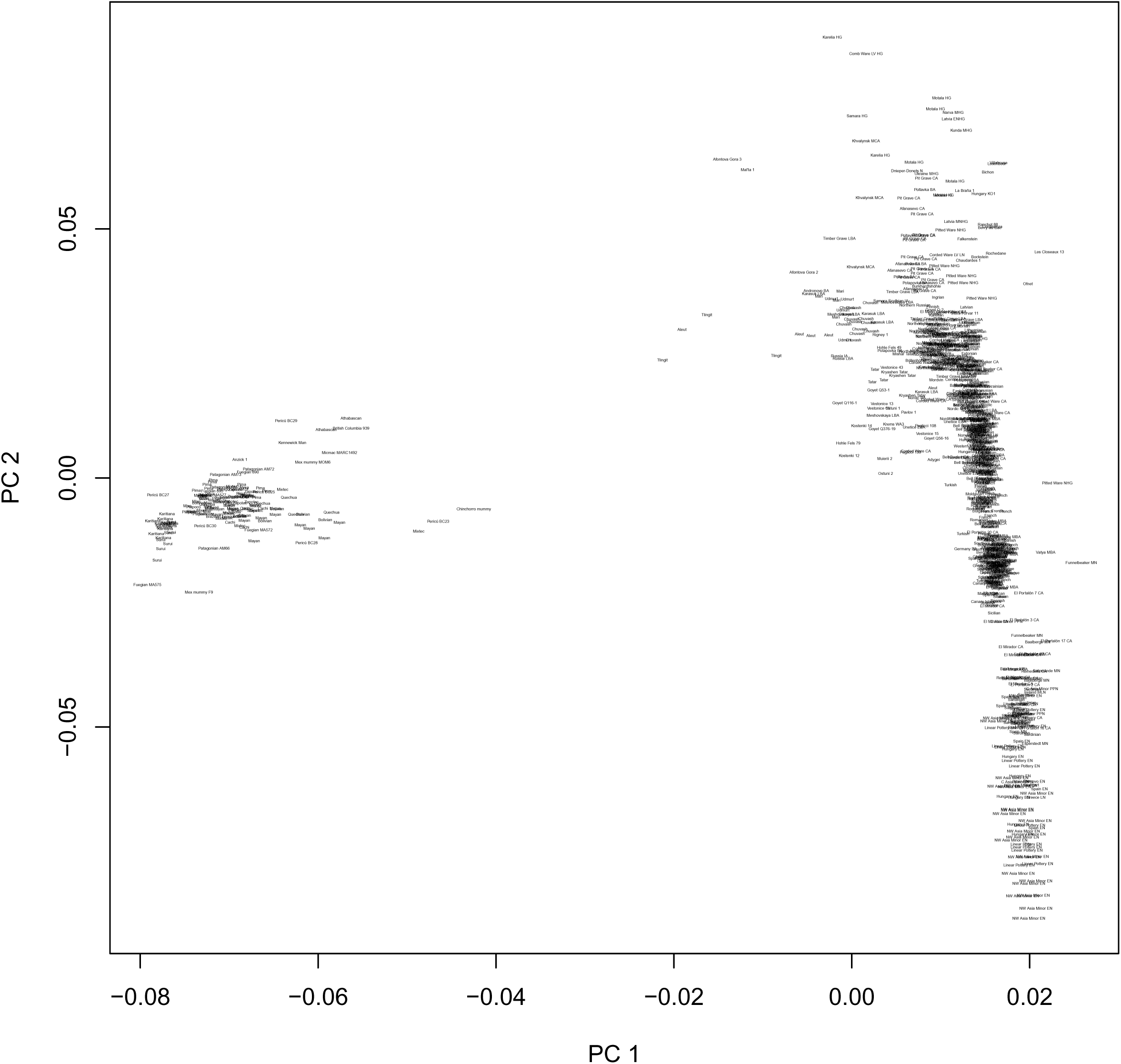
**PCA of Amerindians and Europeans**

### ADMIXTURE analysis

Plots of the results of *K* = 4, 5, 6, 7, 9, and 13 ADMIXTURE analyses of the ancient American samples, along with other ancient and modern samples, are shown in Figures 3 through 8.

**Figure 3:**
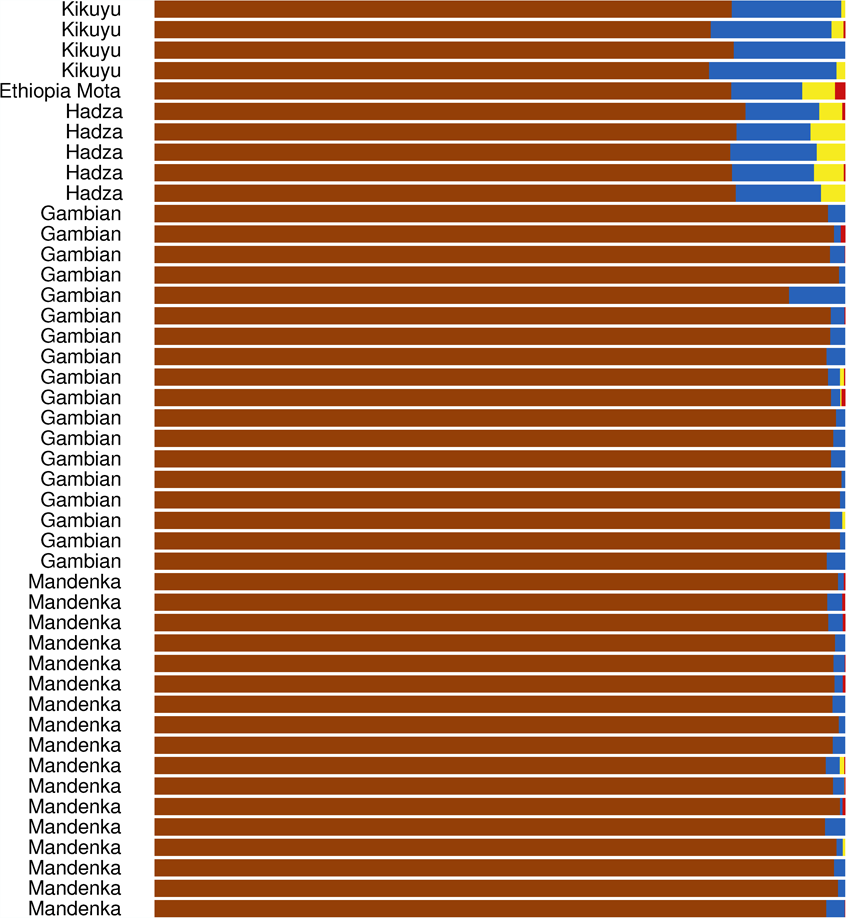
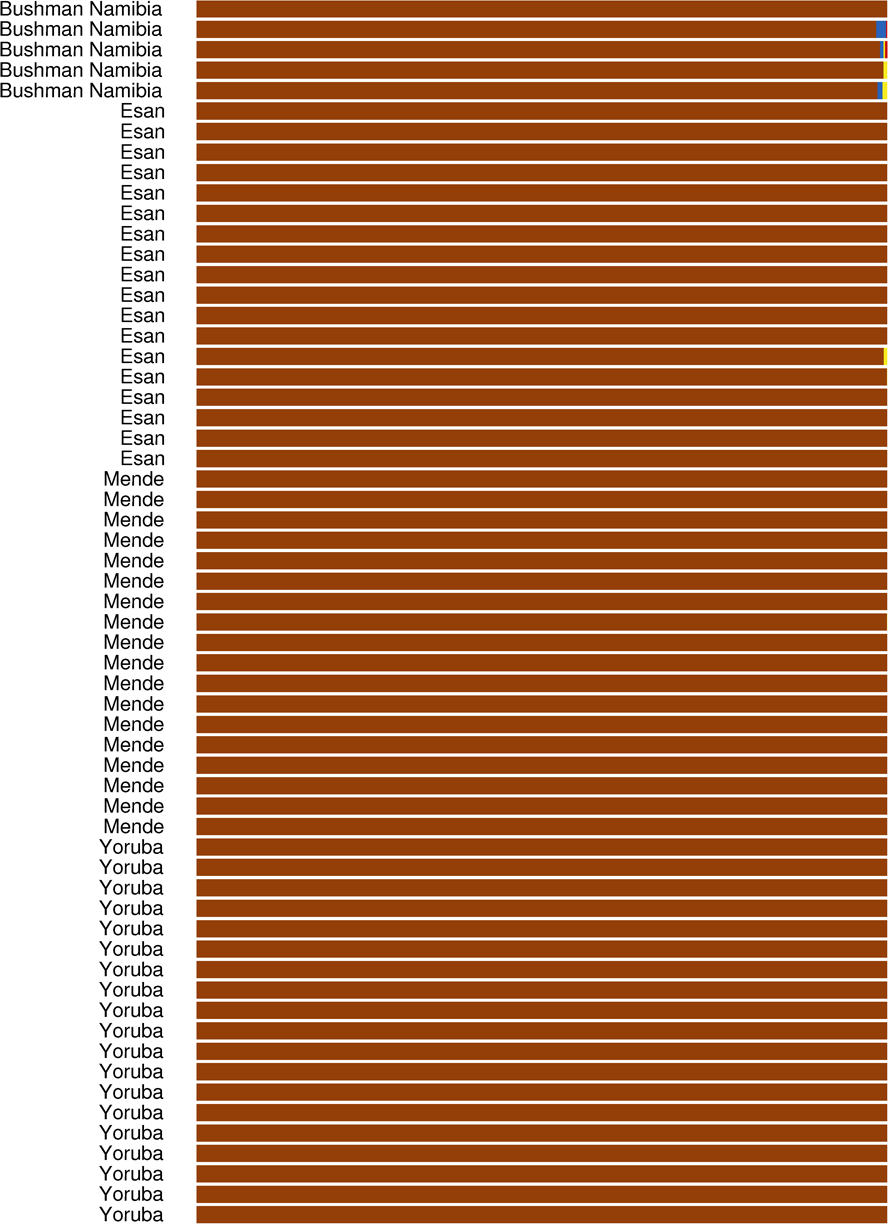
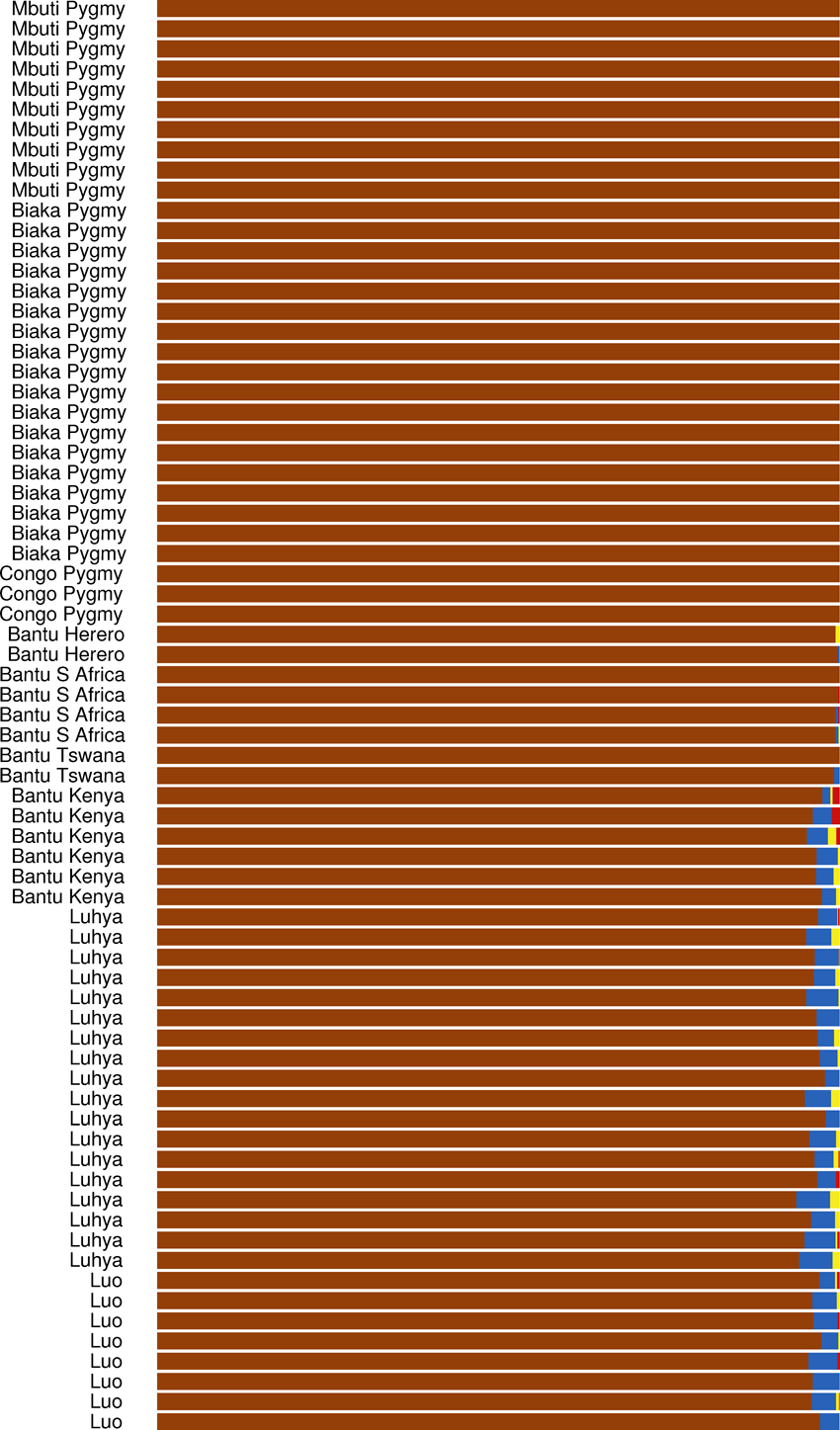
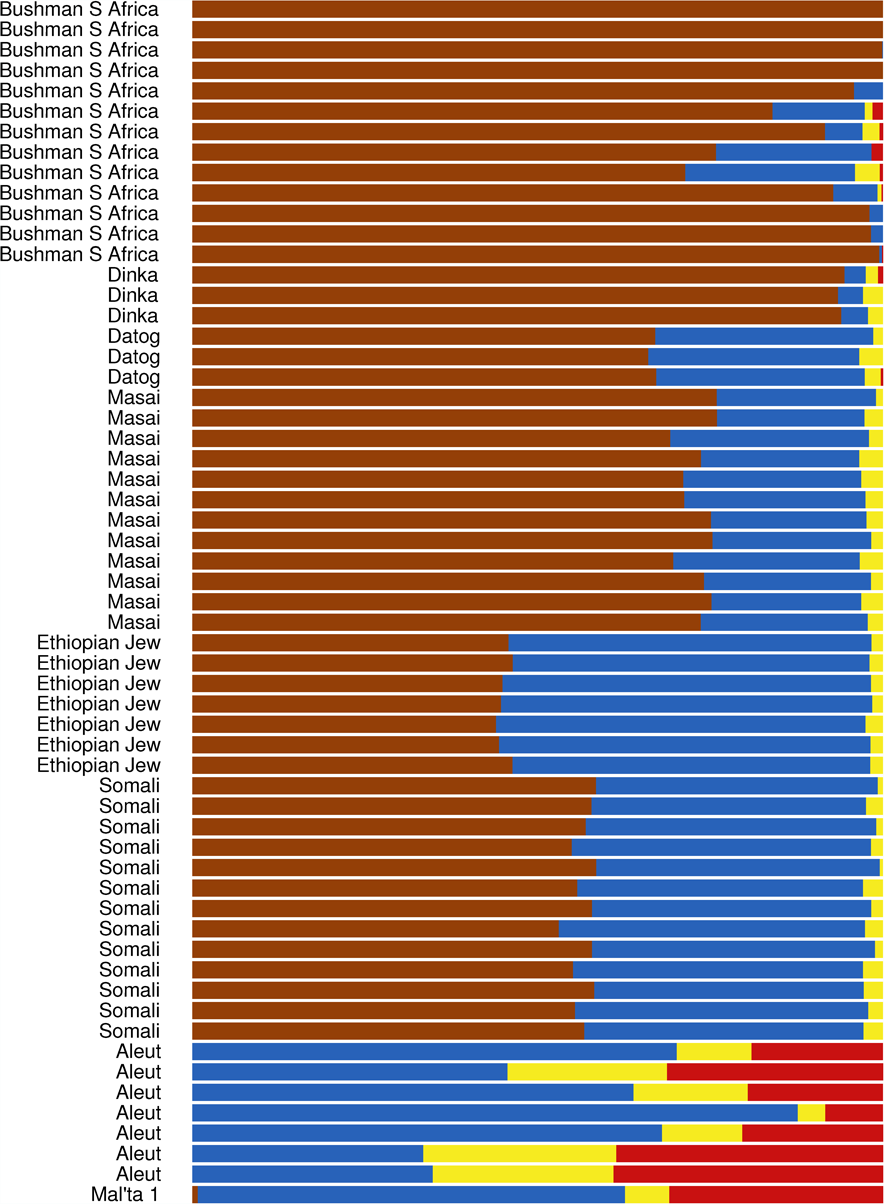
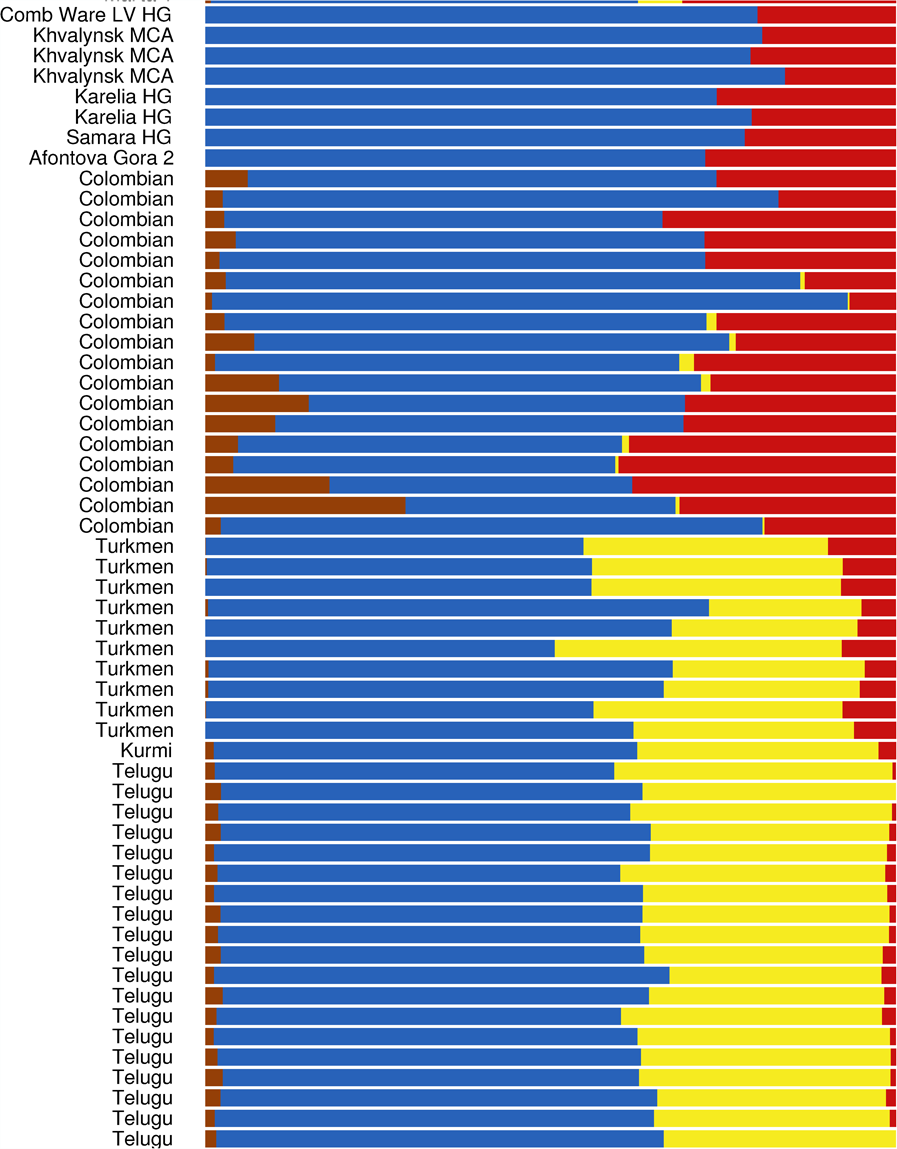
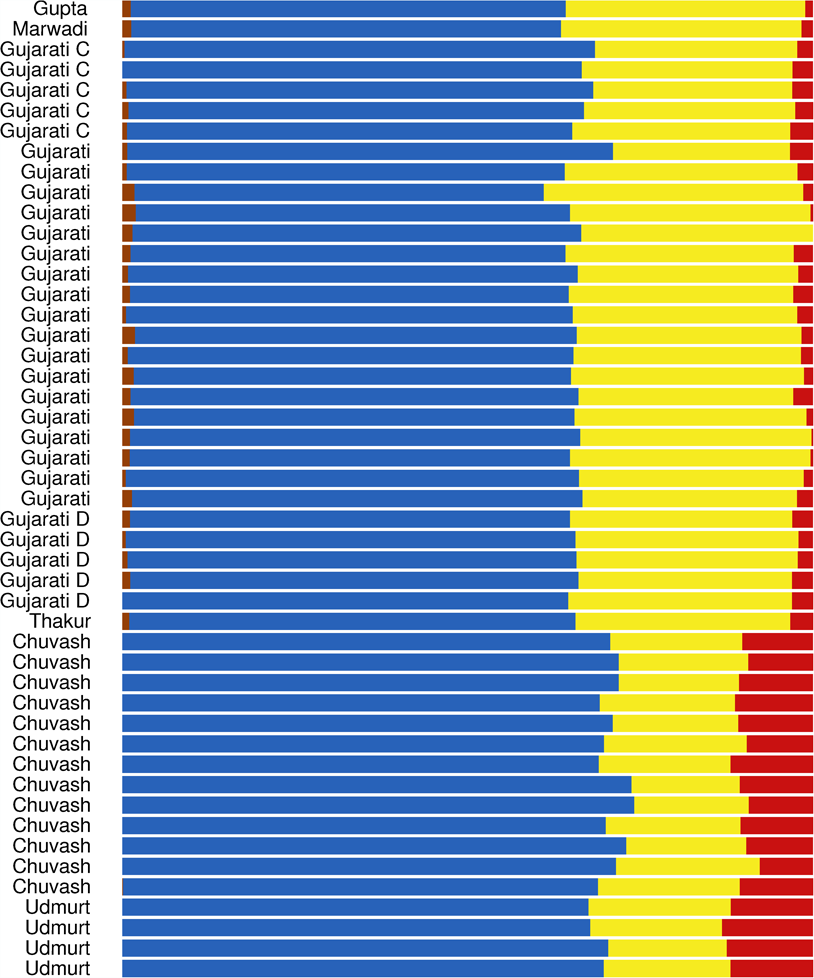
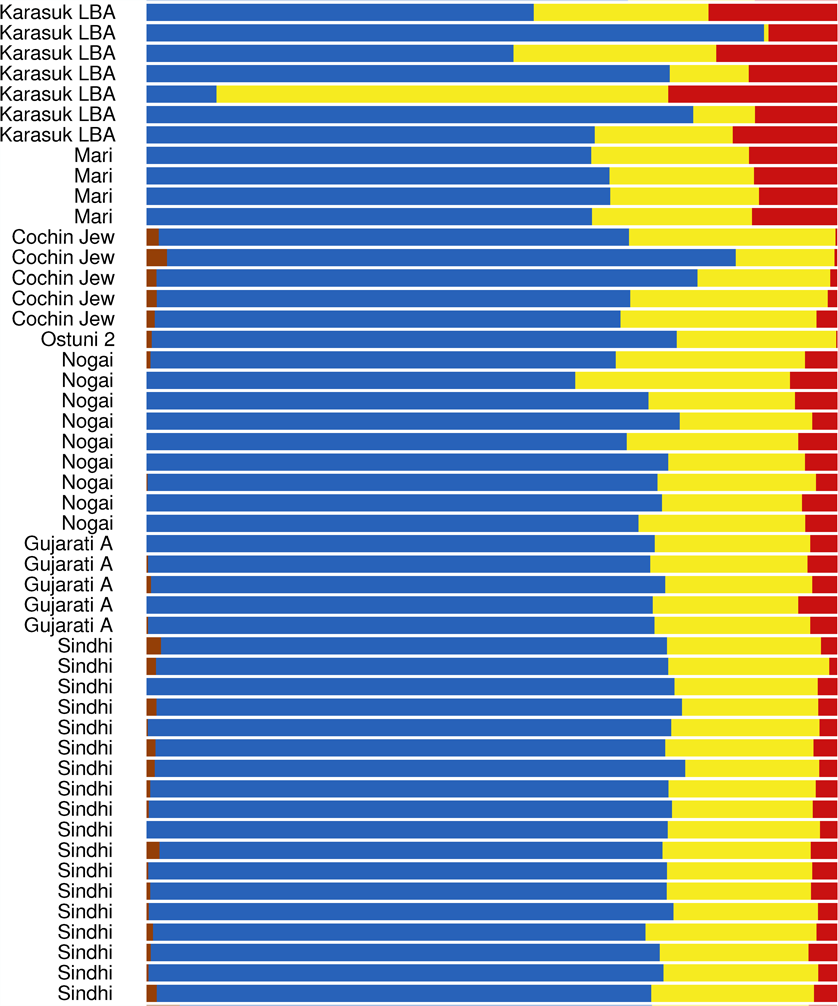
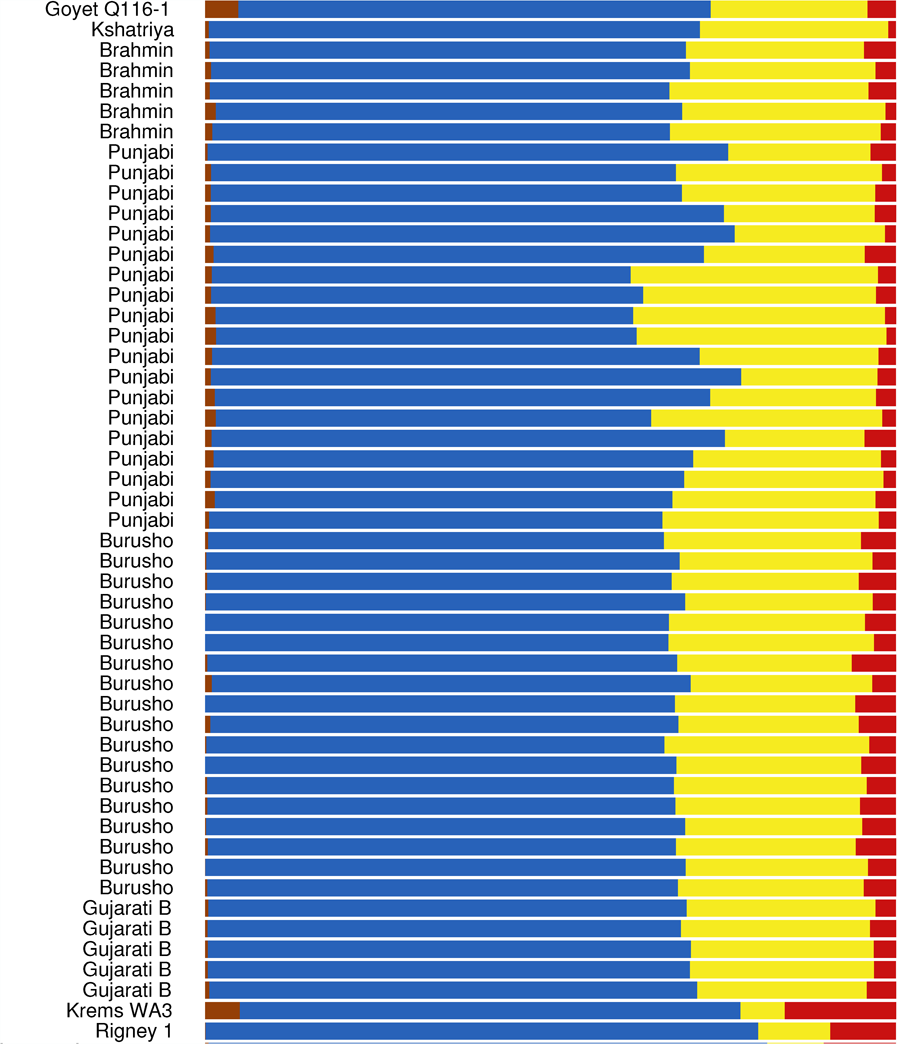
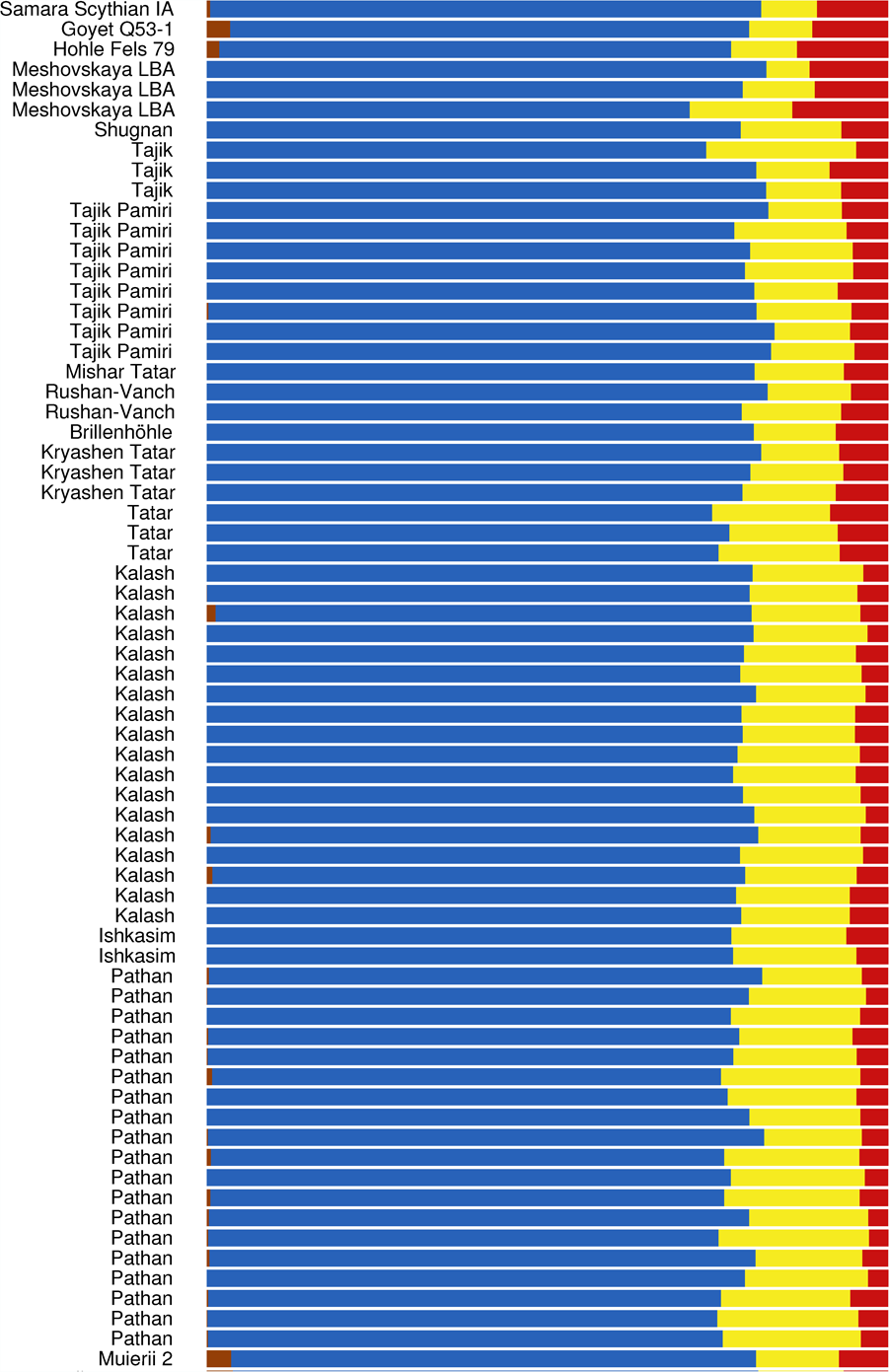
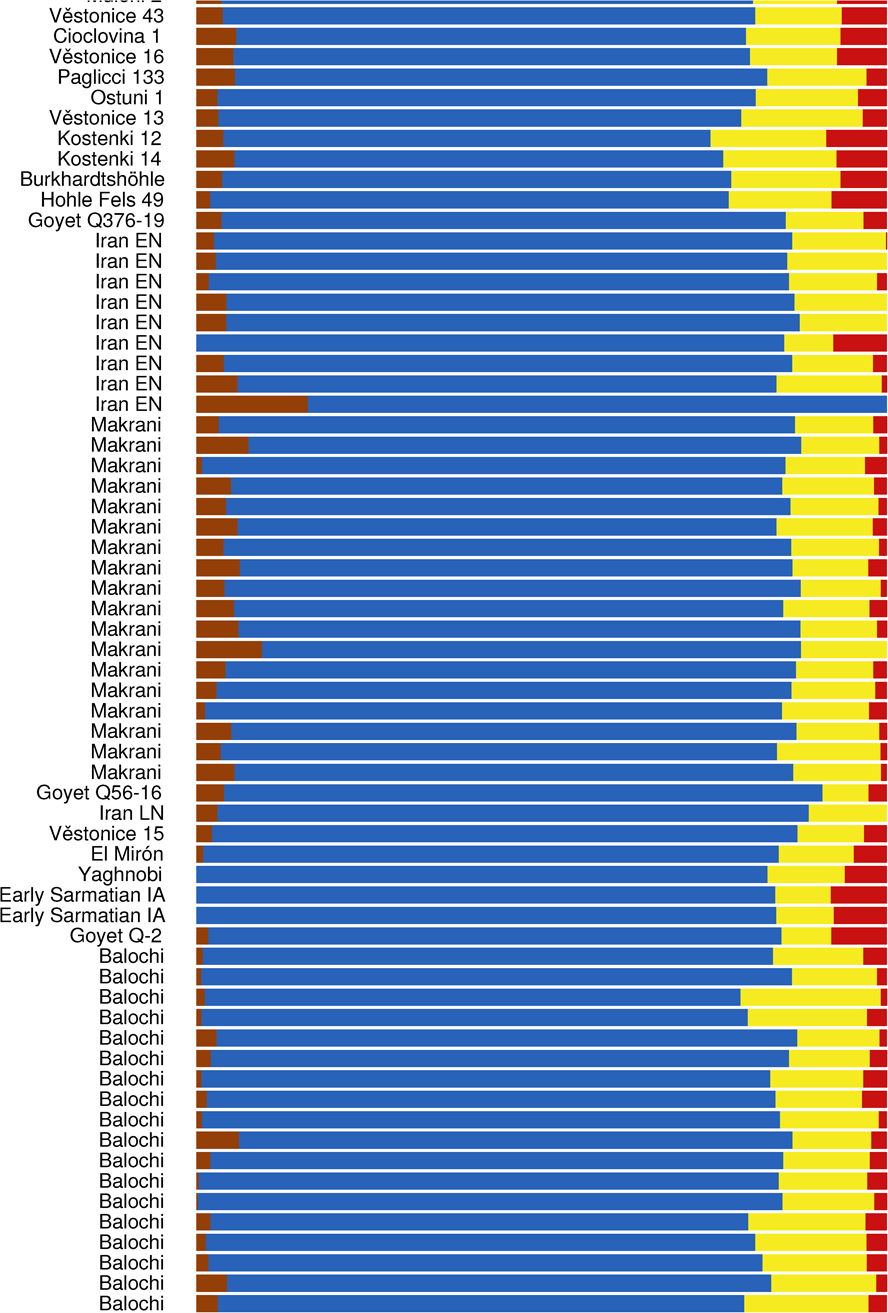
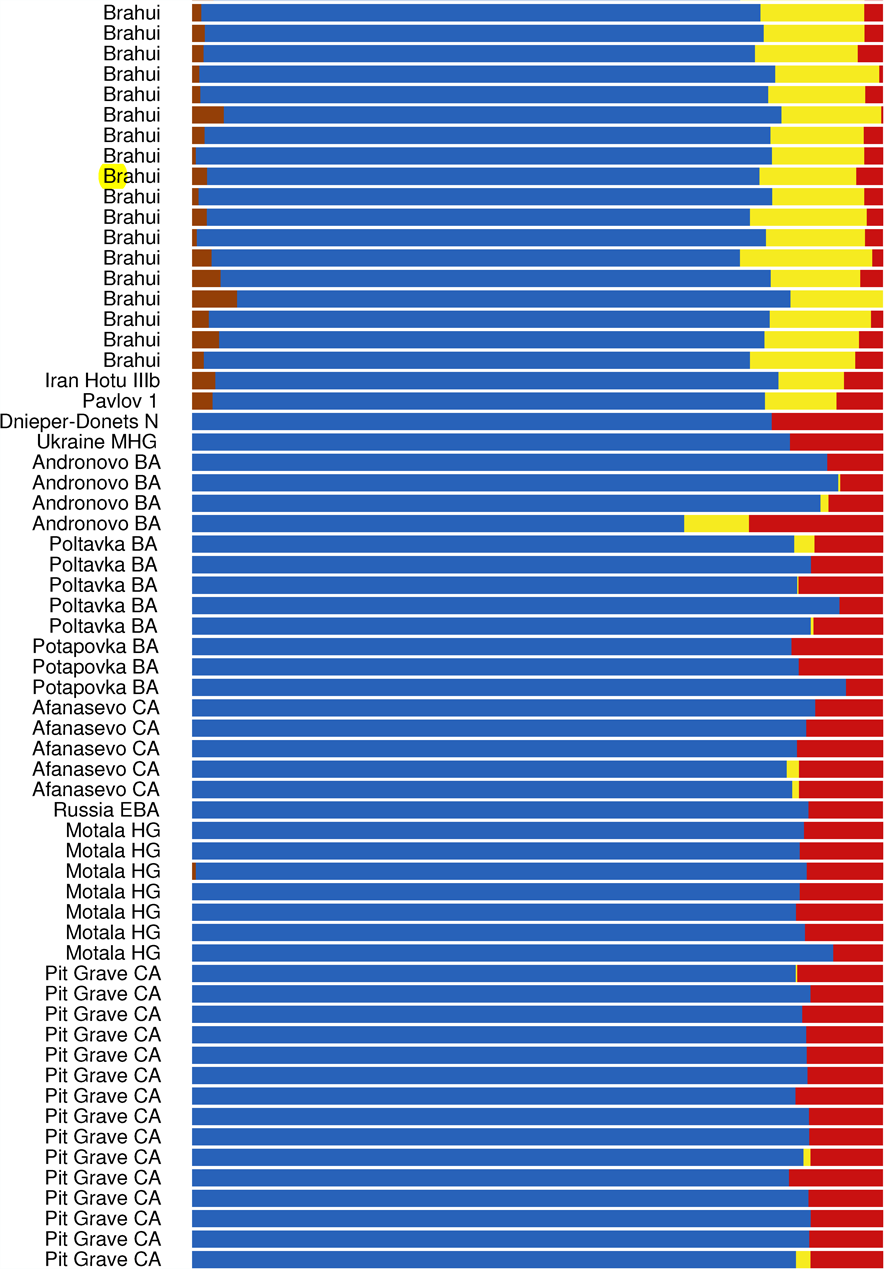
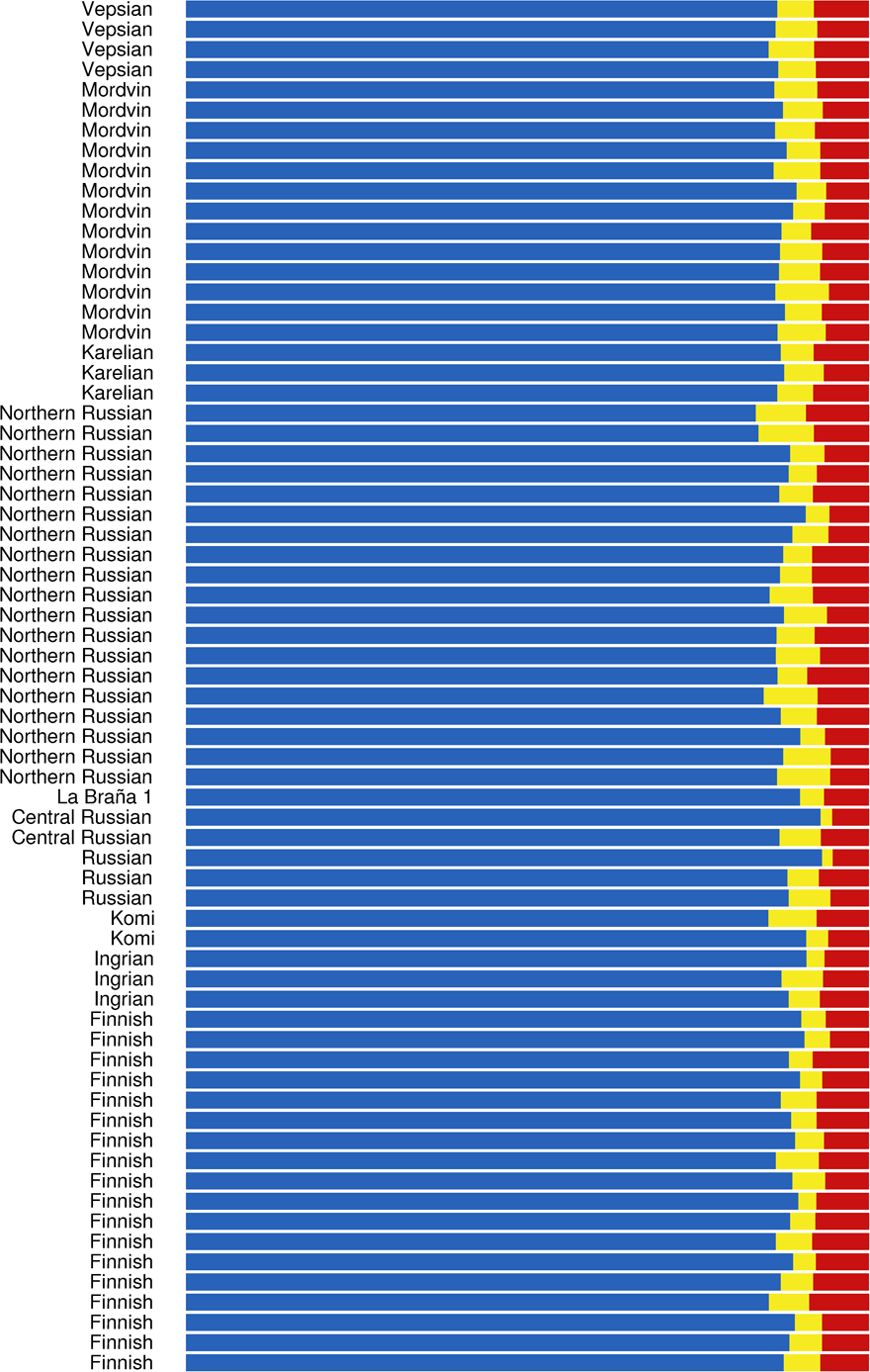
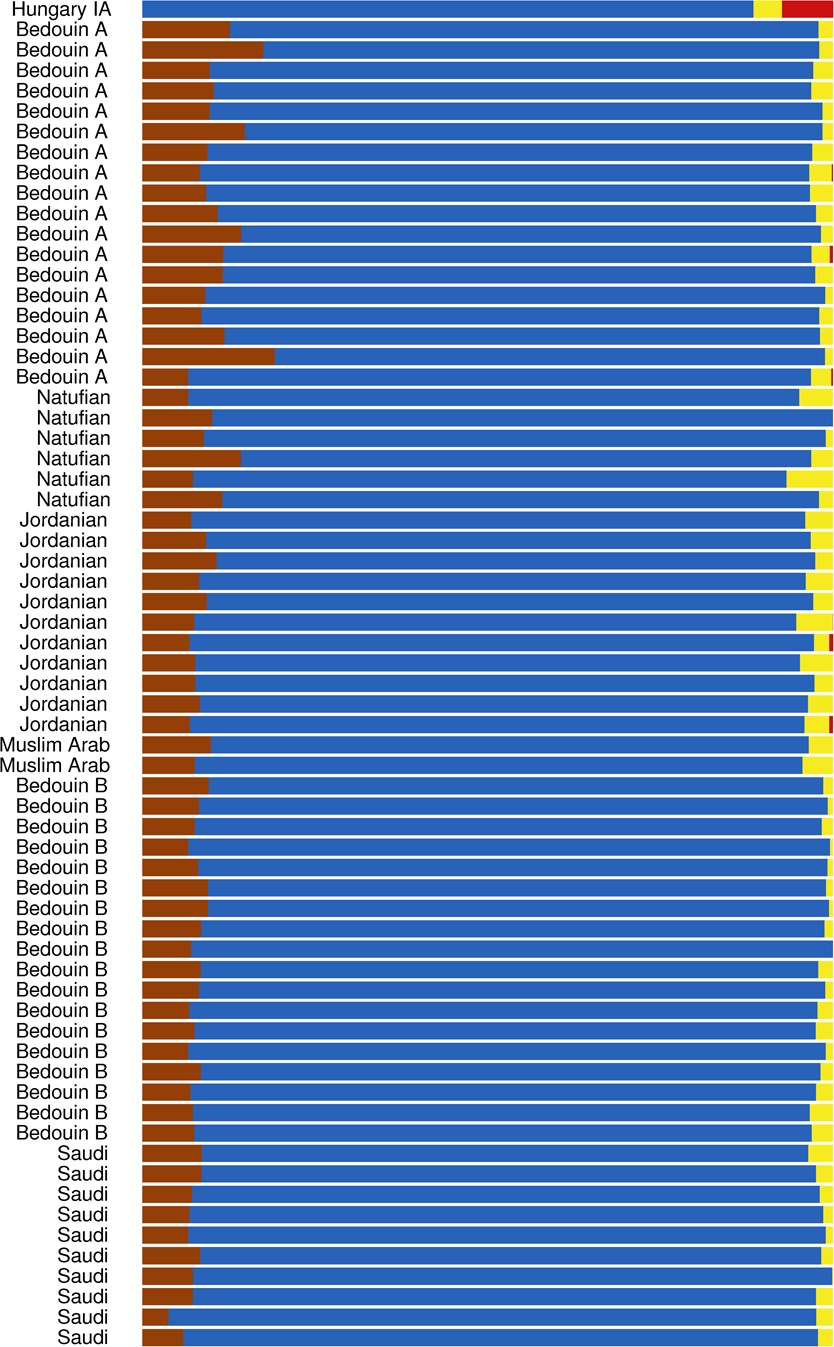
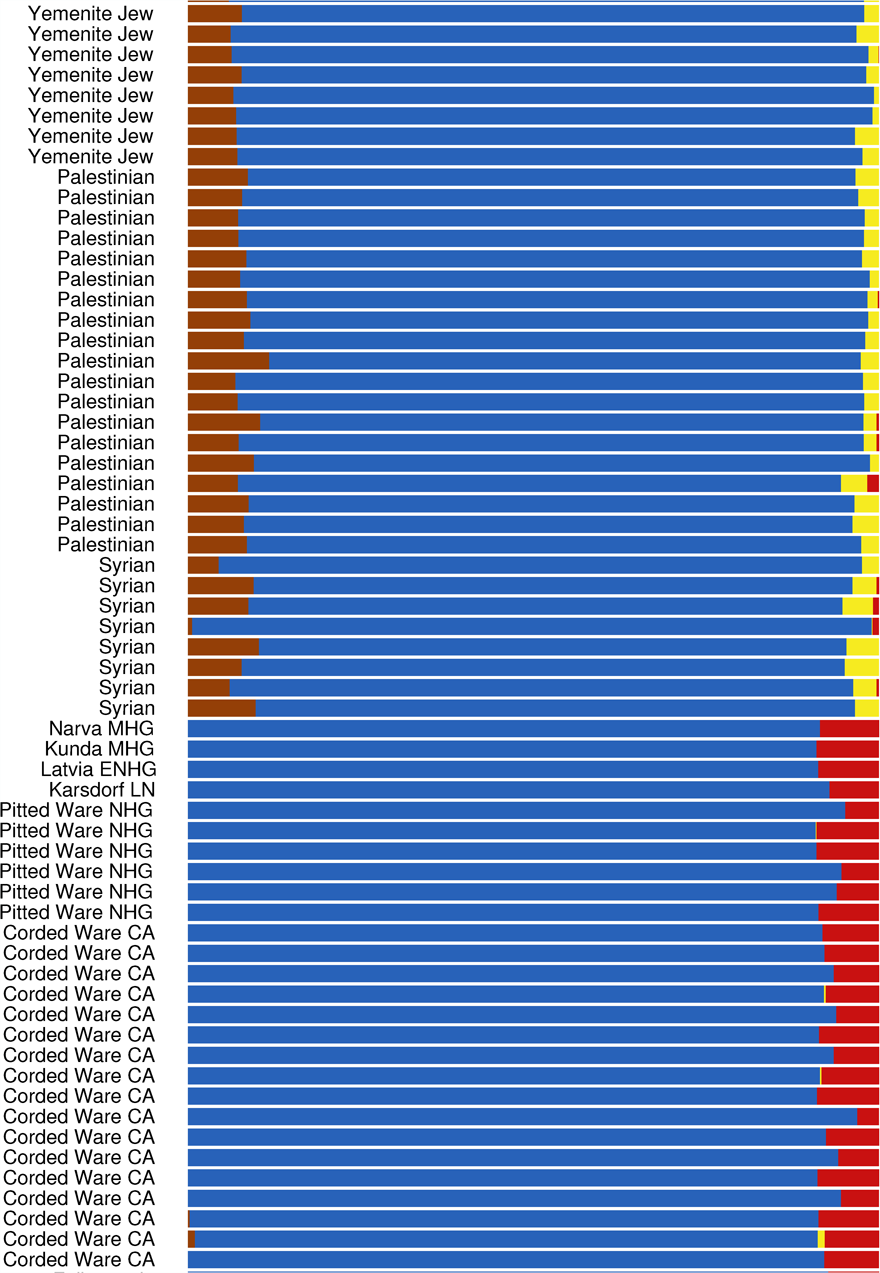
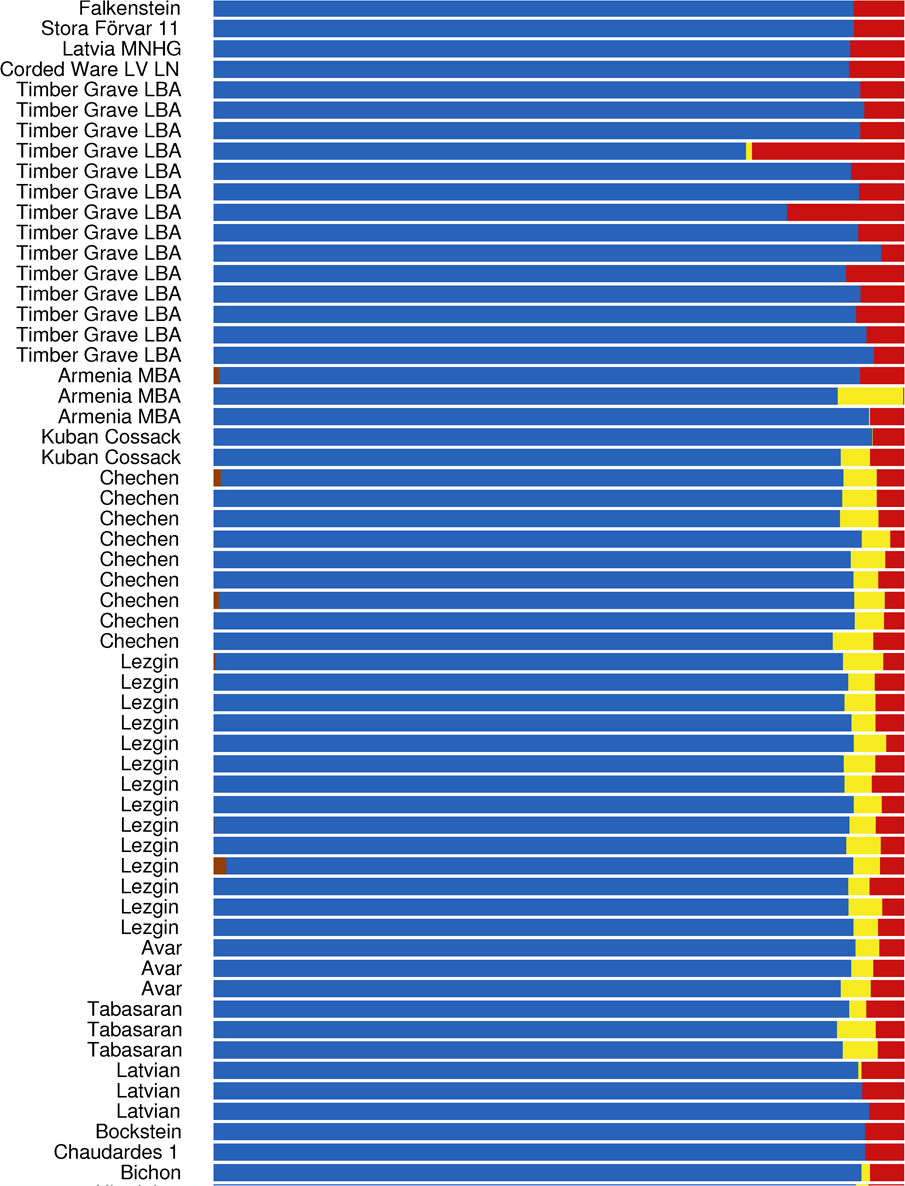
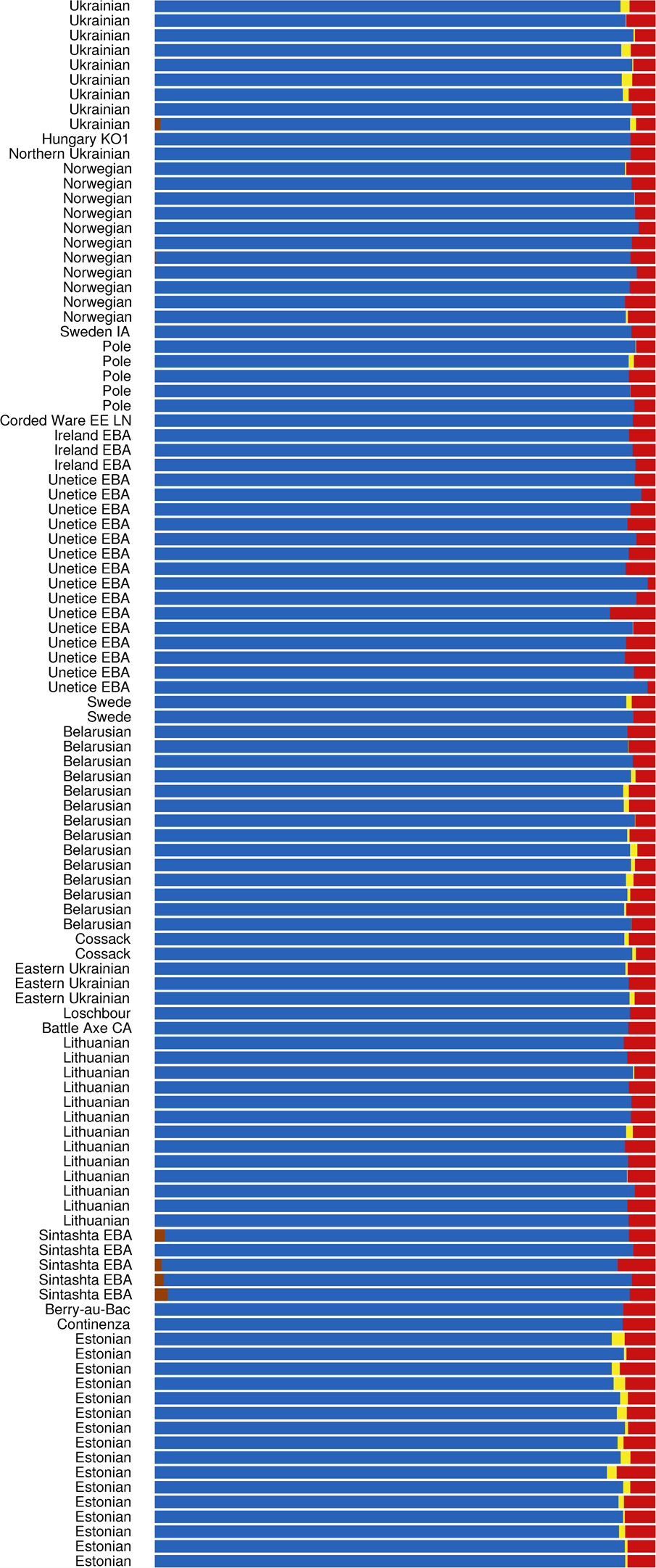
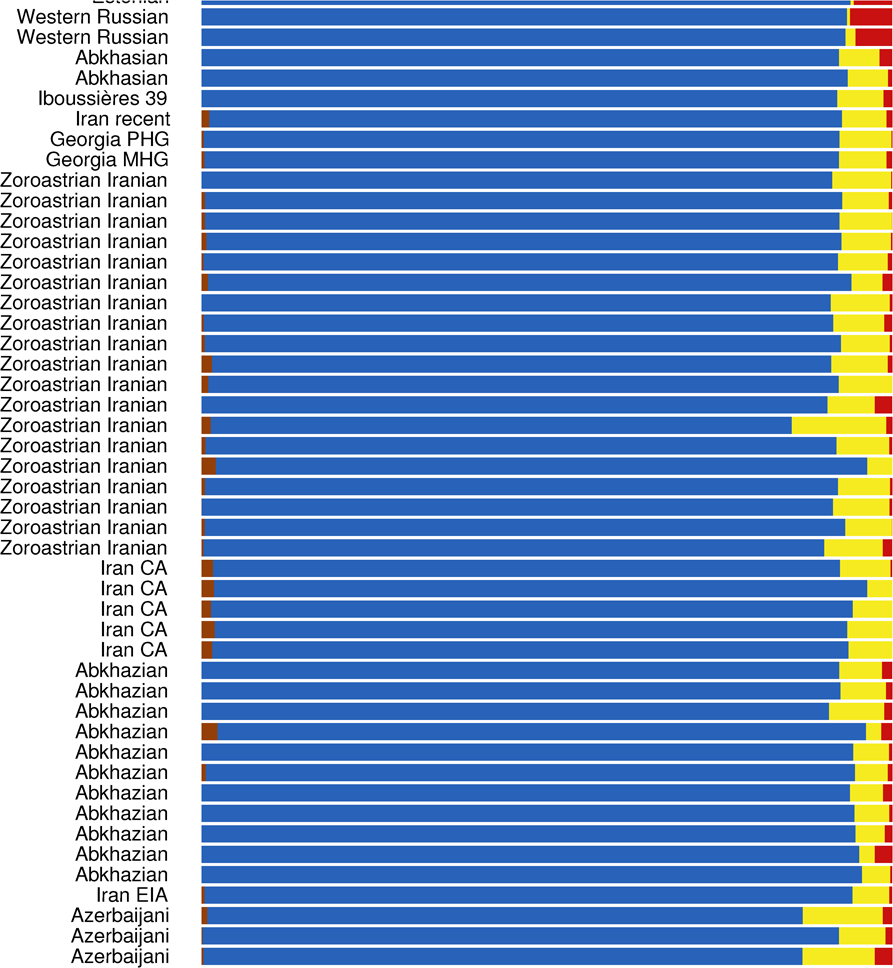
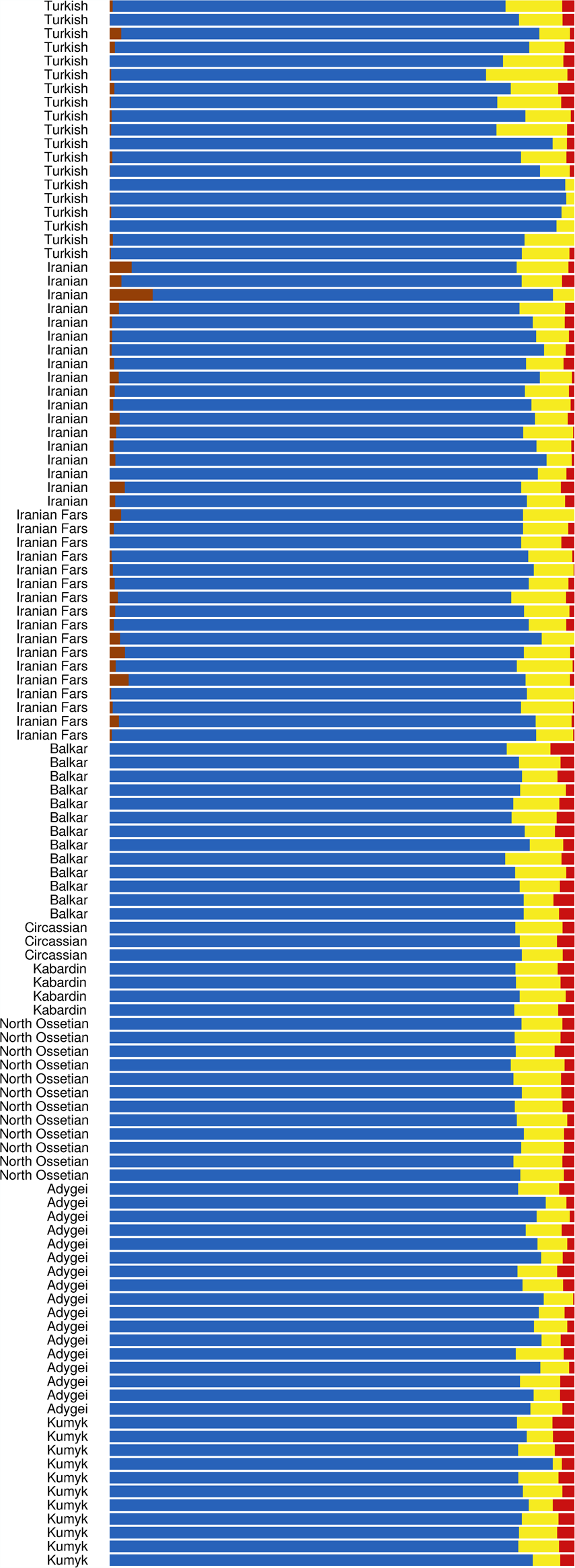
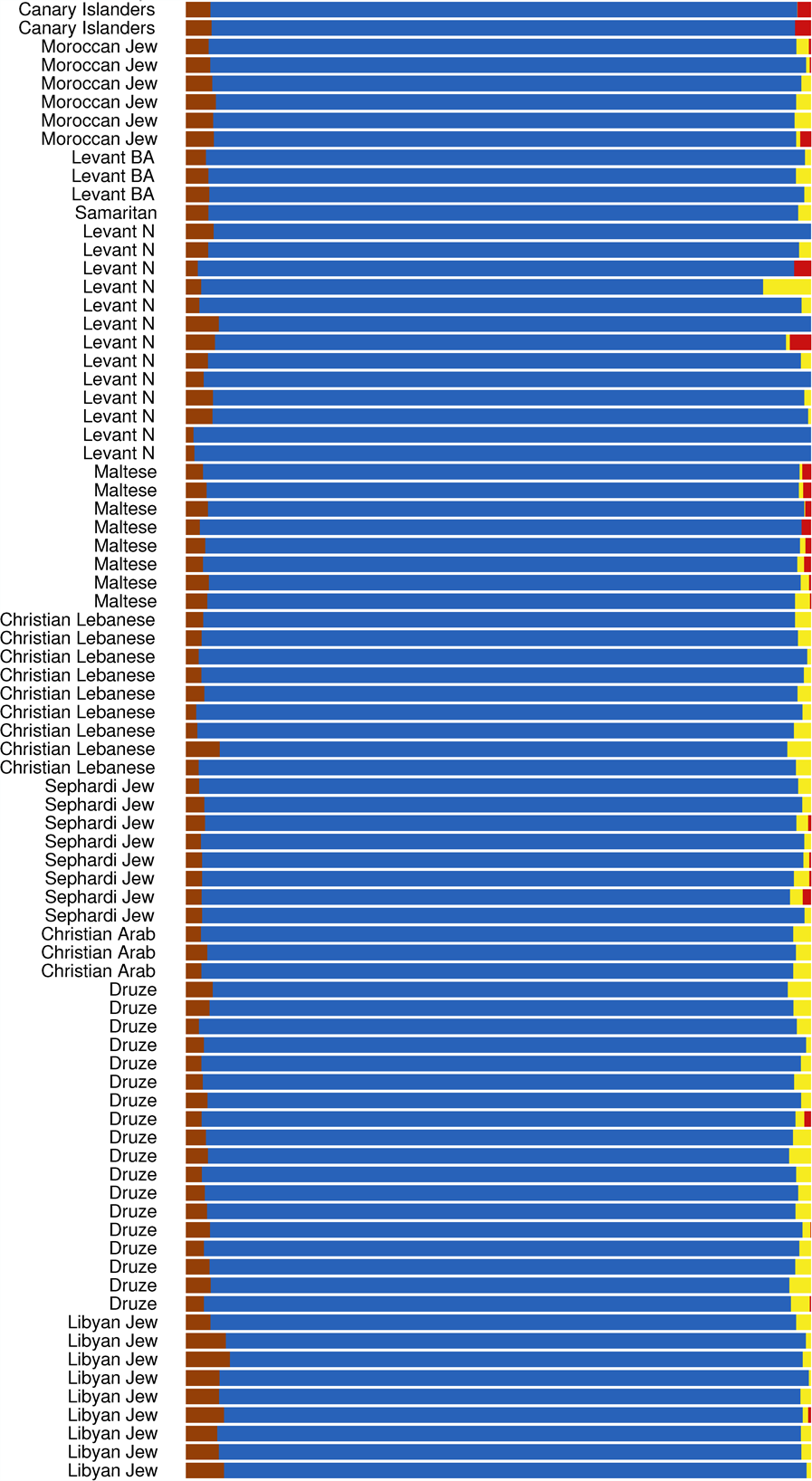
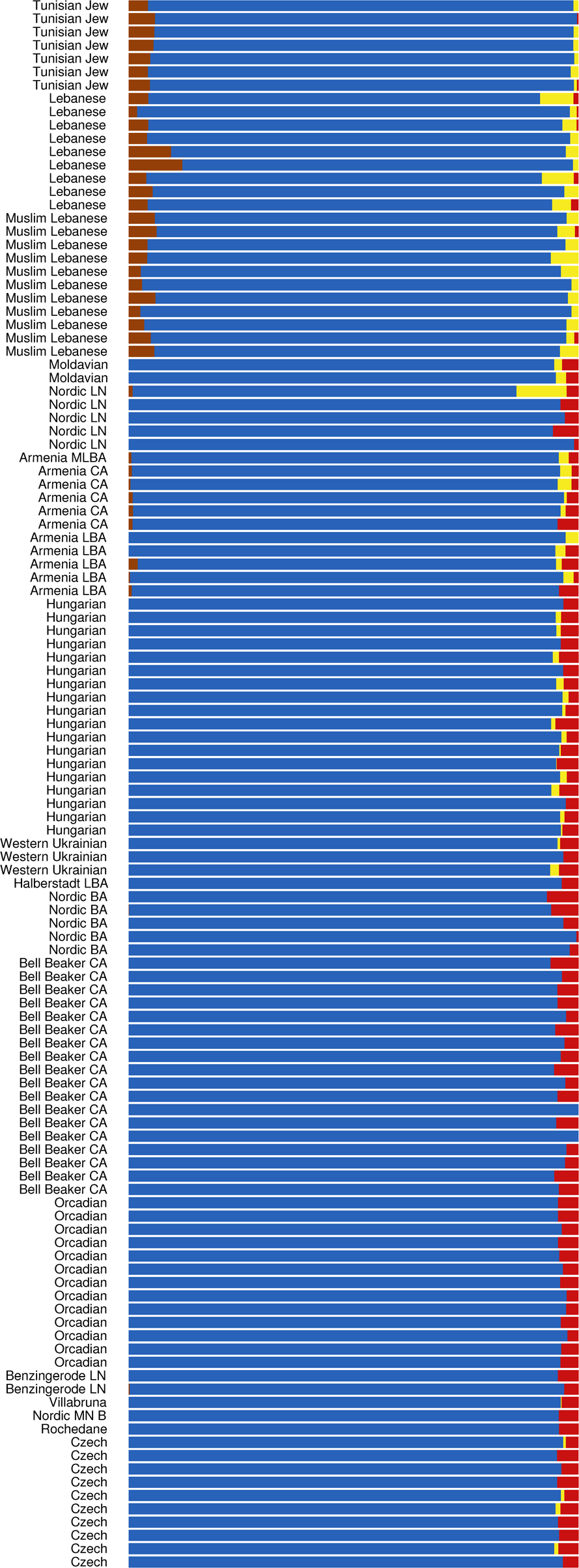
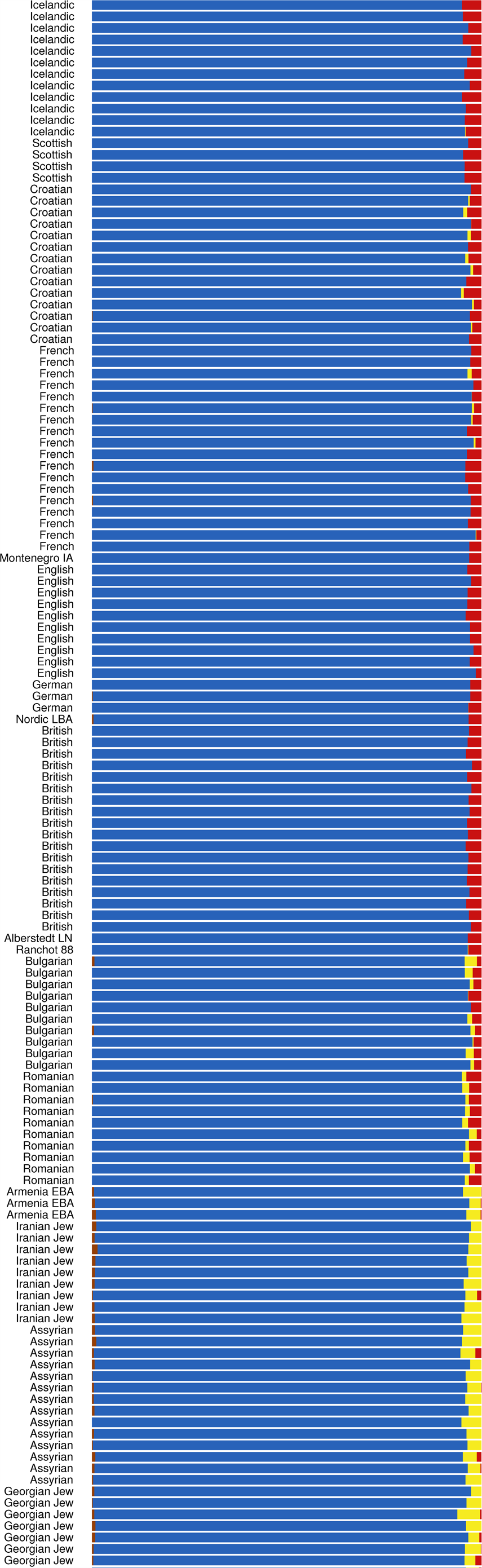
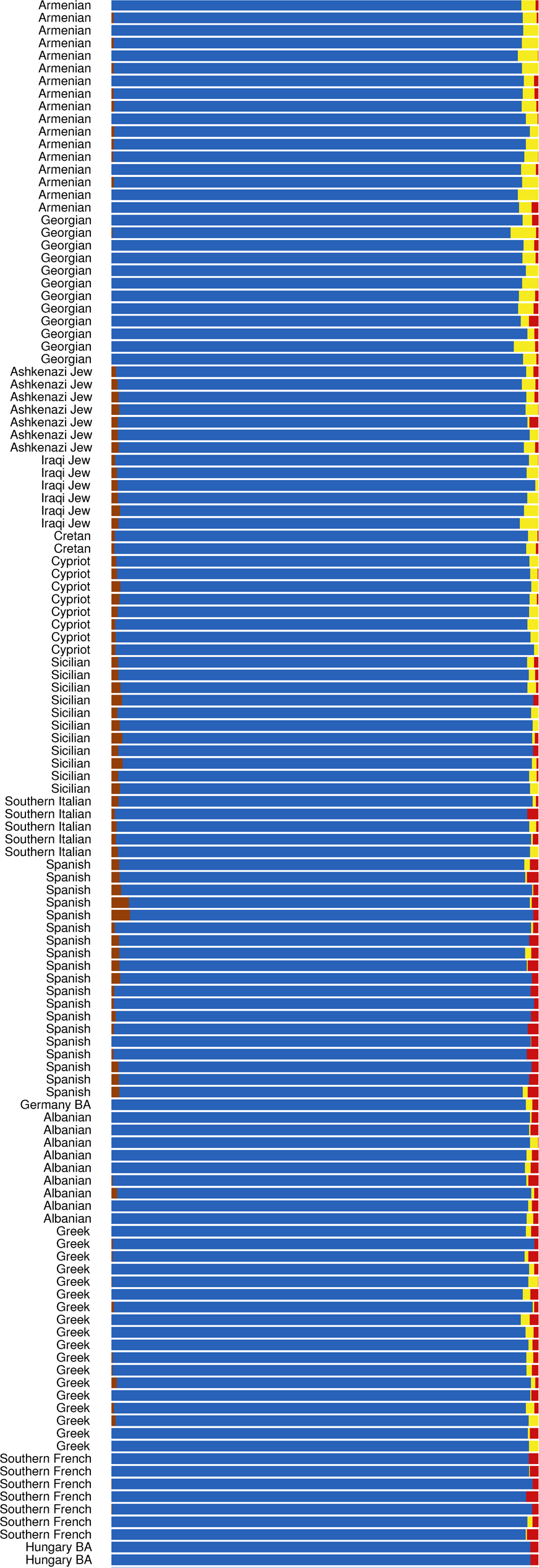
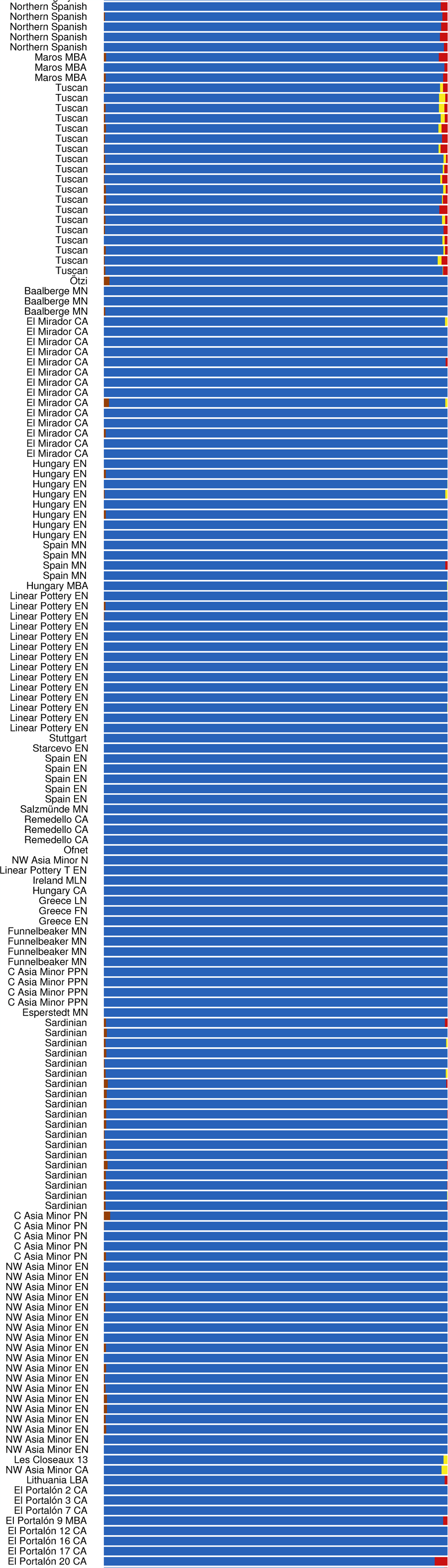
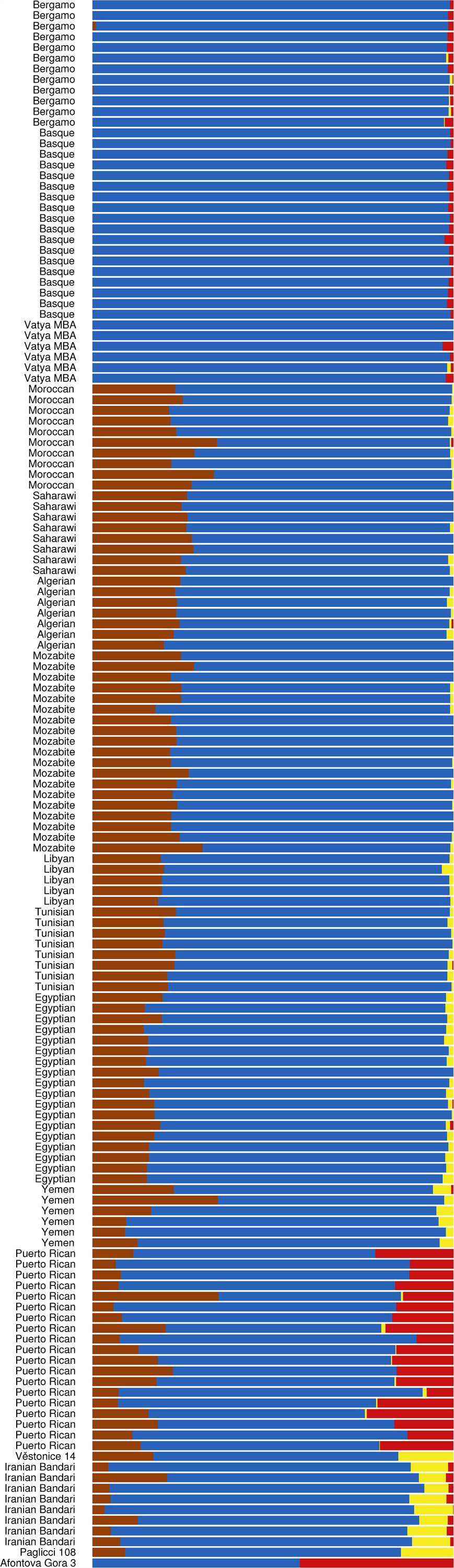
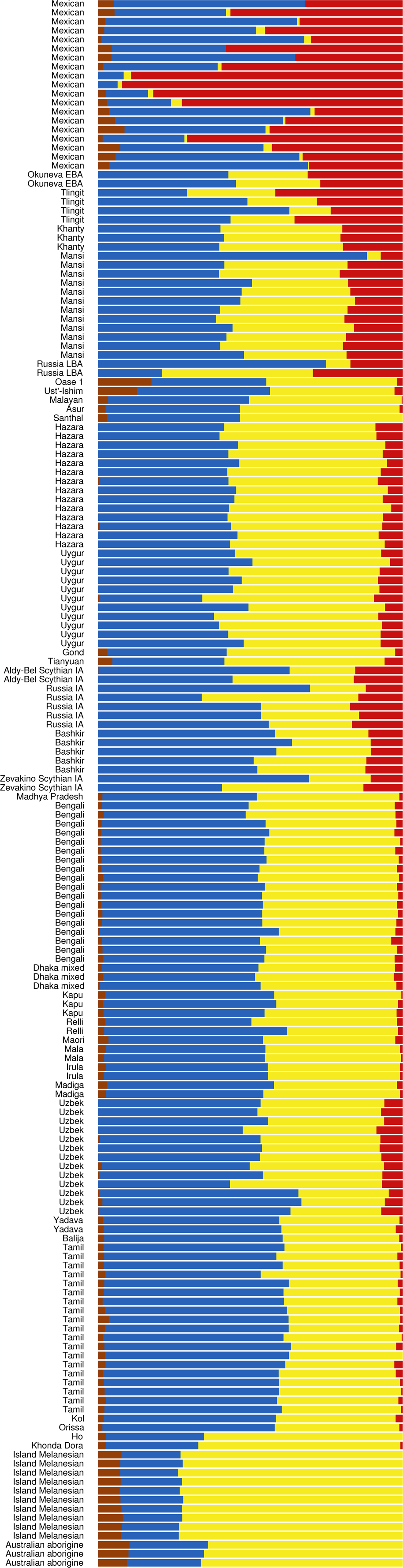
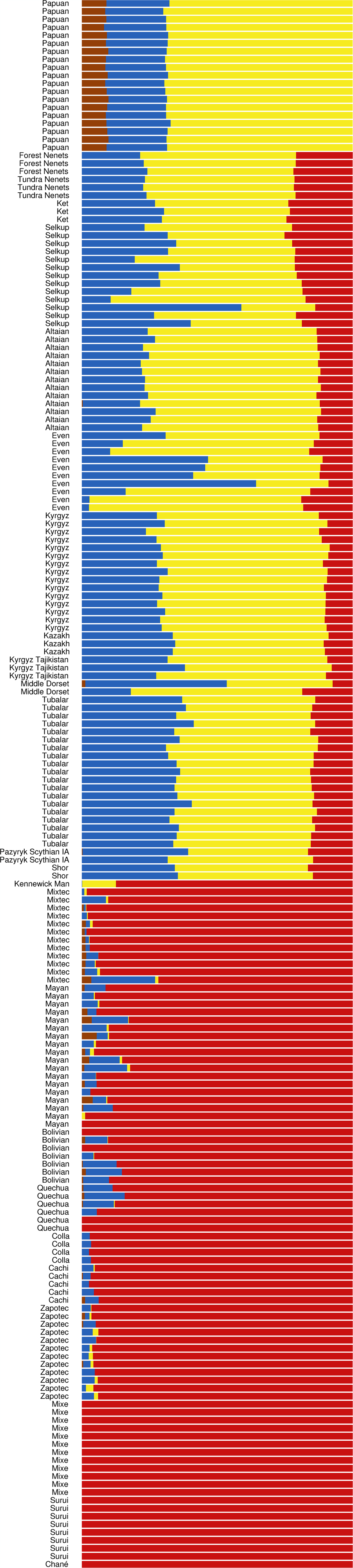
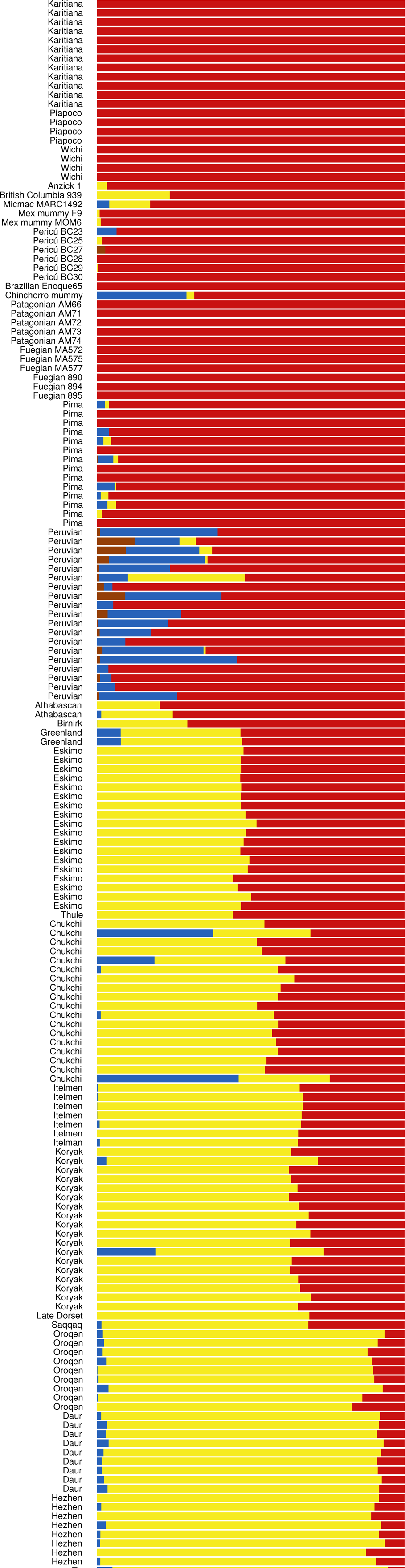
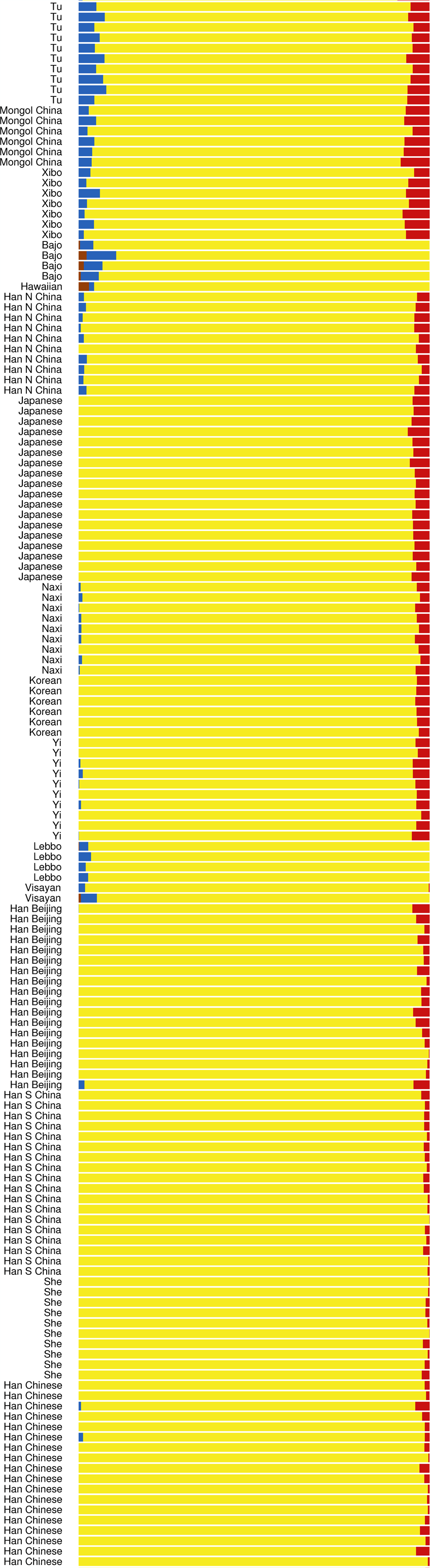
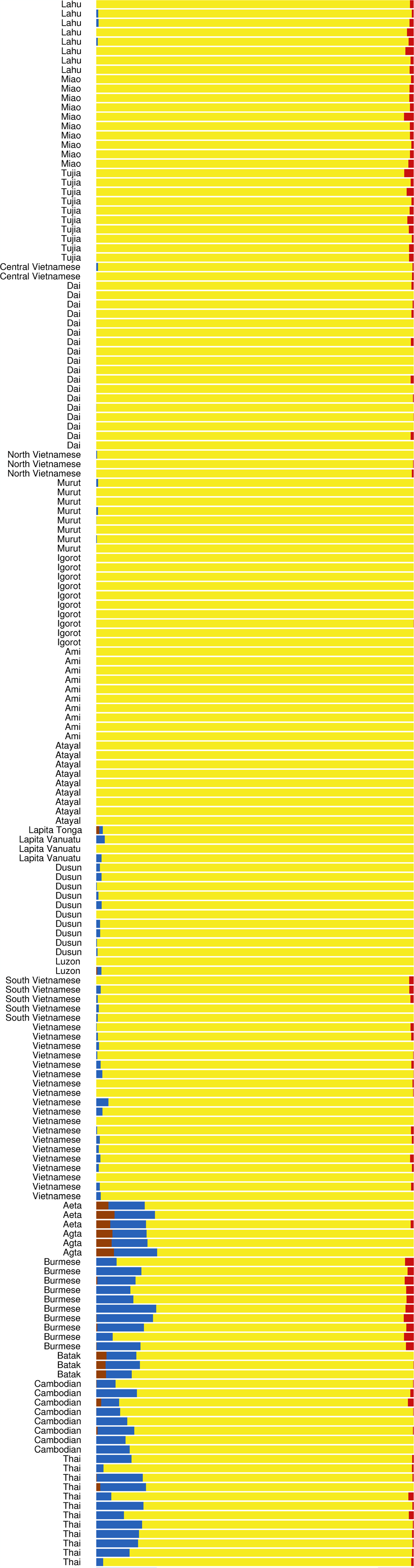
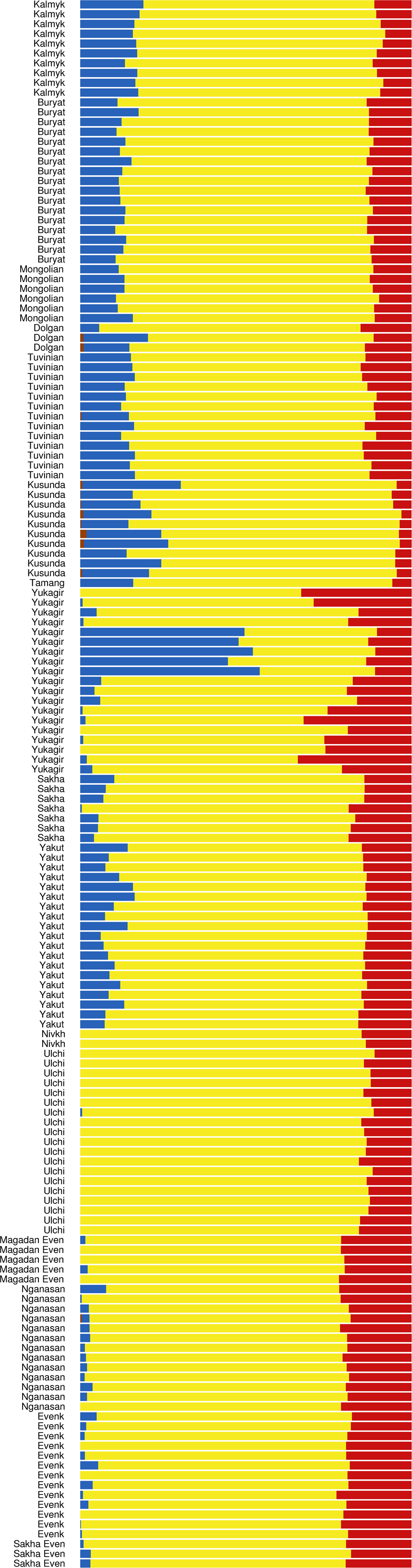
**K = 4 ADMIXTURE analysis**

**Figure 4:**
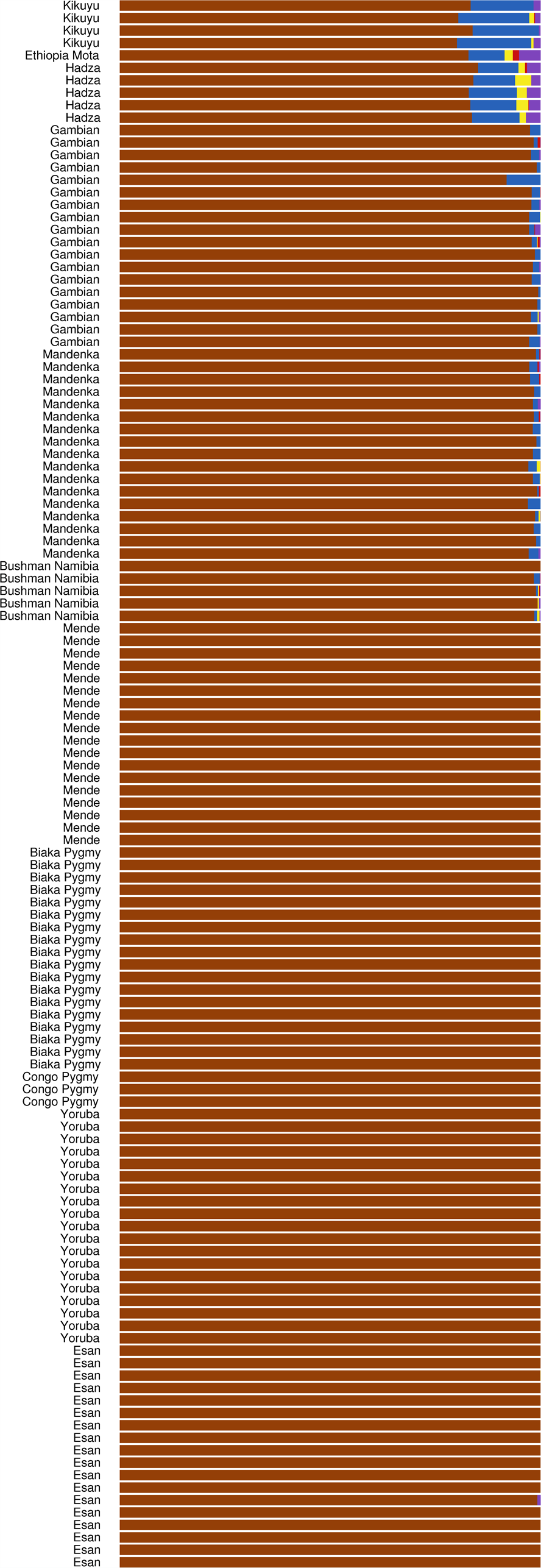
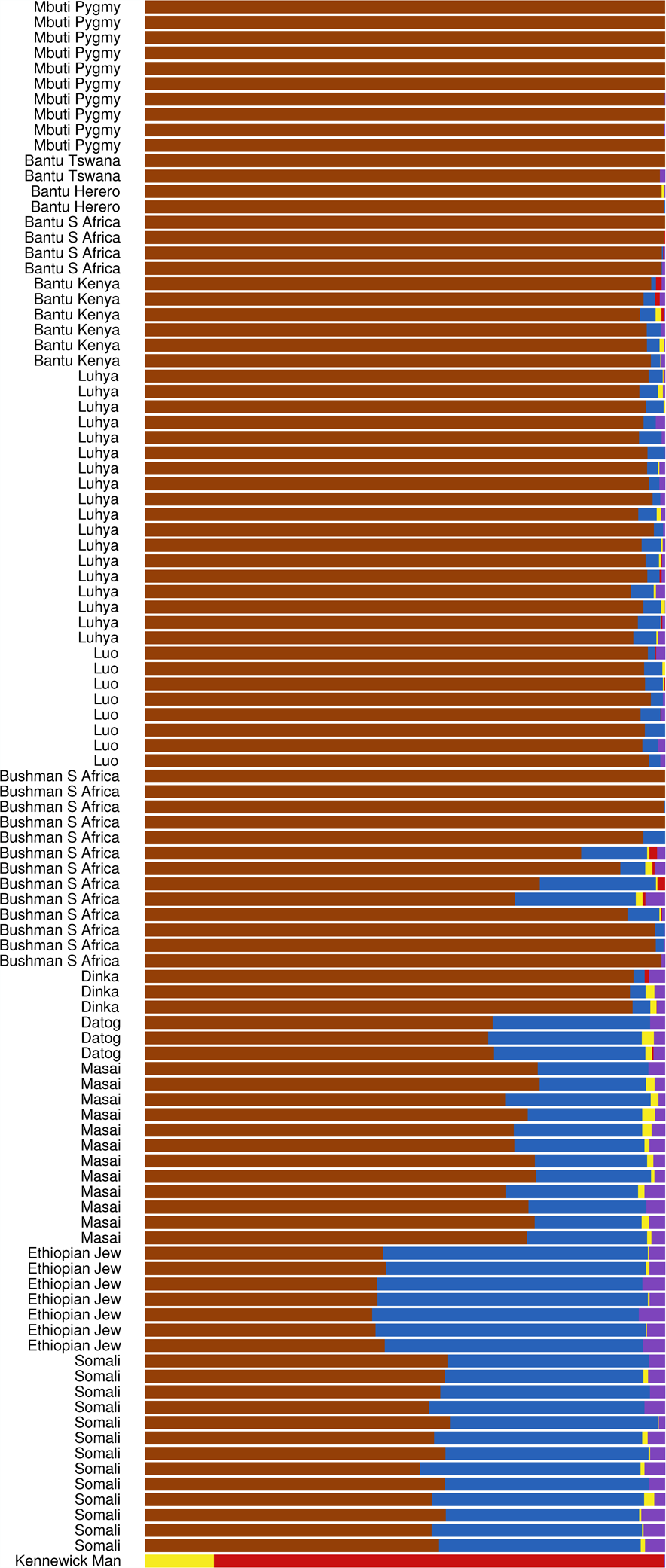
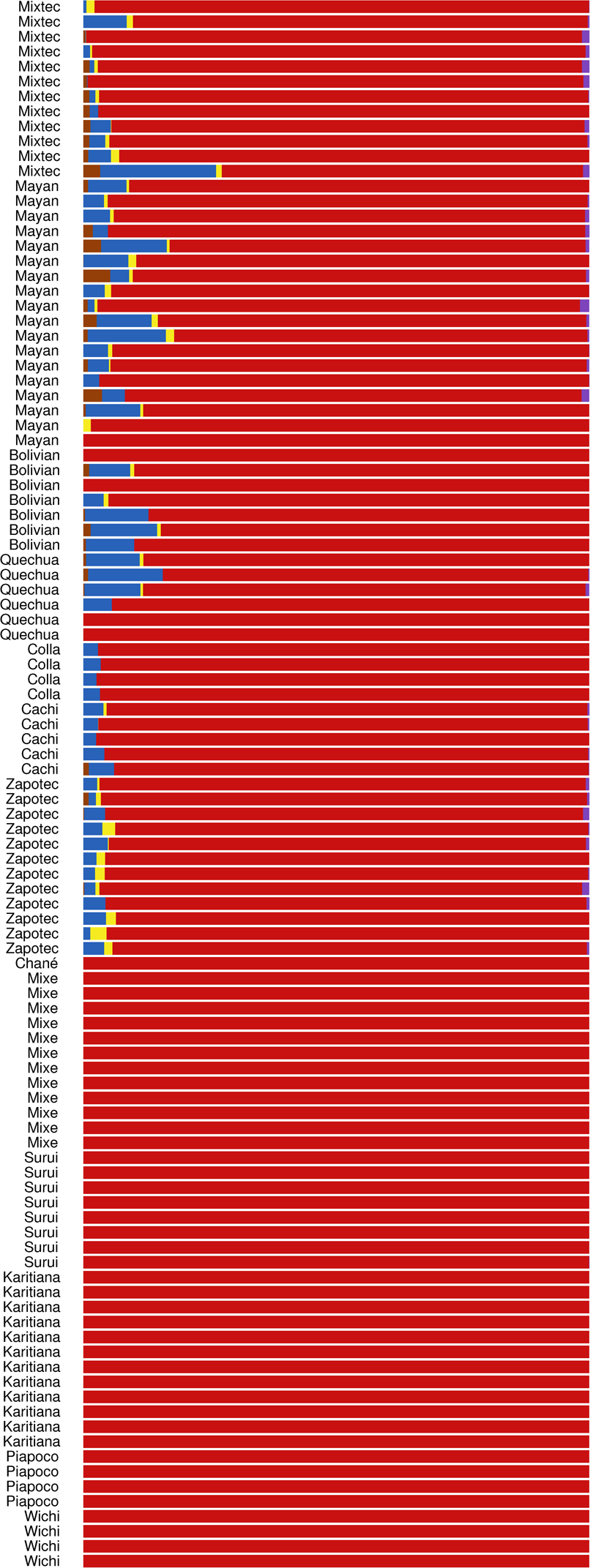
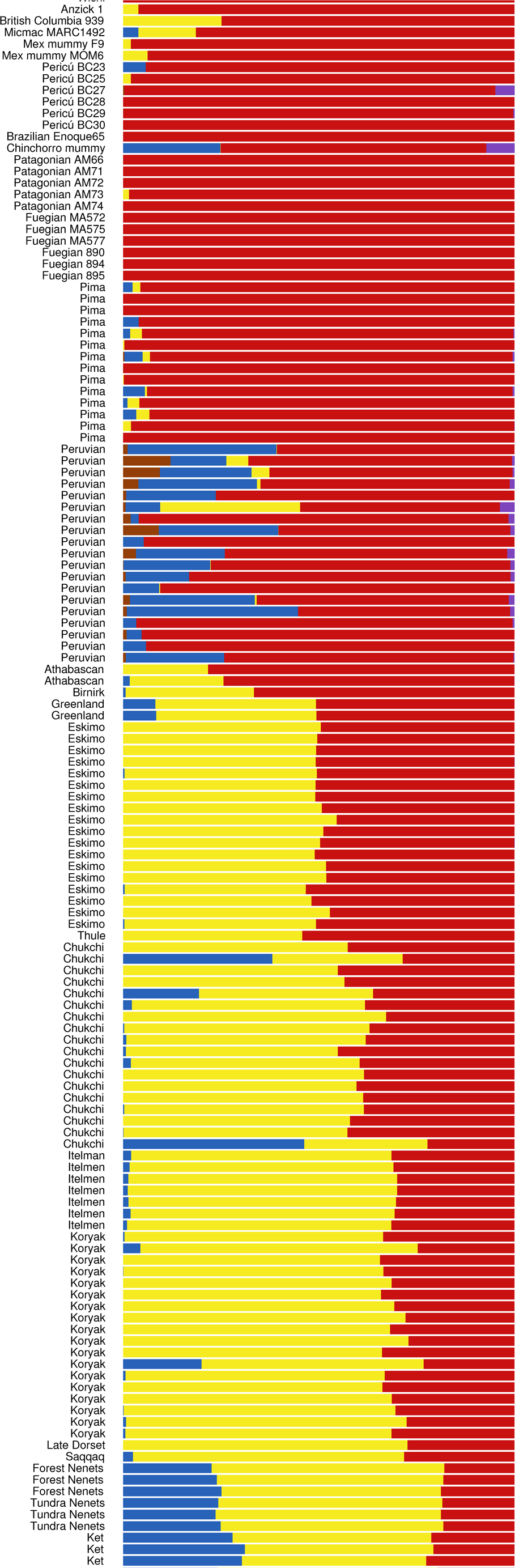
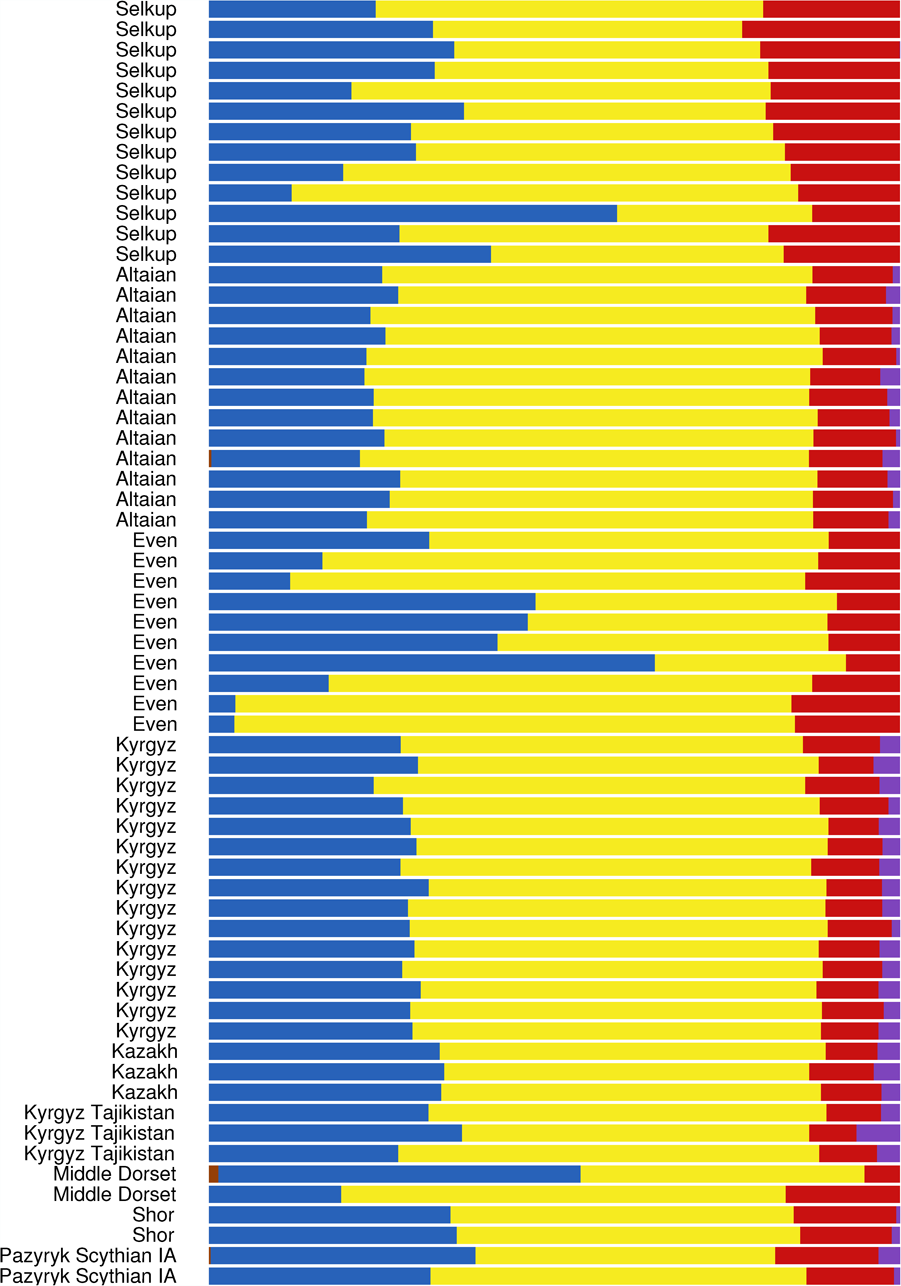
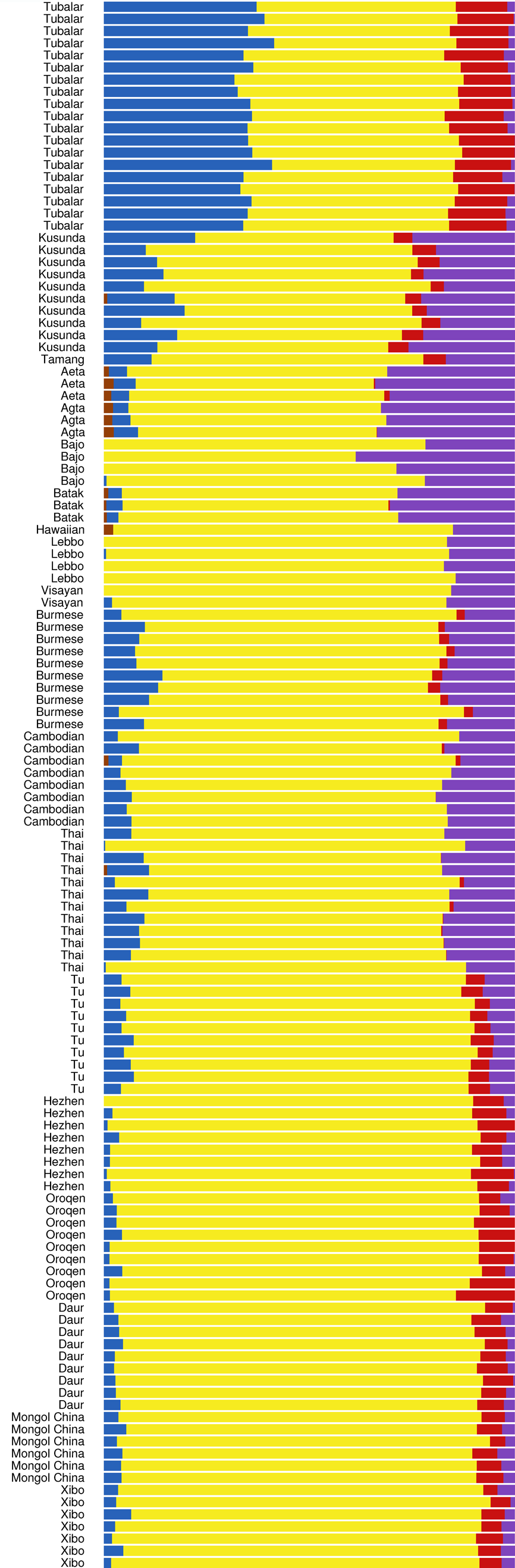
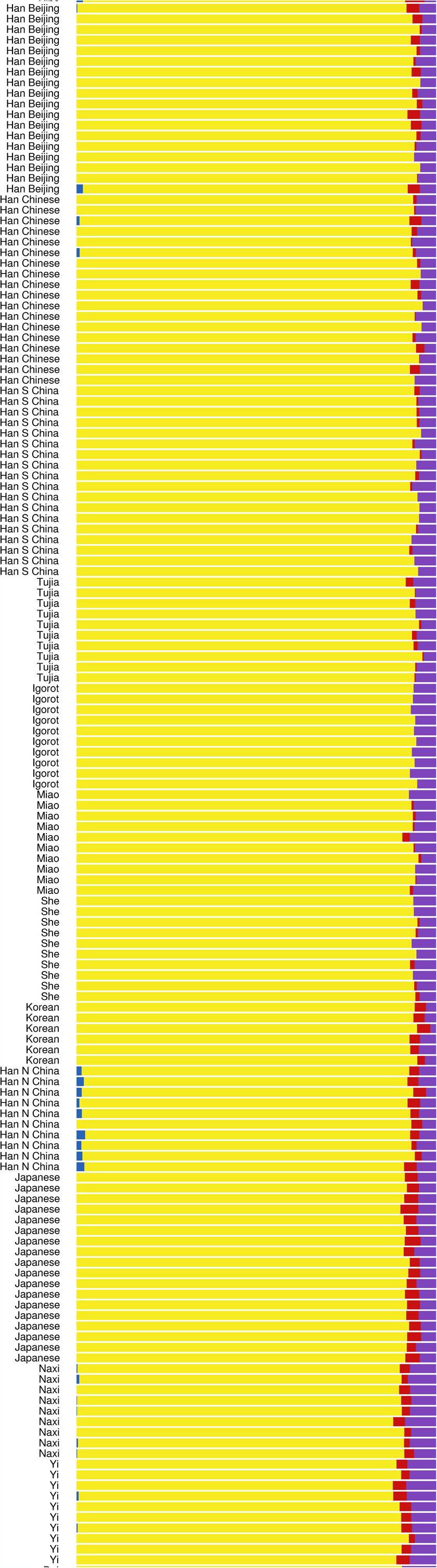
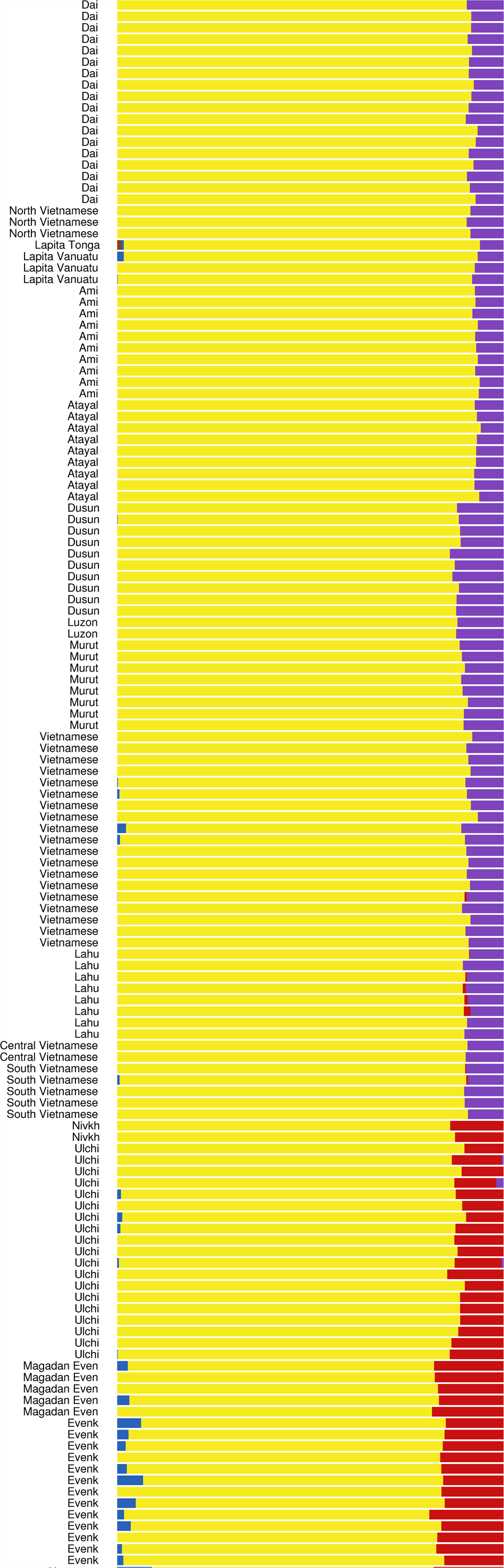
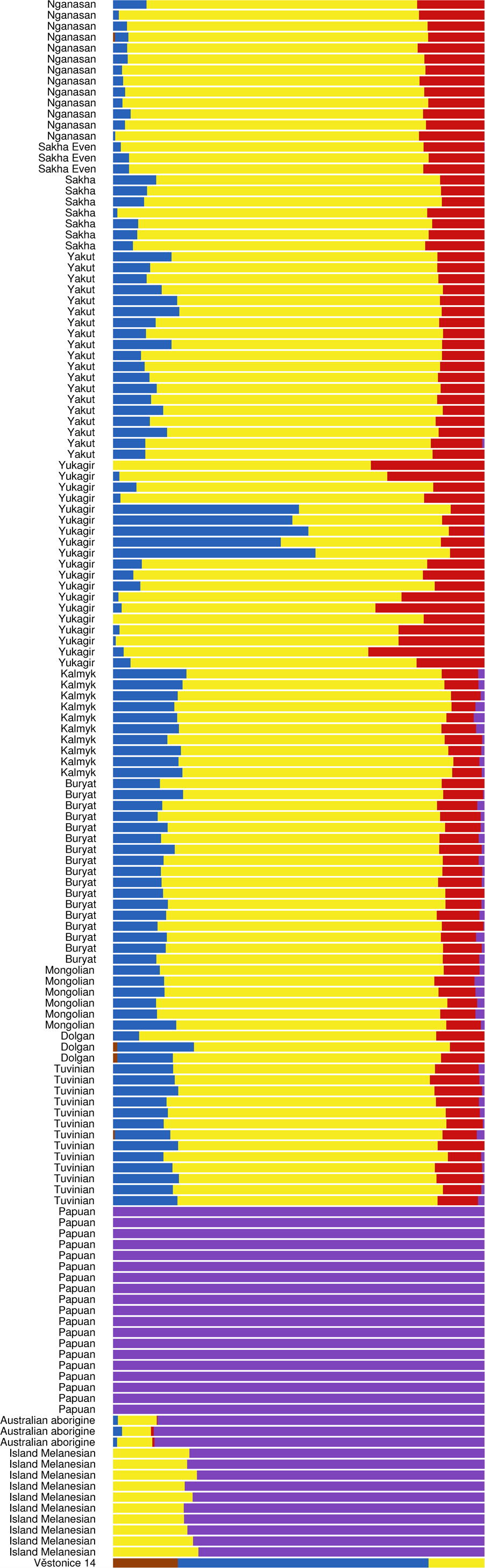
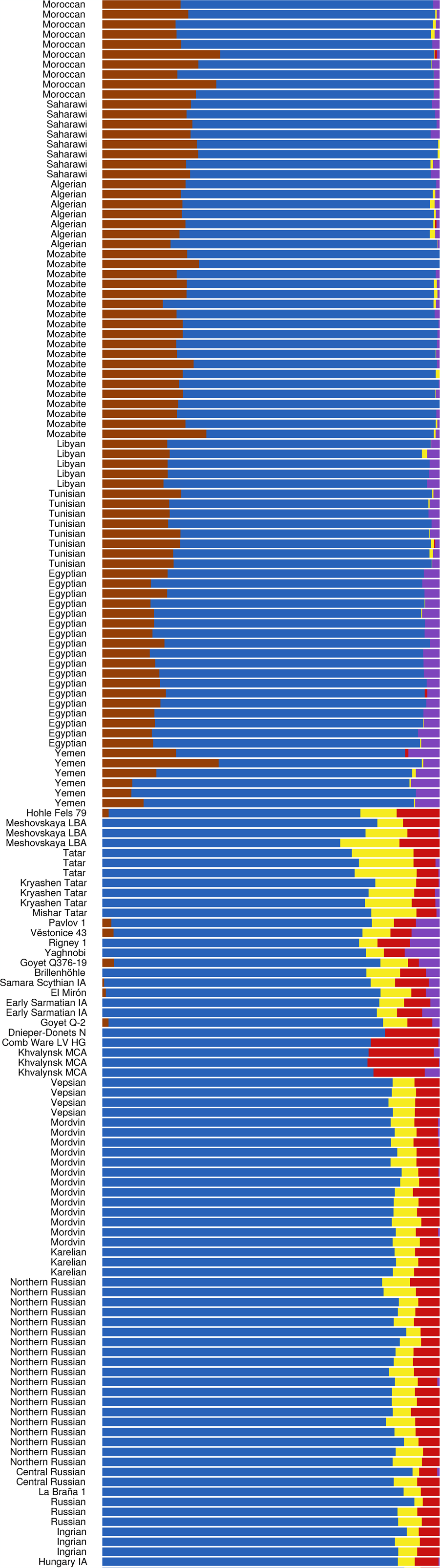
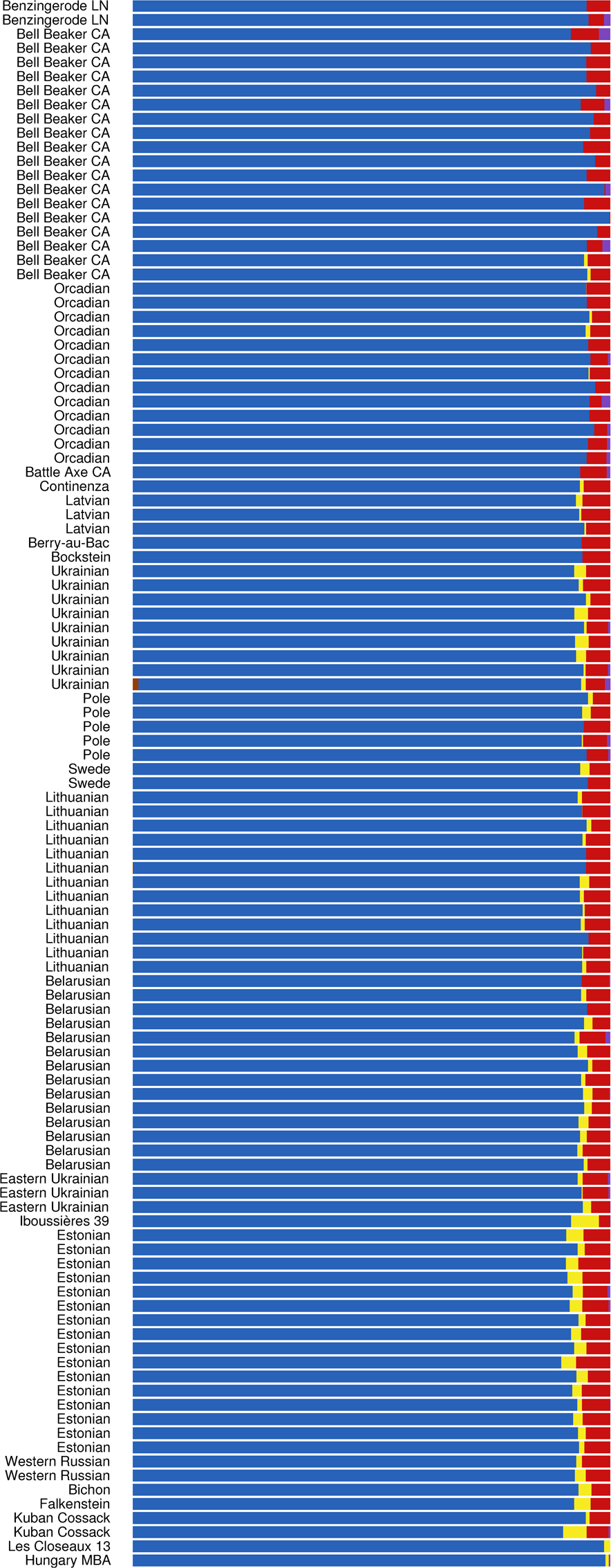
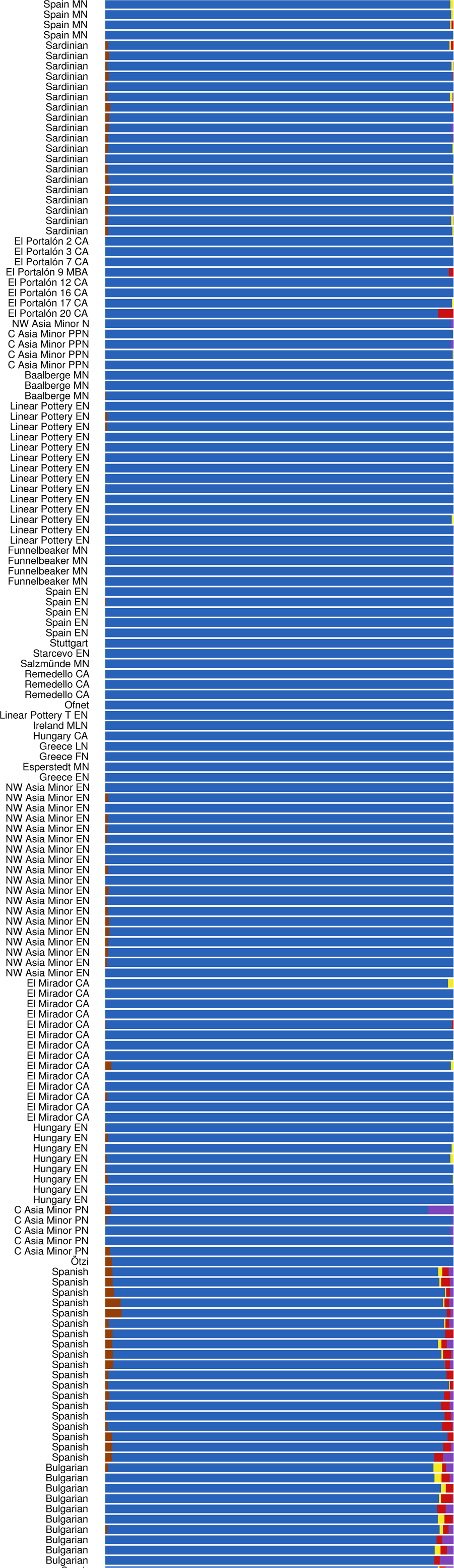
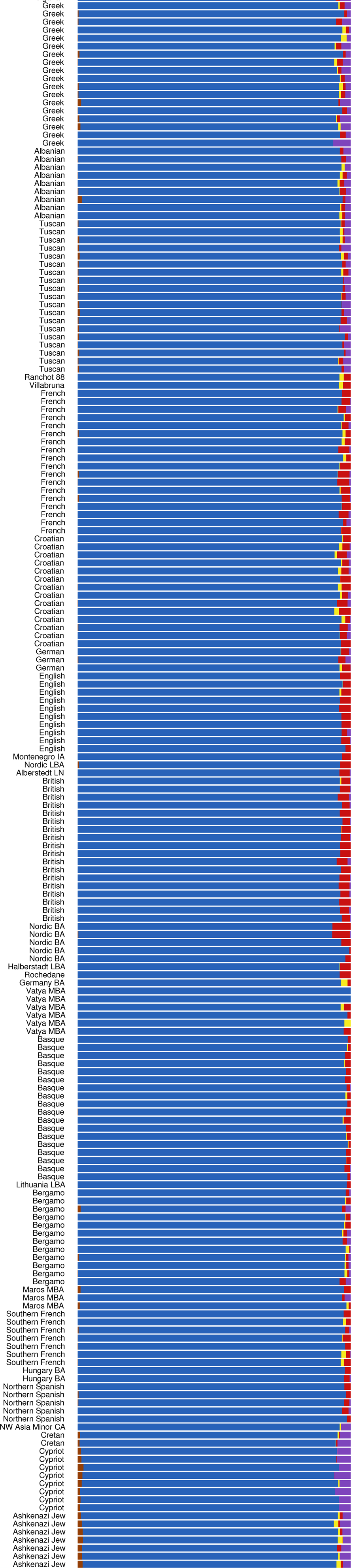
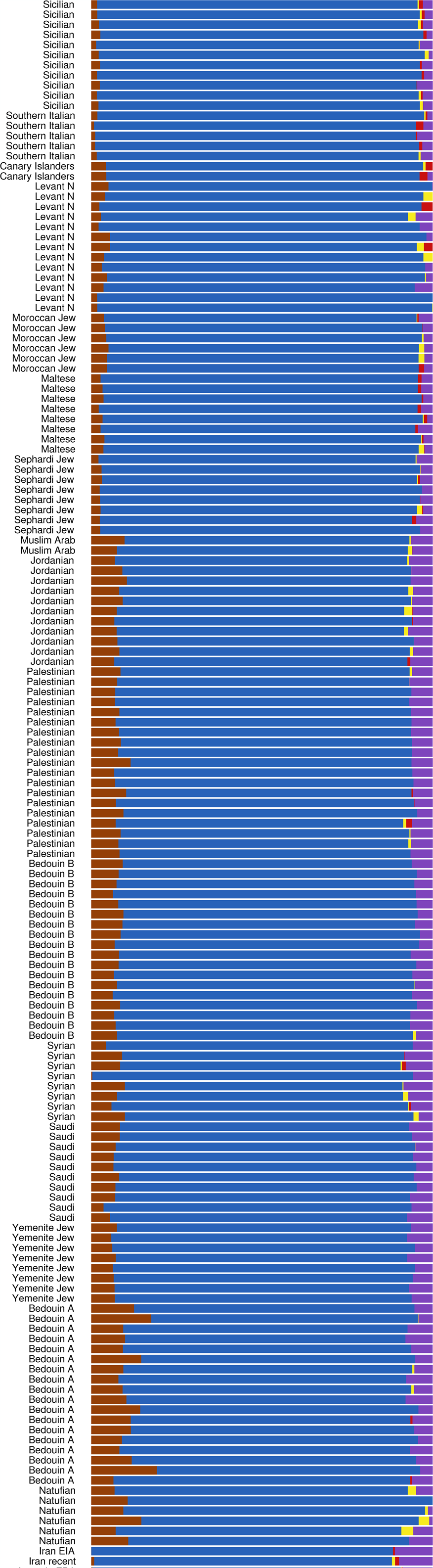
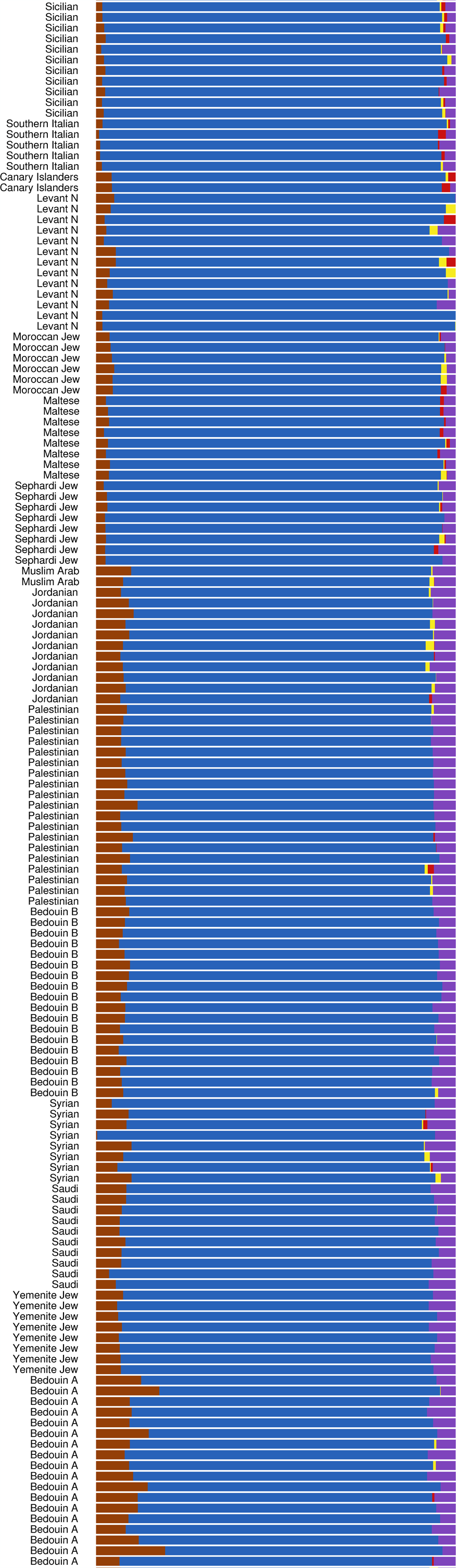
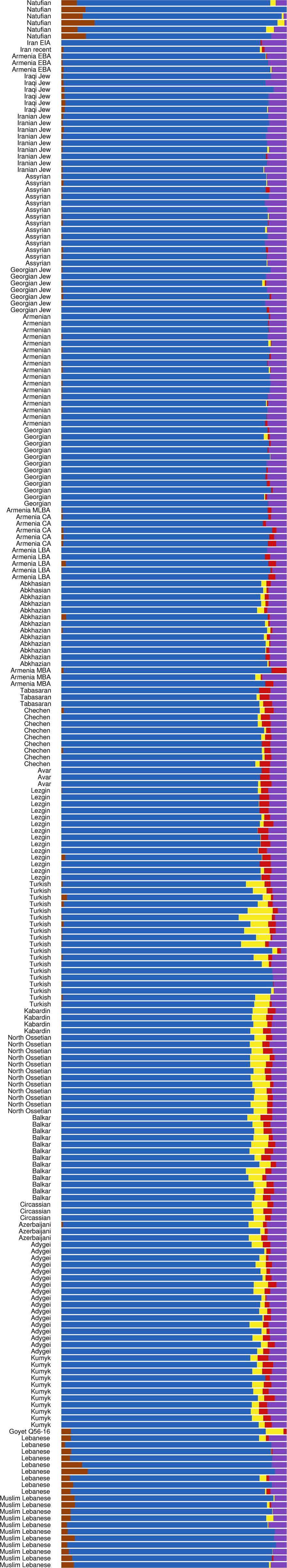
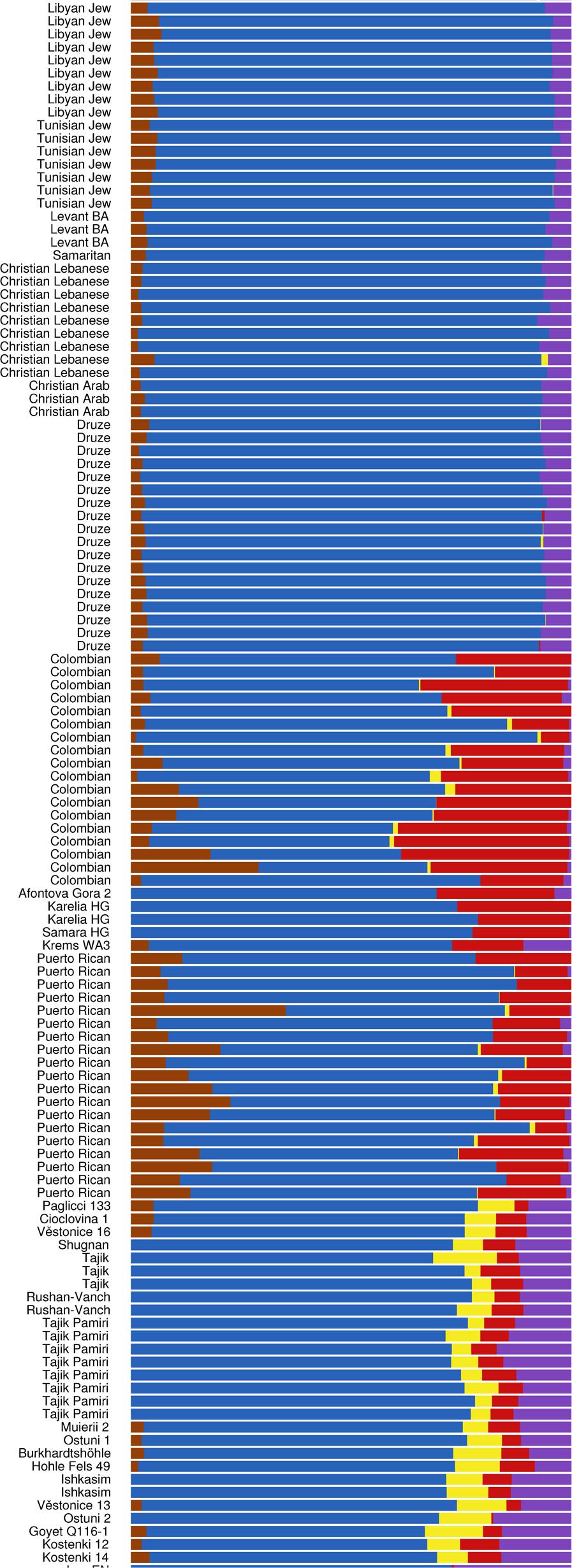
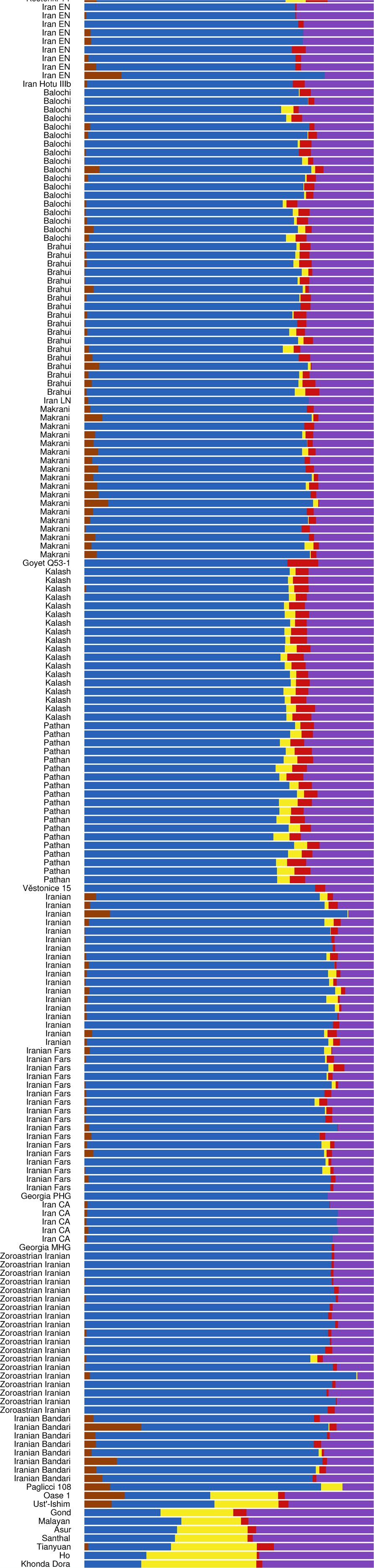
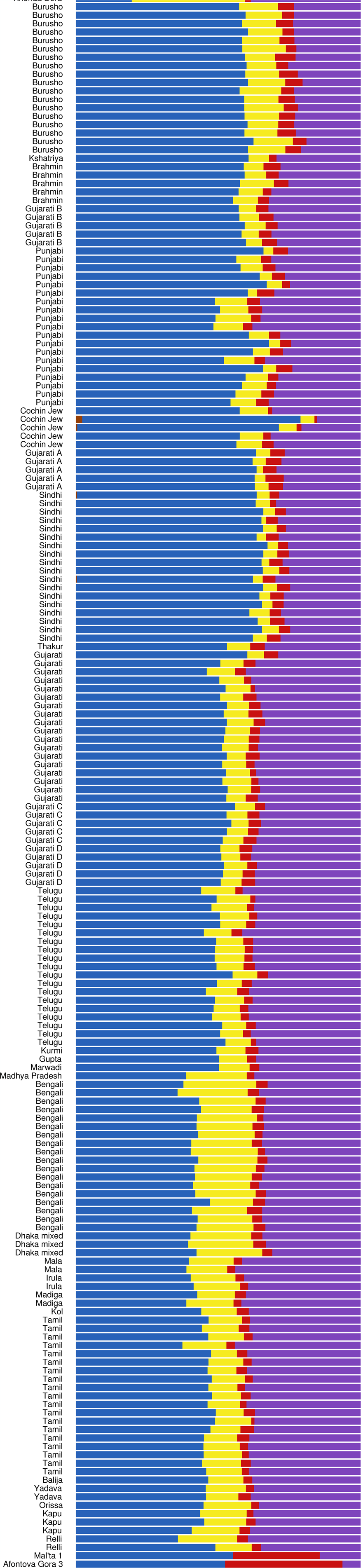
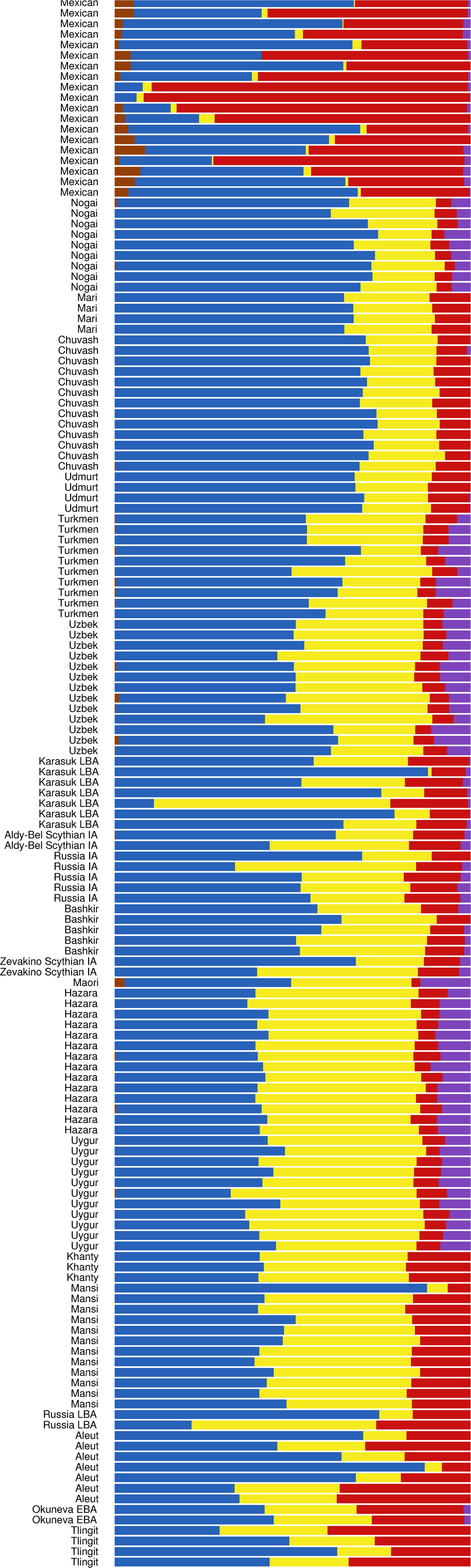
**K = 5 ADMIXTURE analysis**

**Figure 5:**
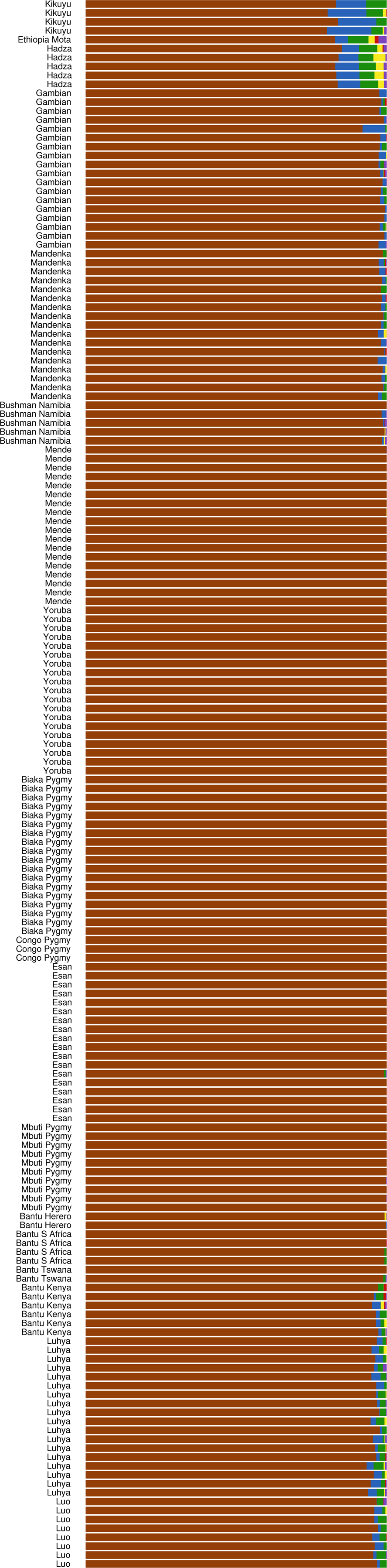
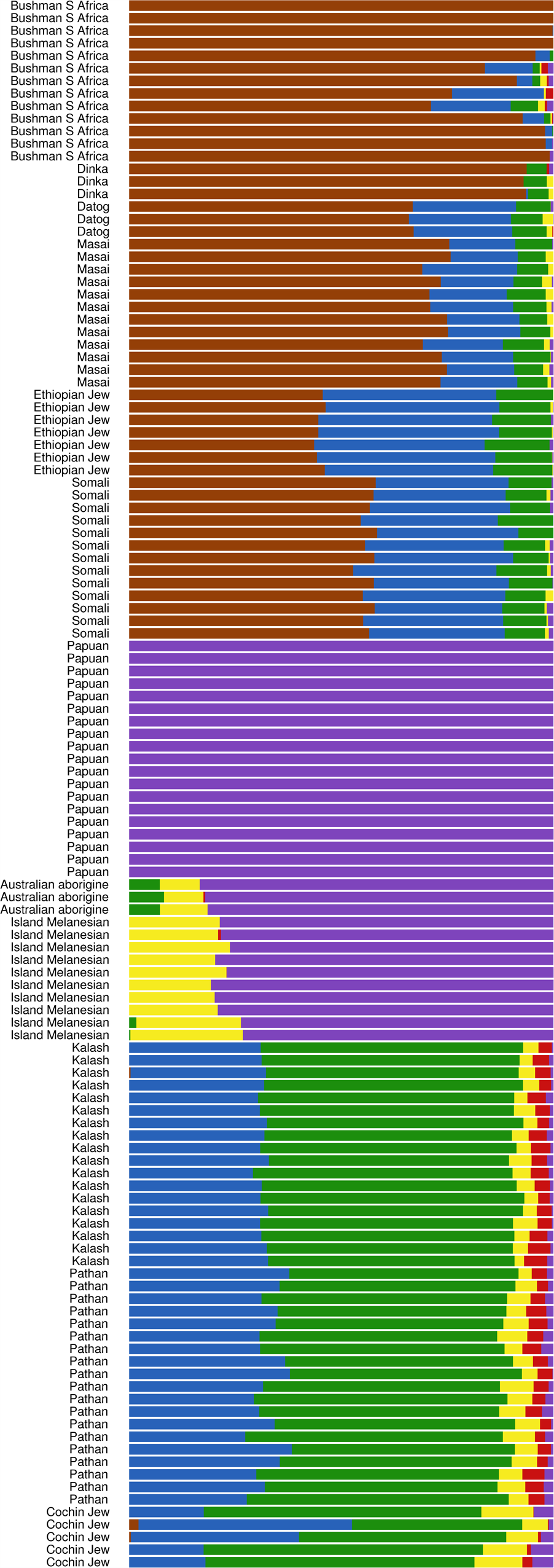
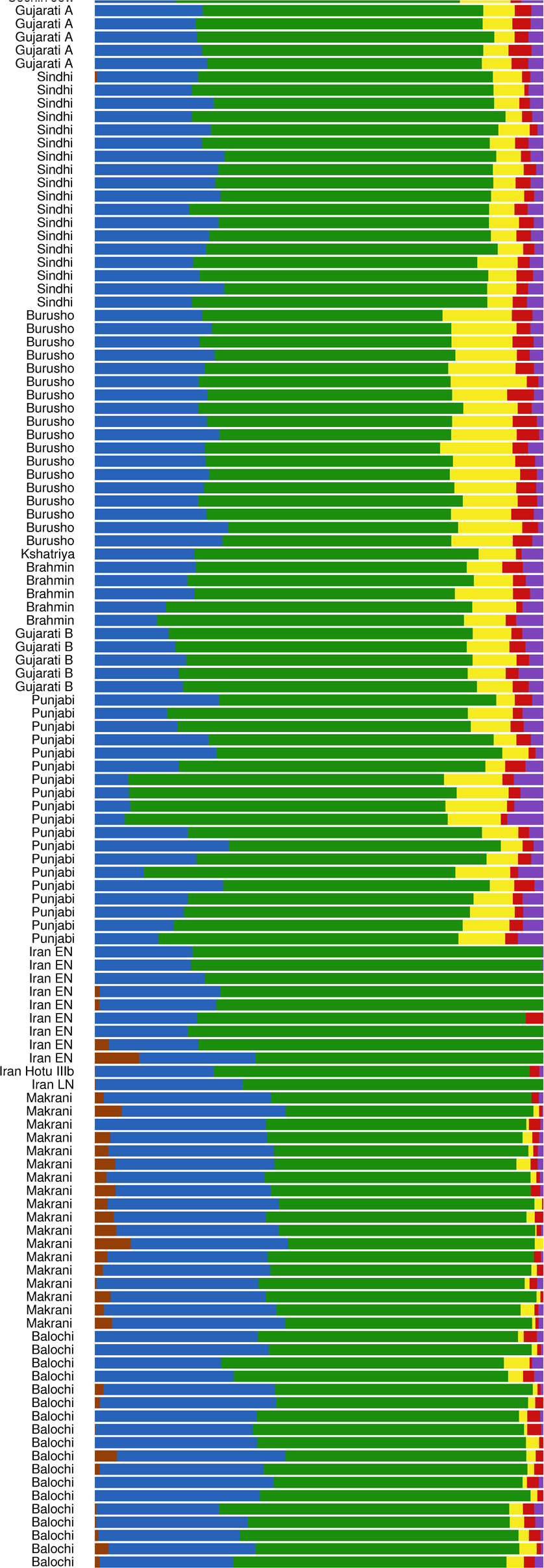
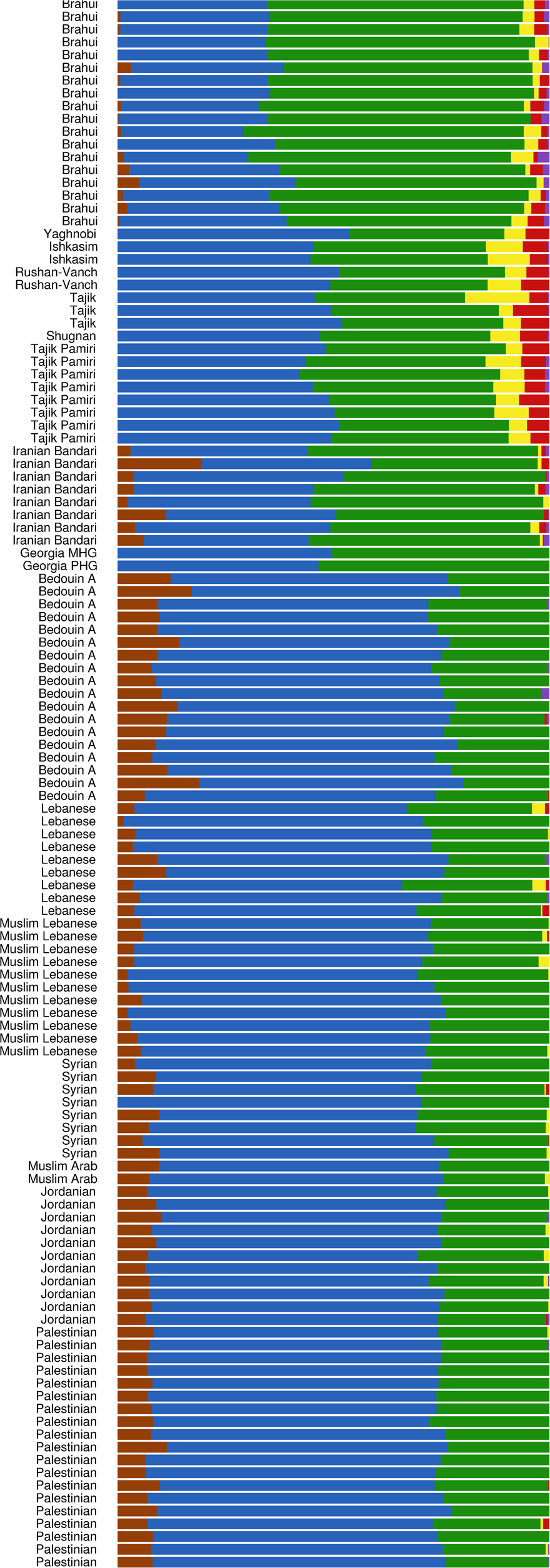
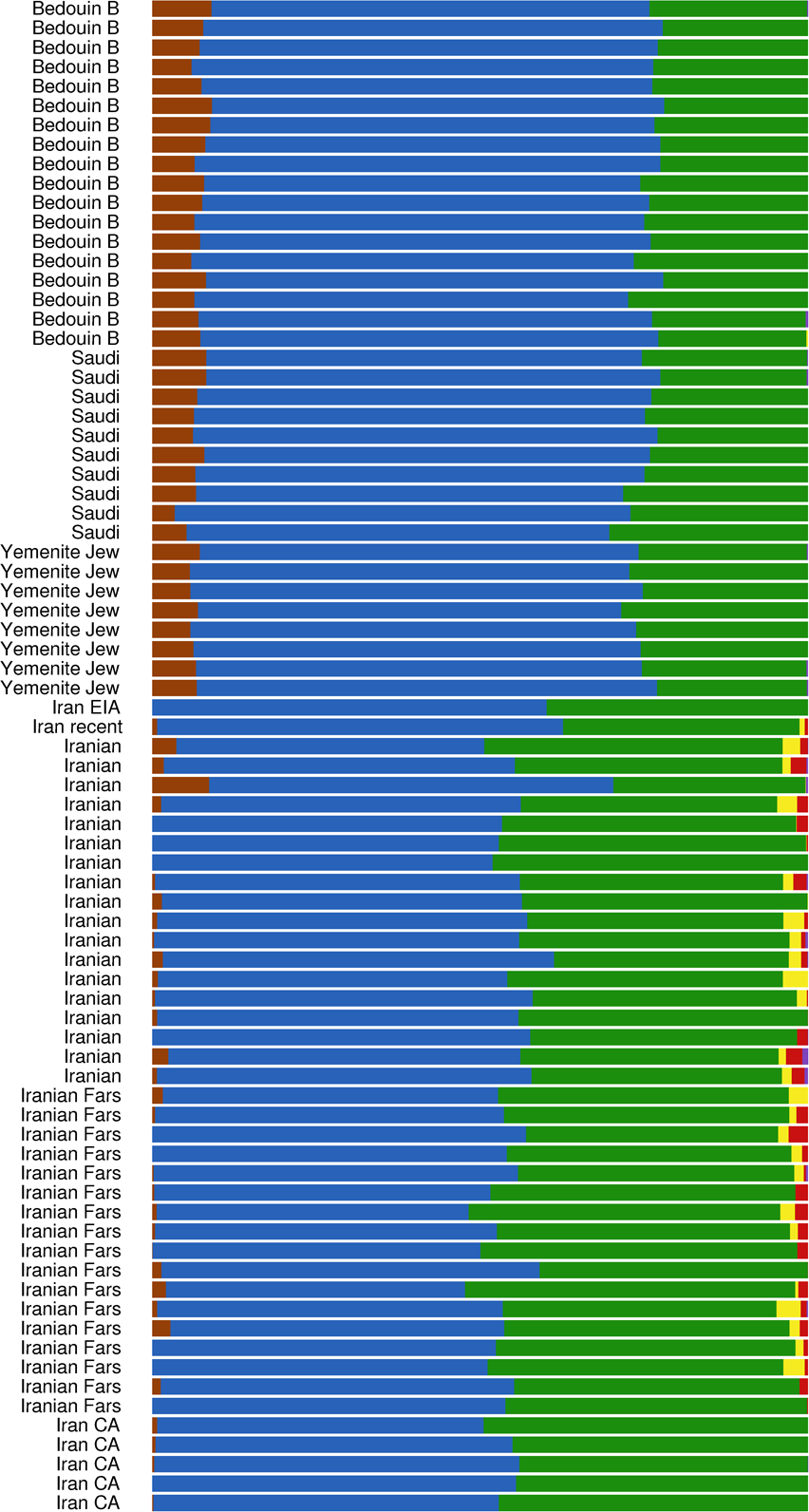
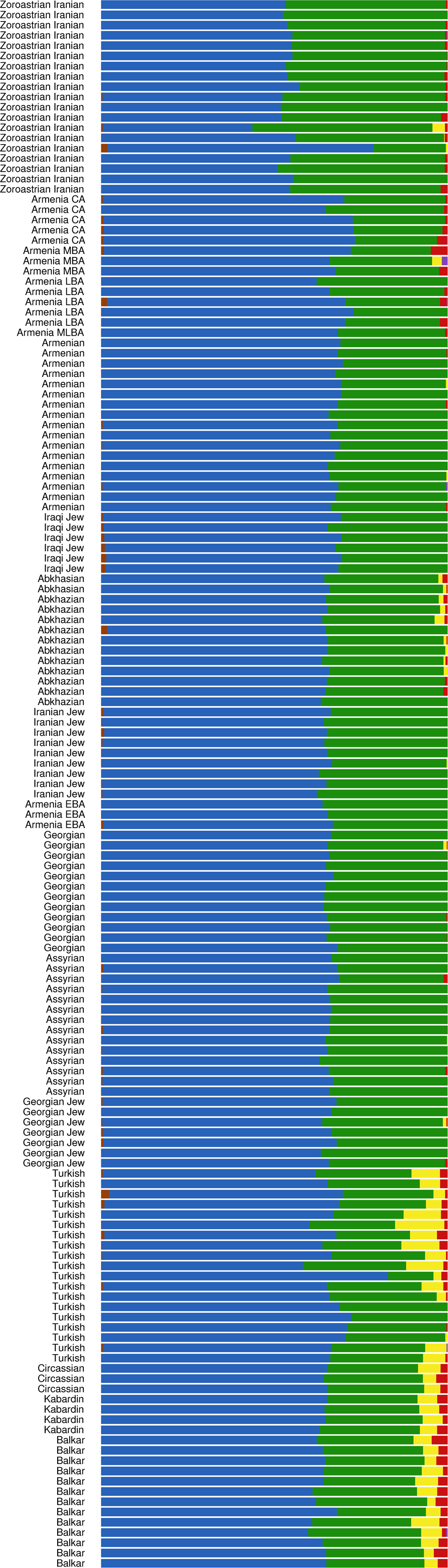
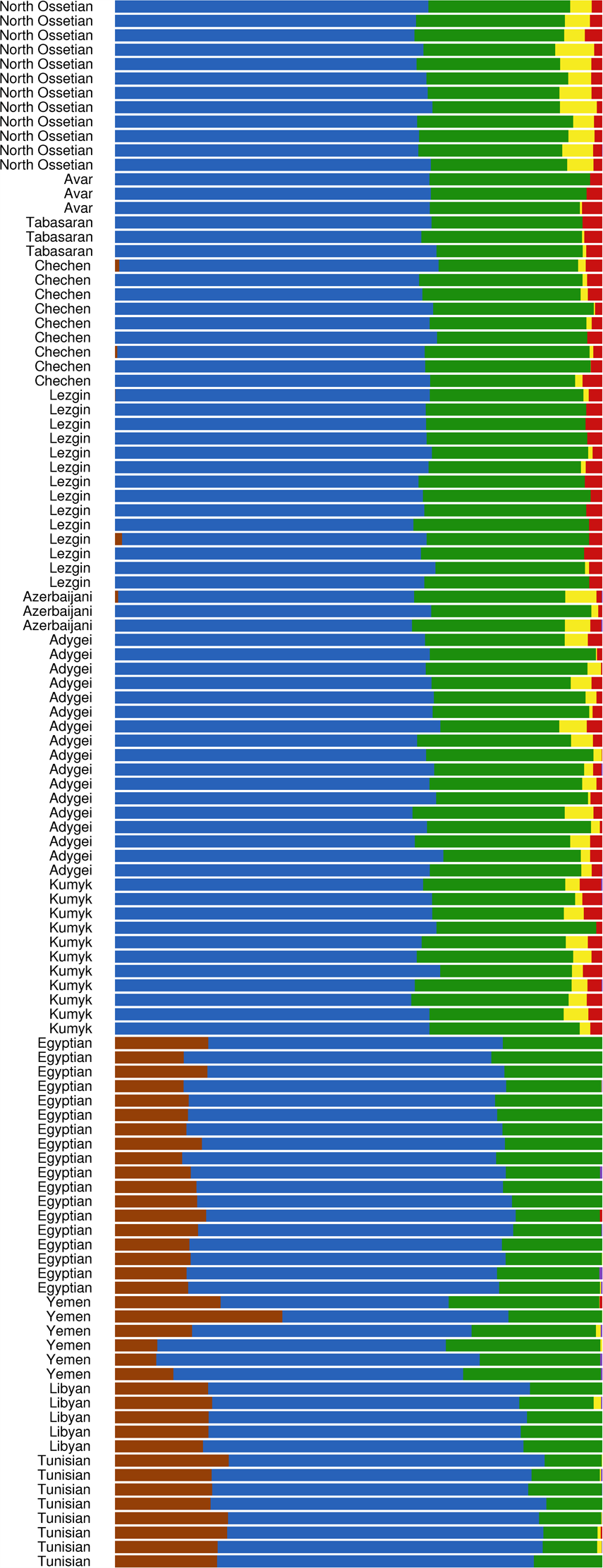
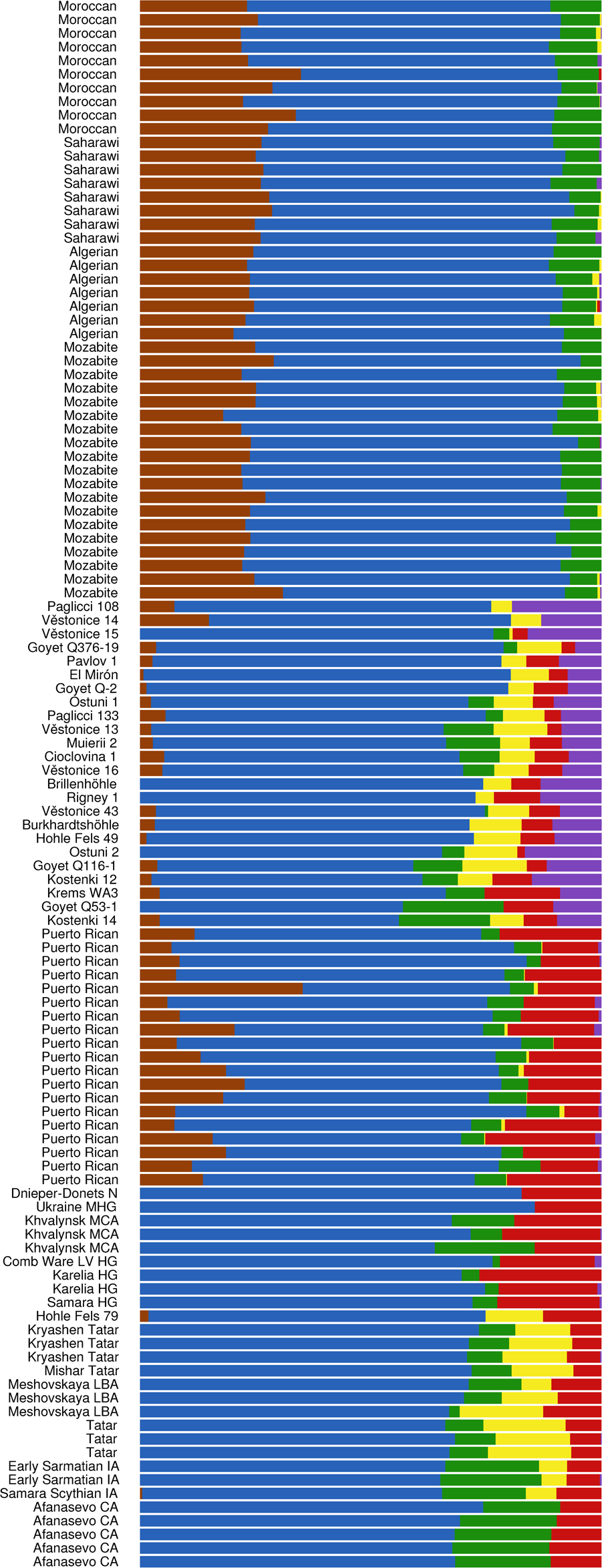
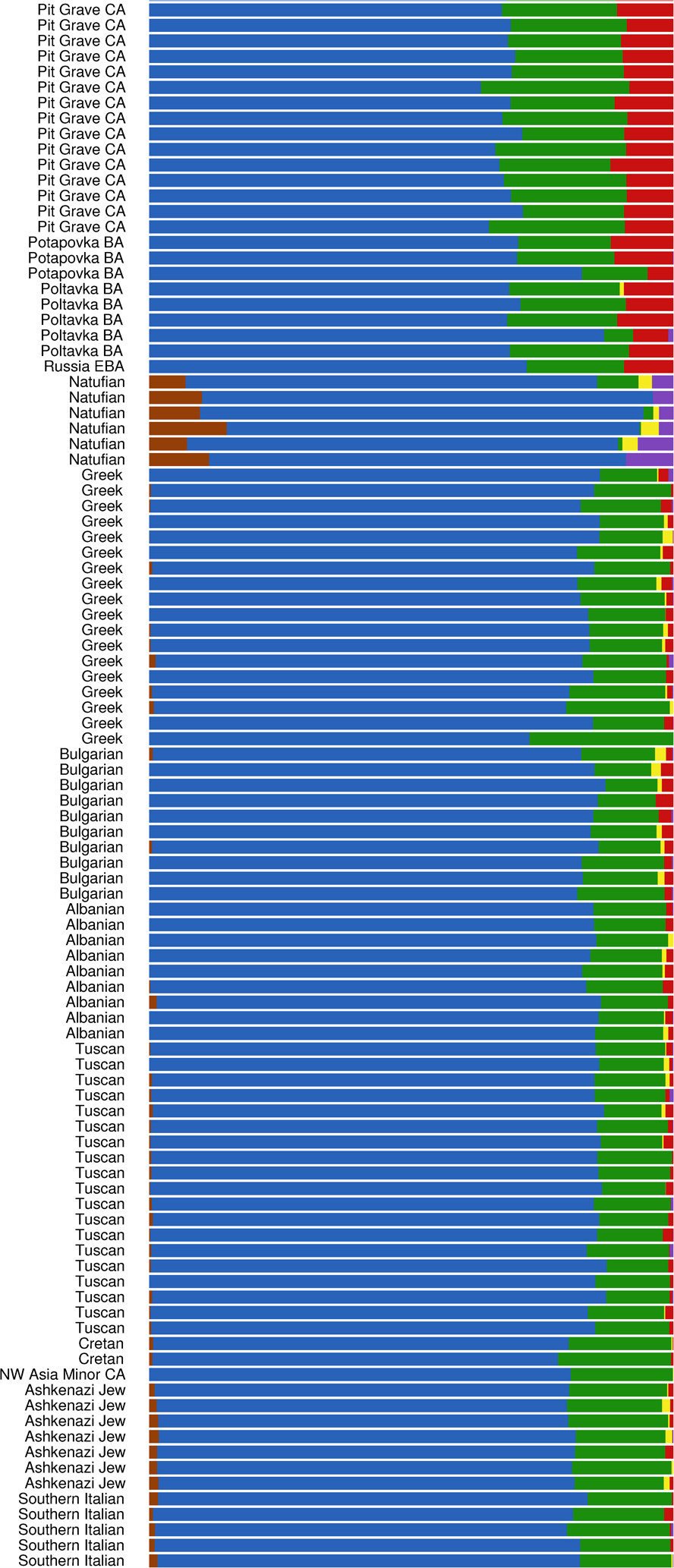
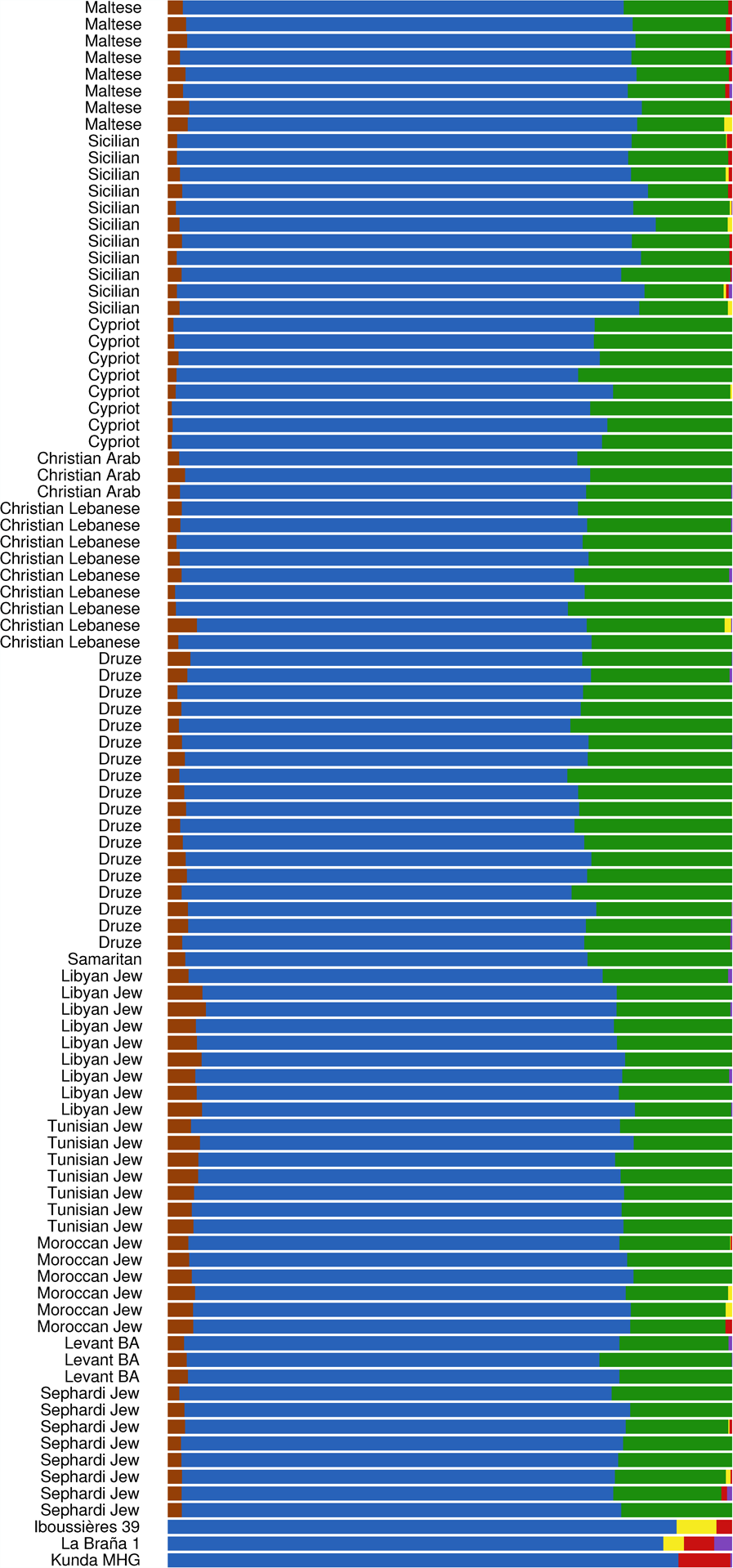
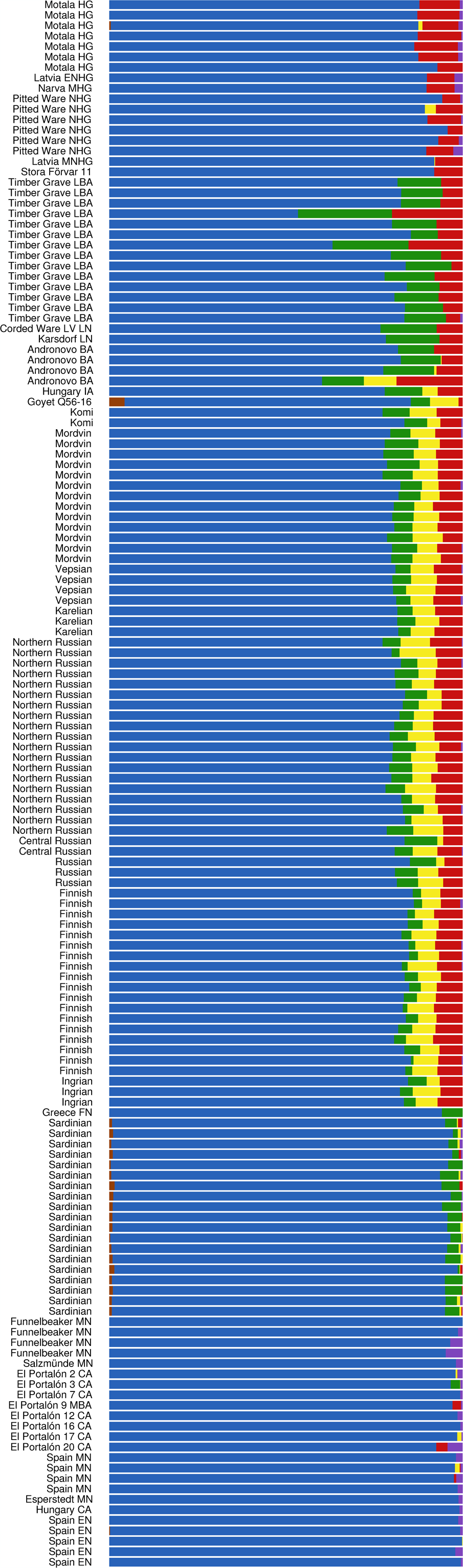
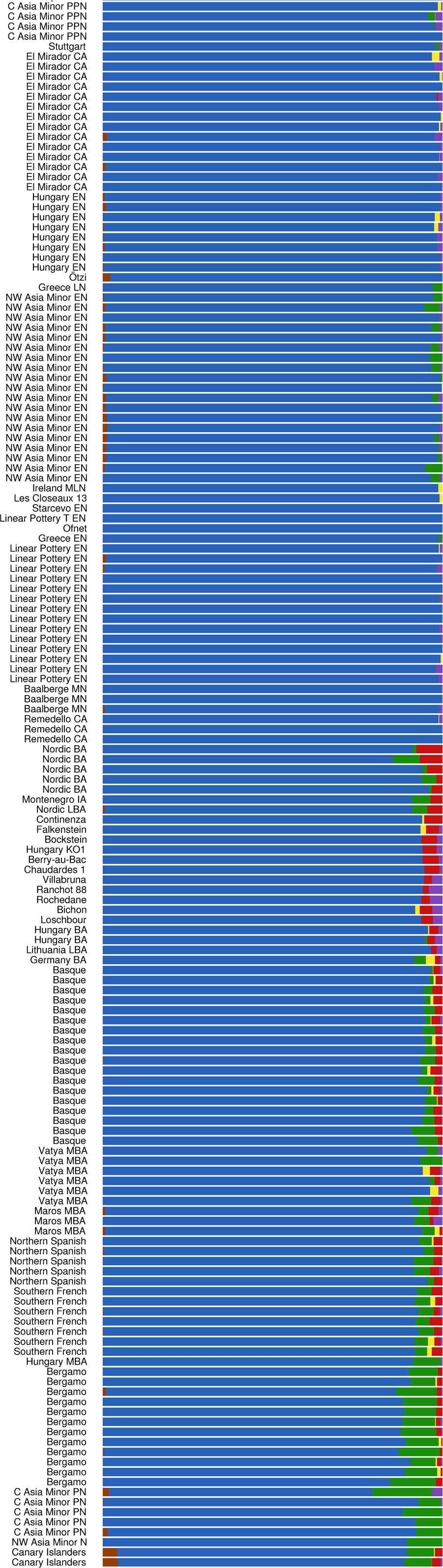
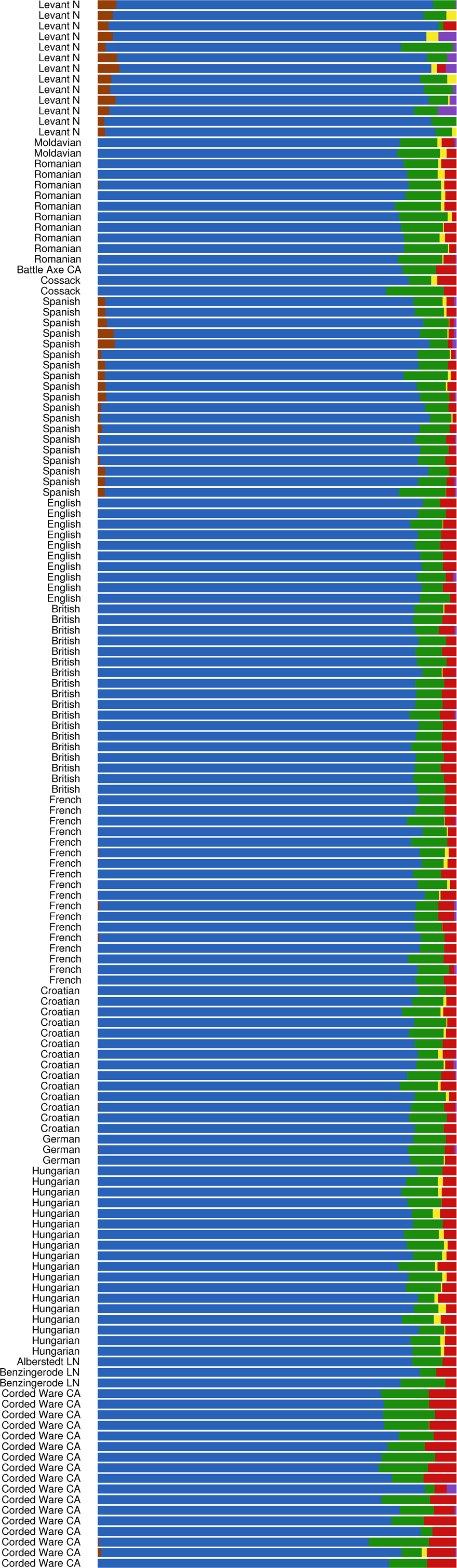
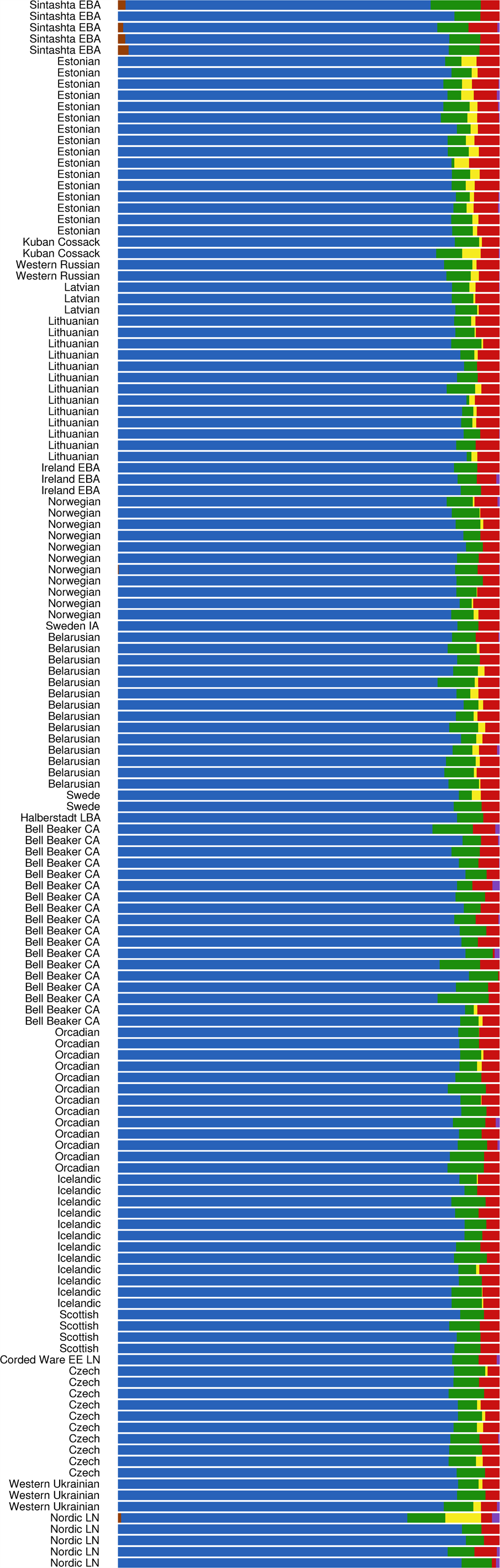
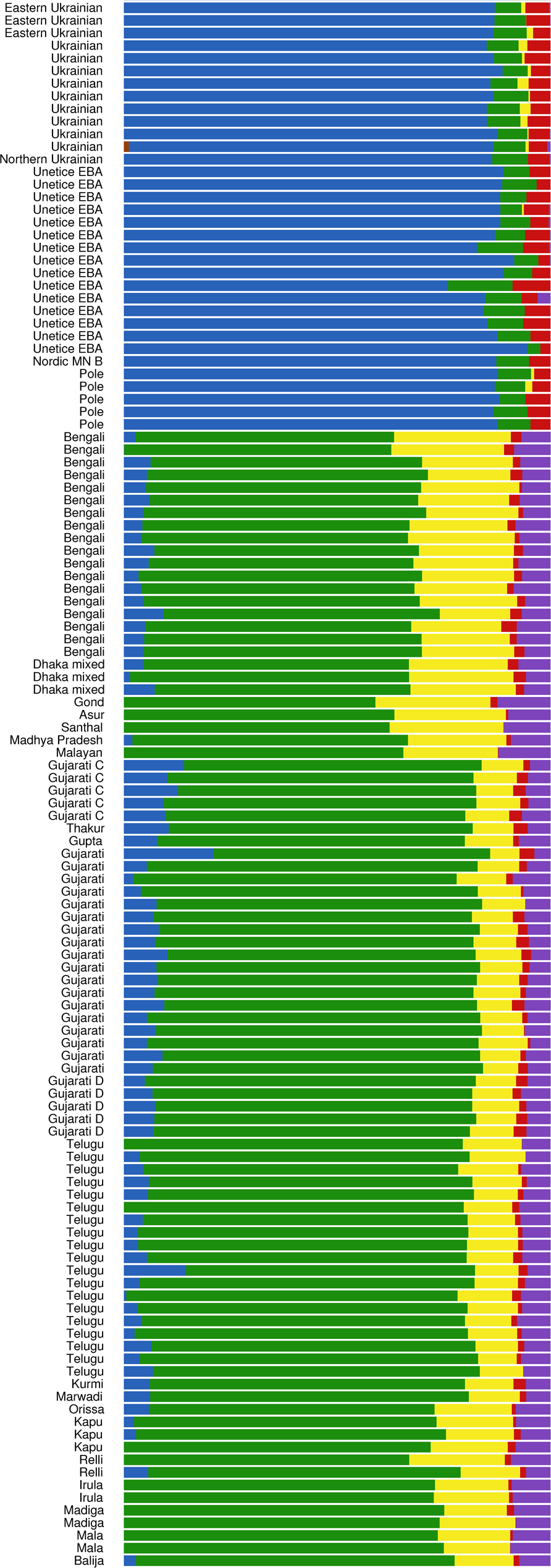
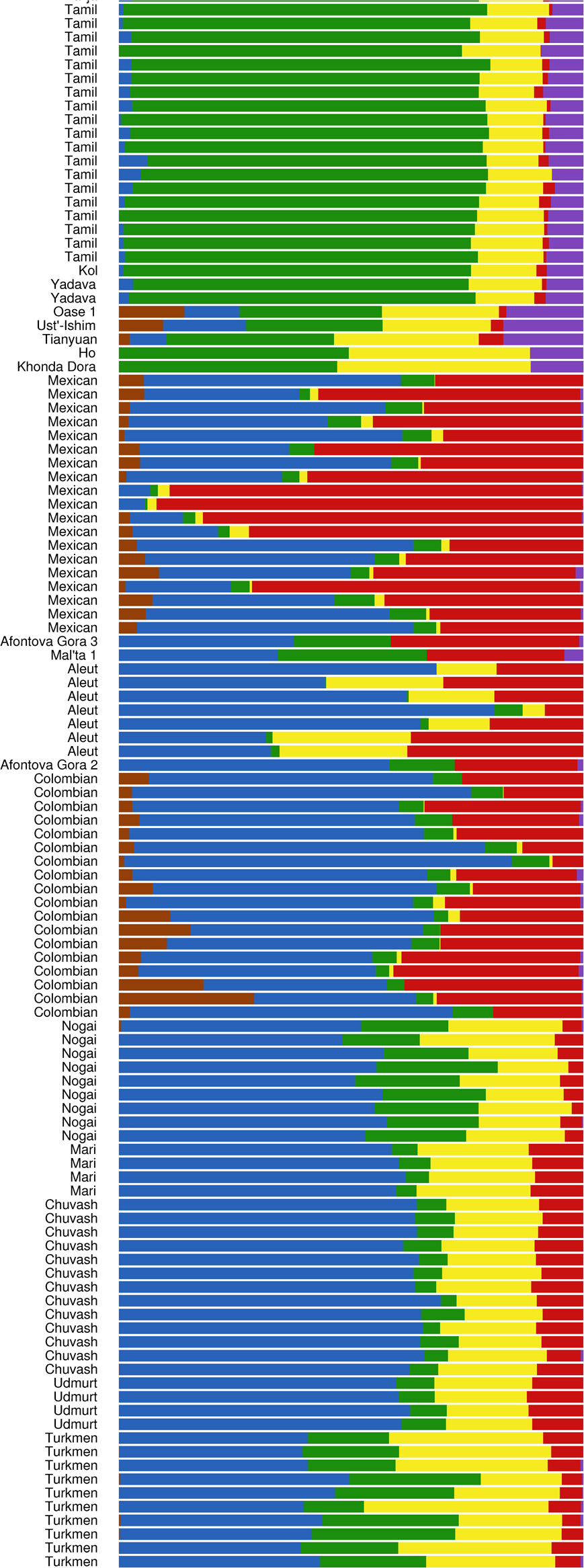
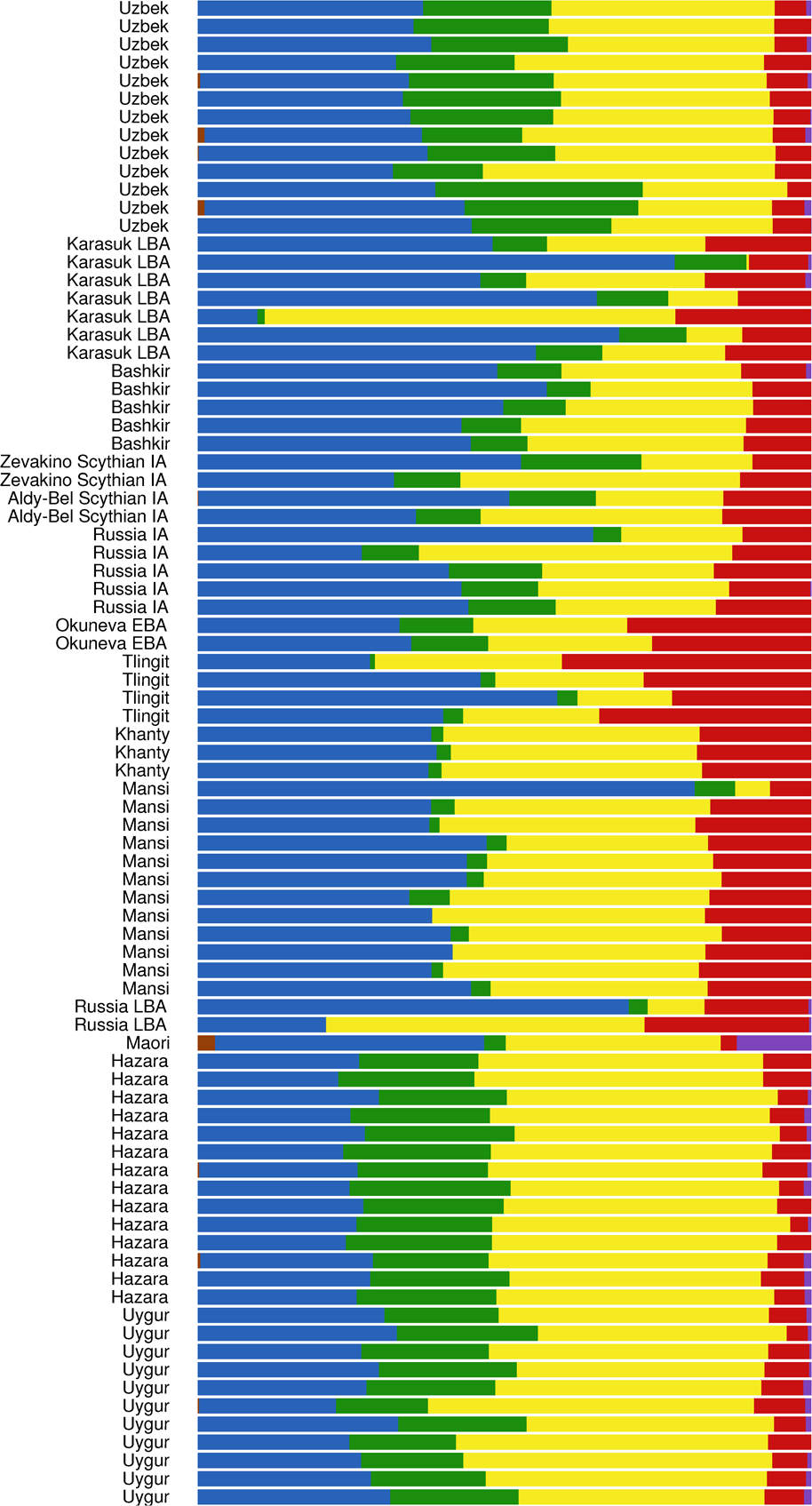
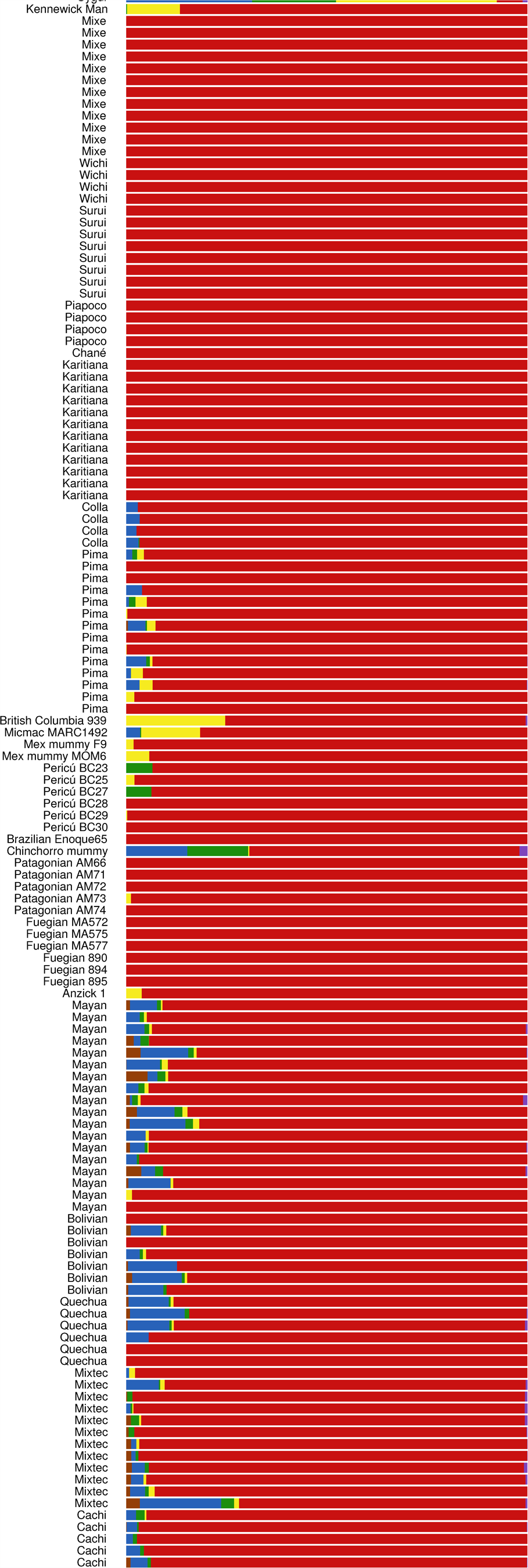
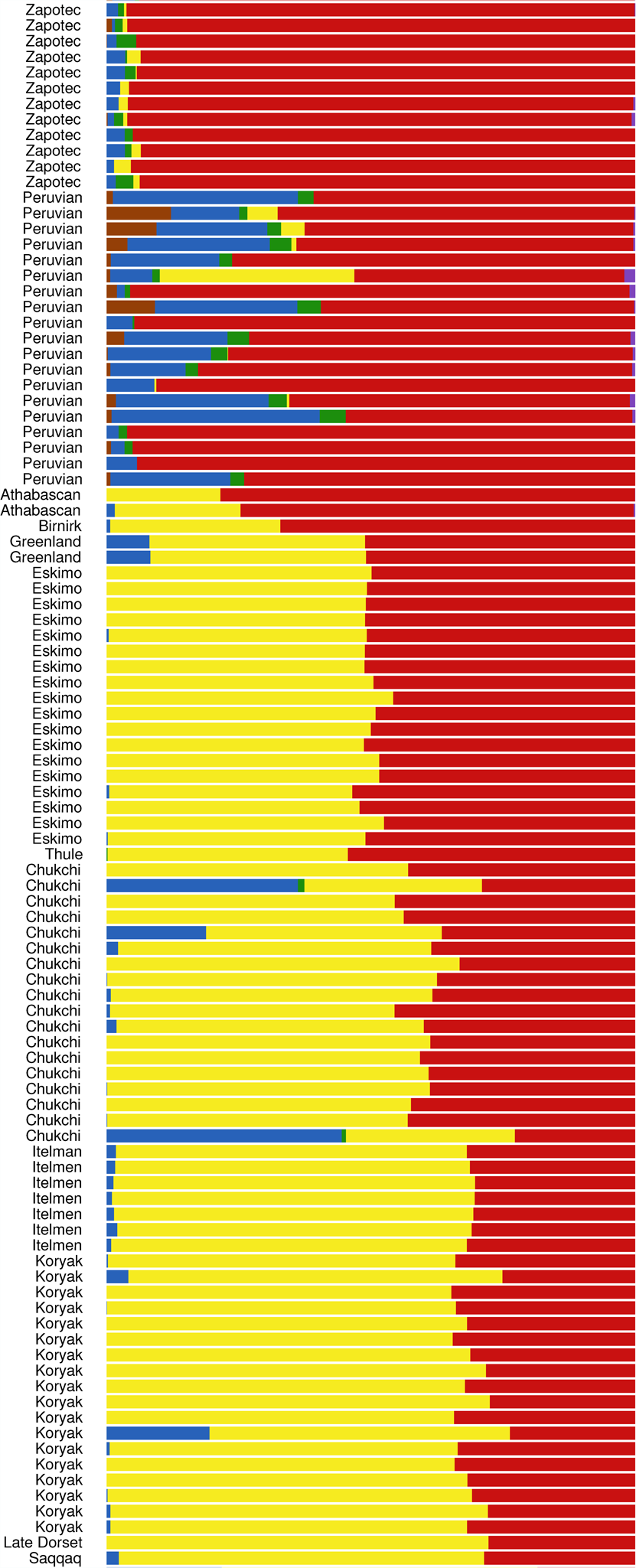
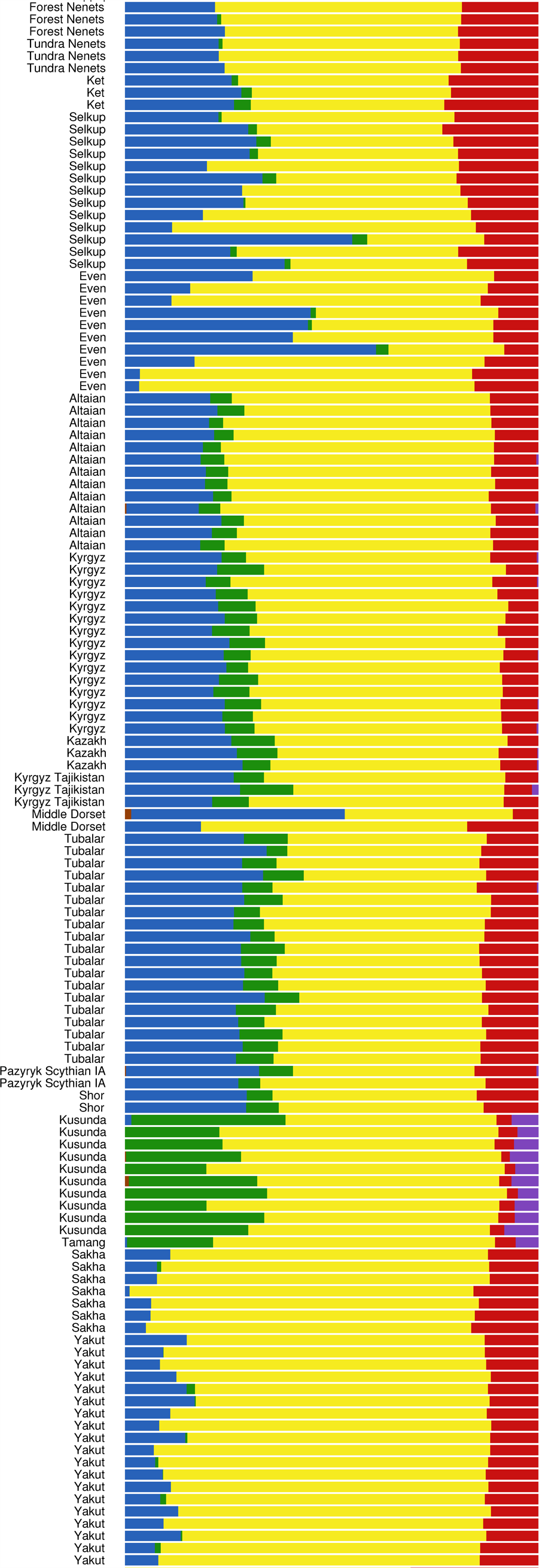
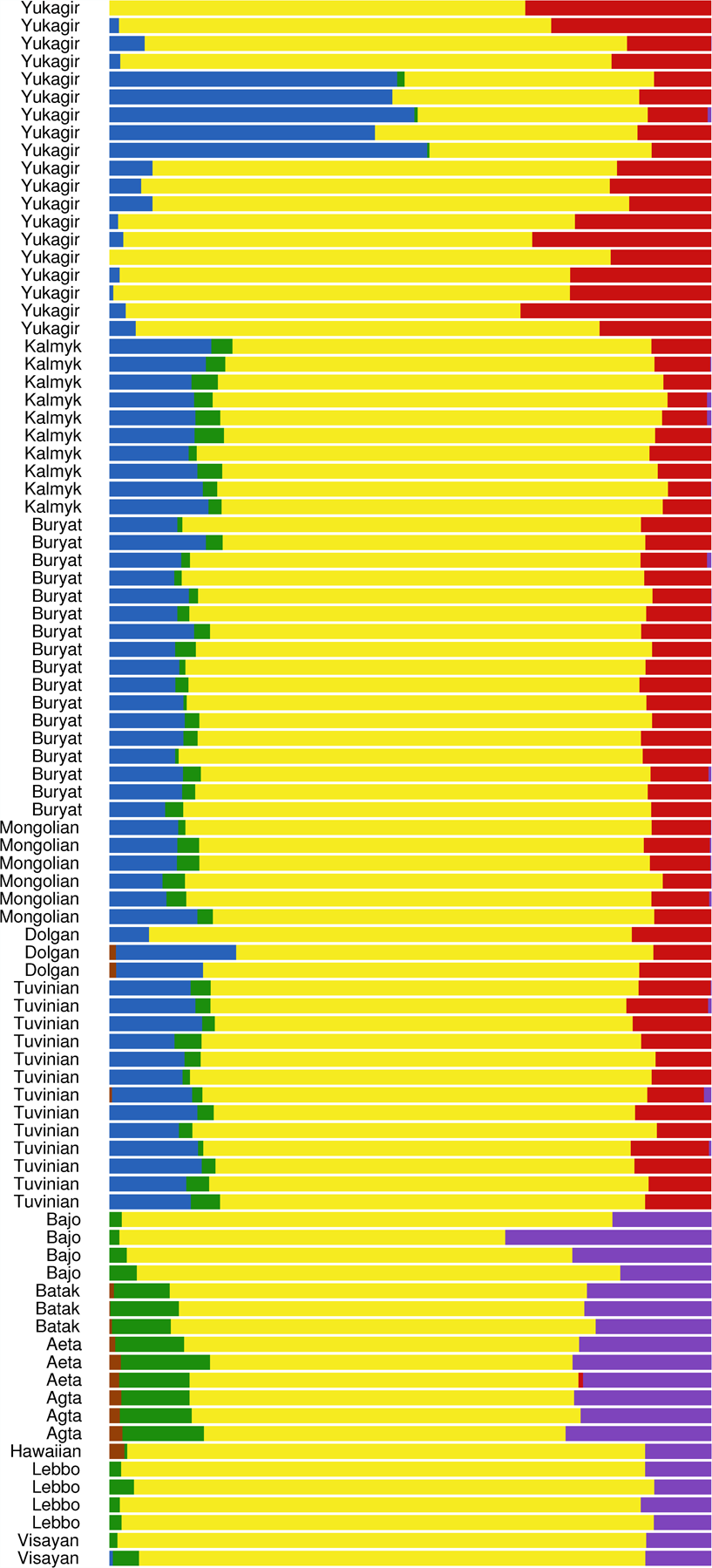
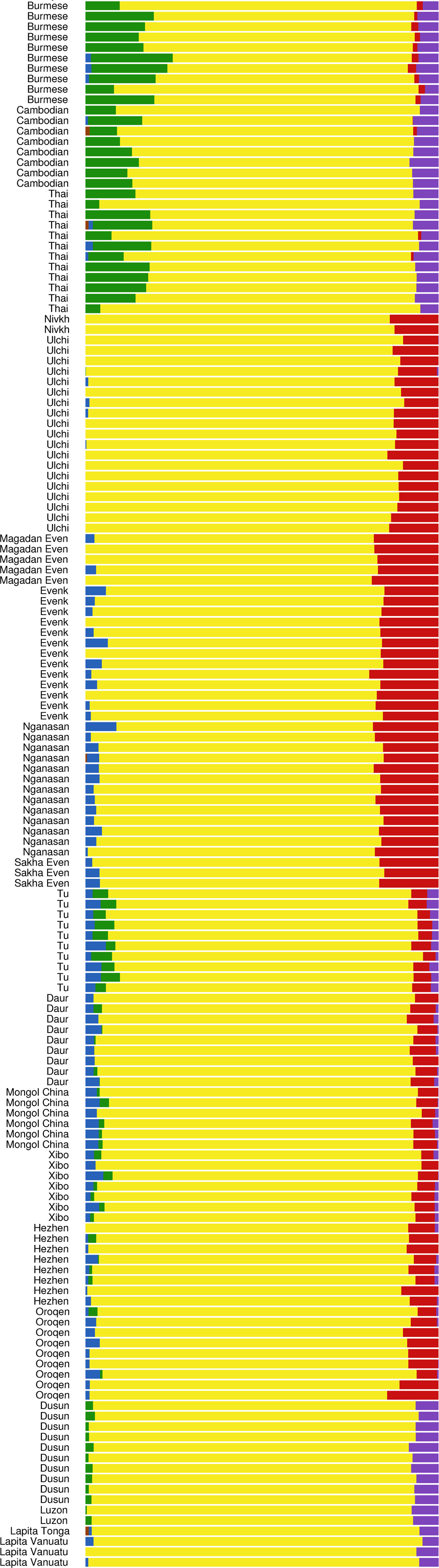
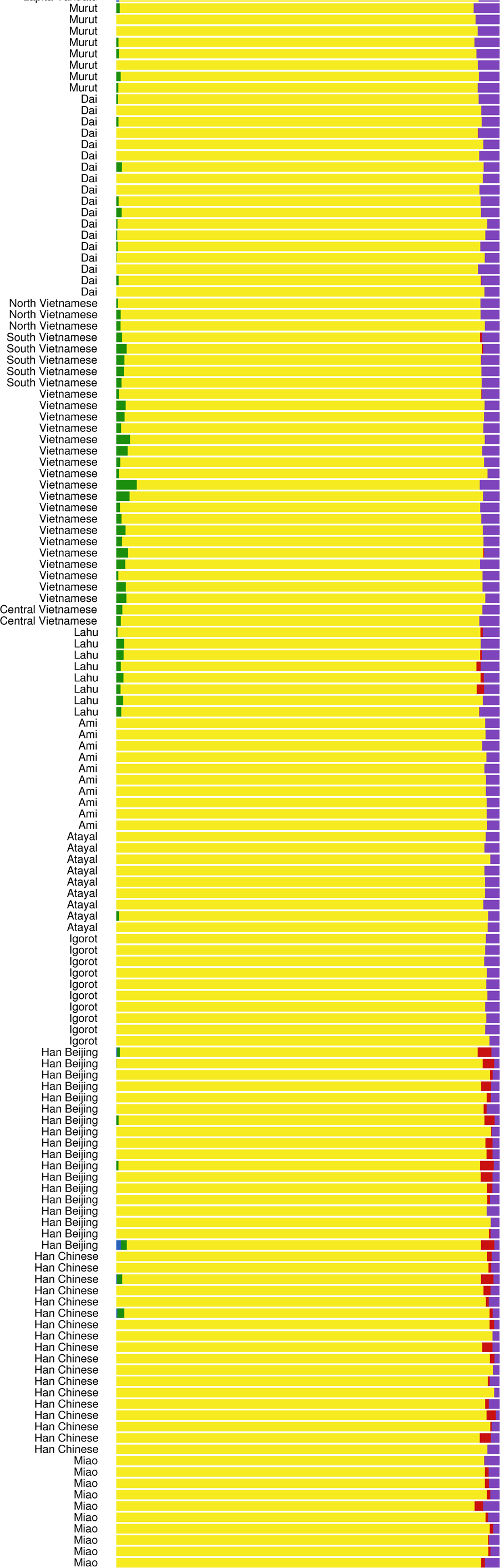
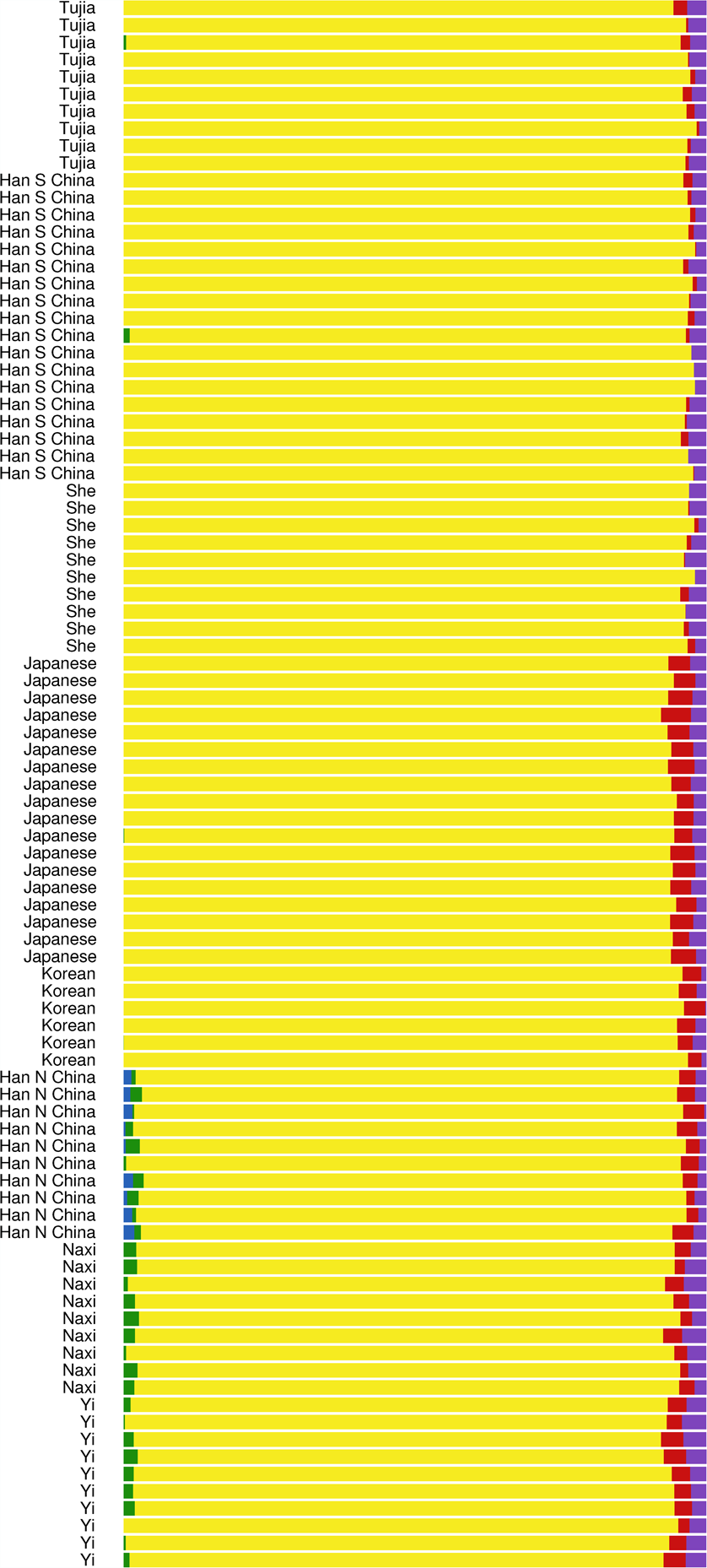
**K = 6 ADMIXTURE analysis**

**Figure 6:**
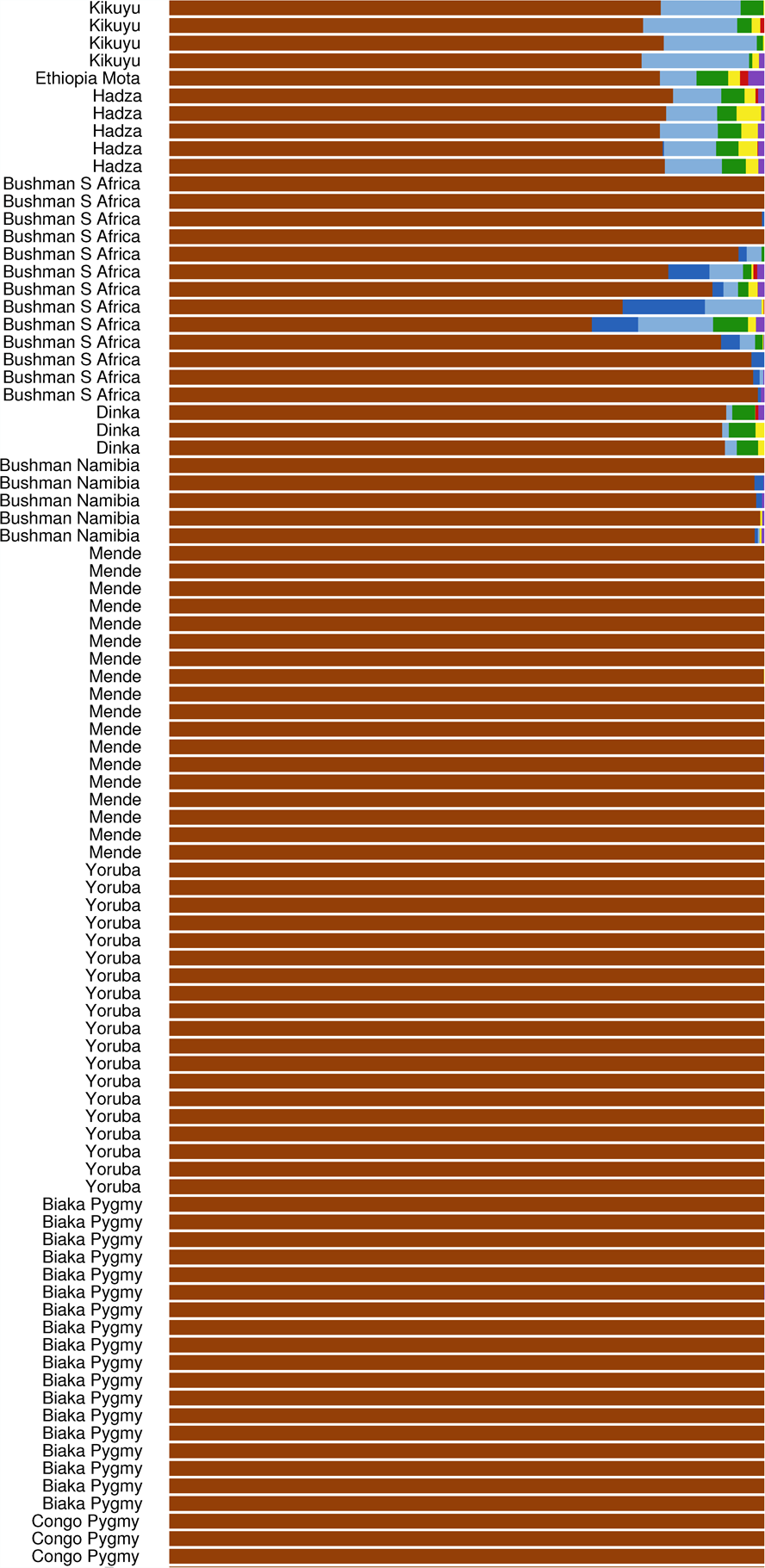
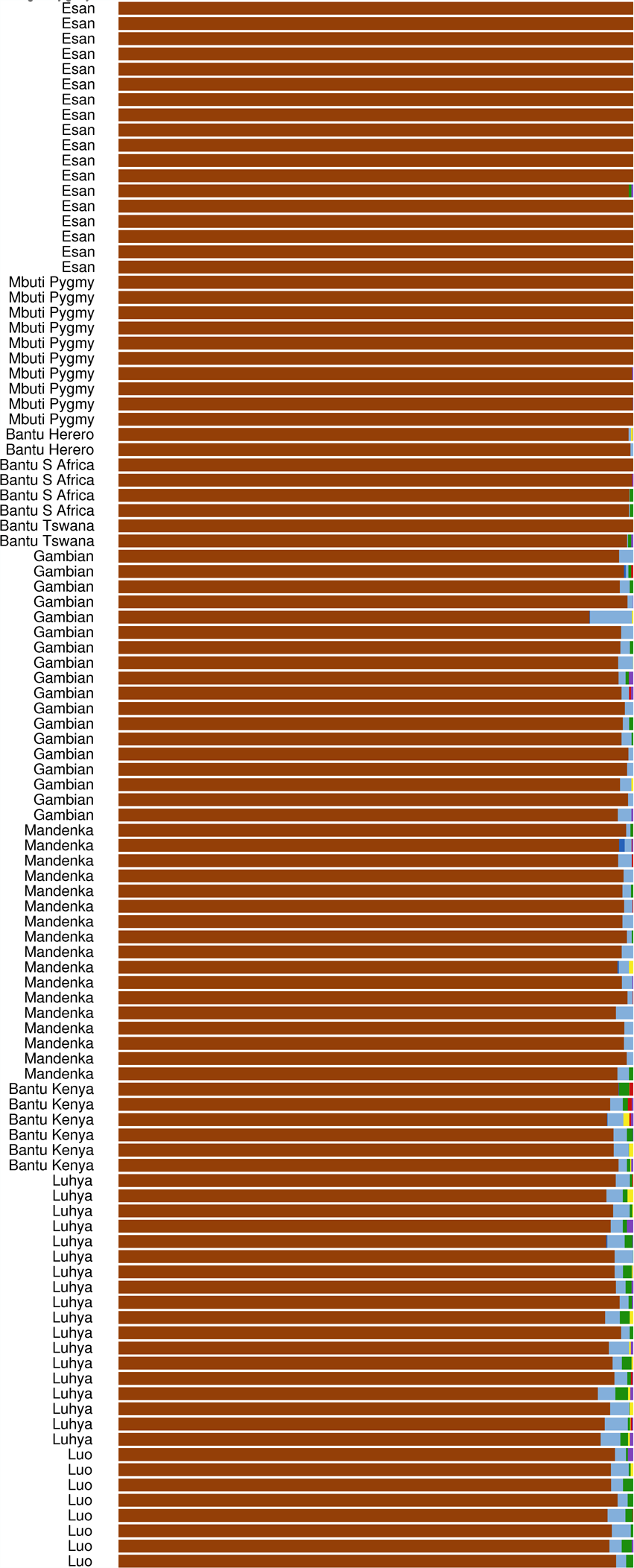
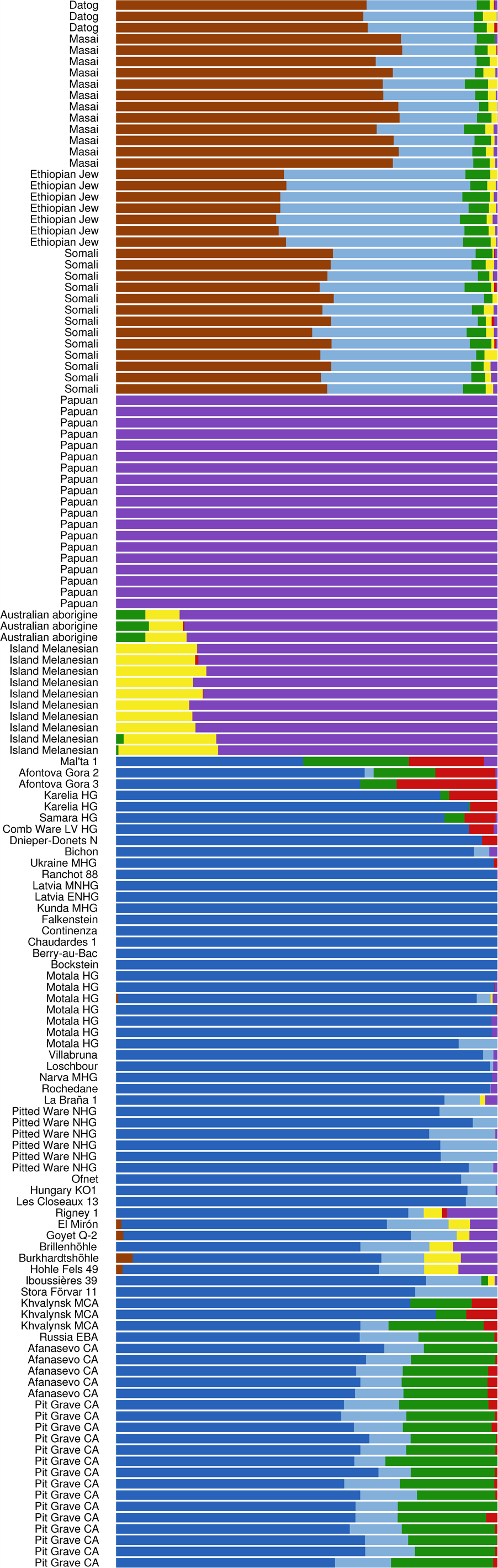
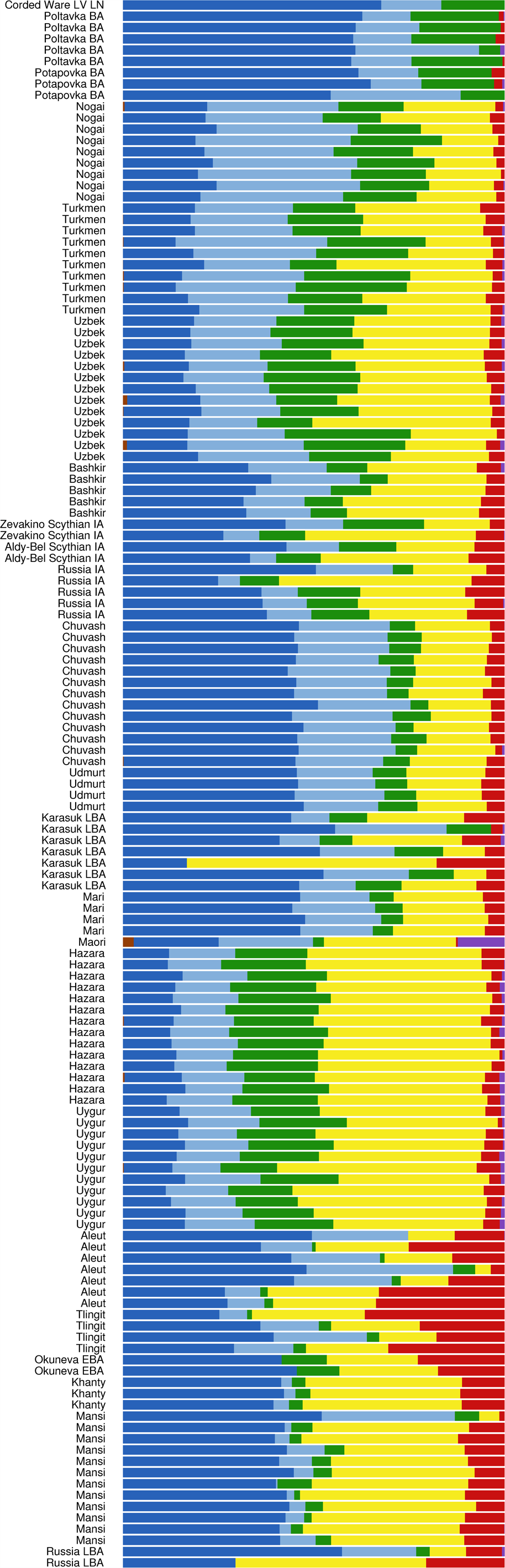
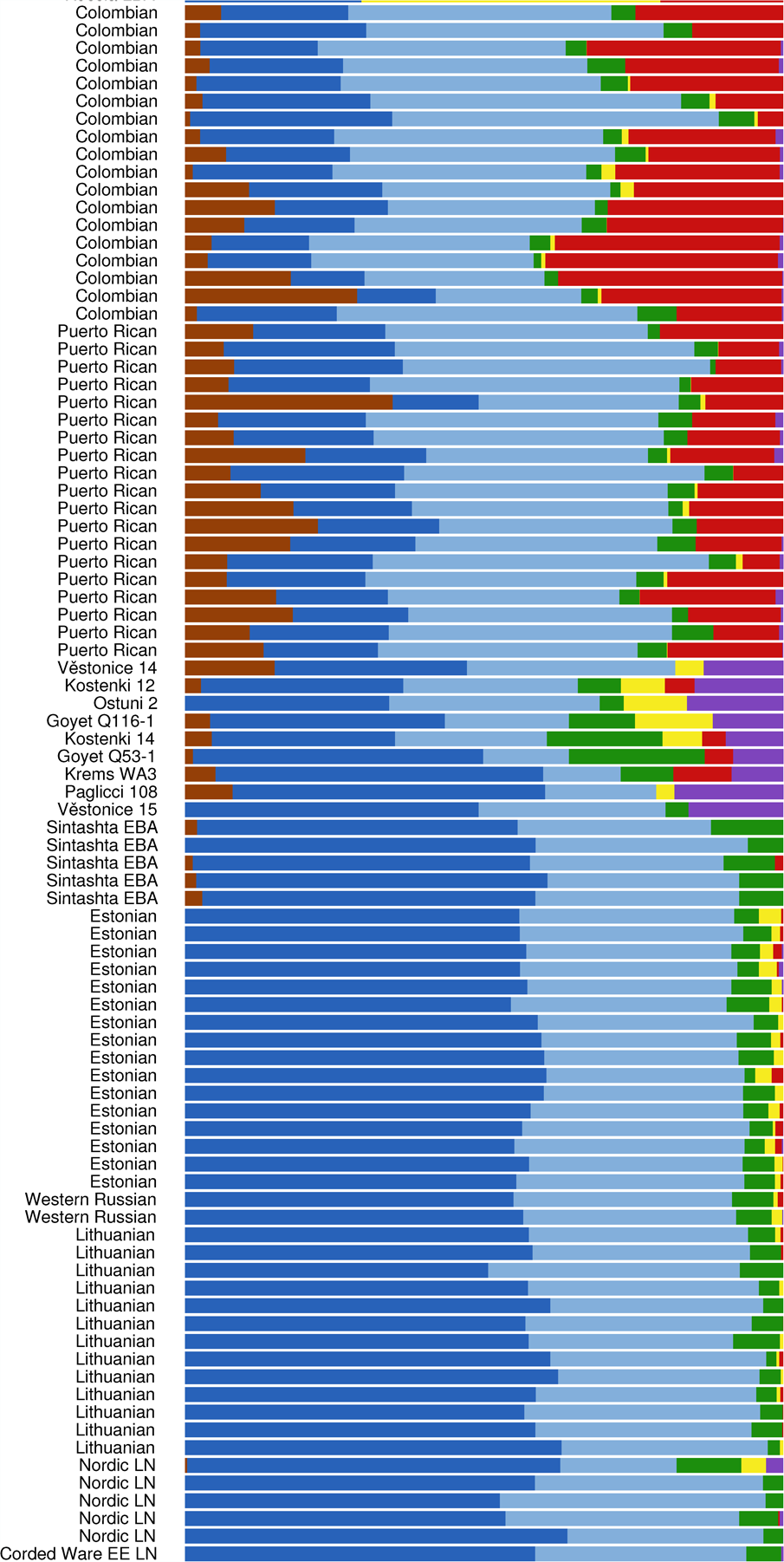
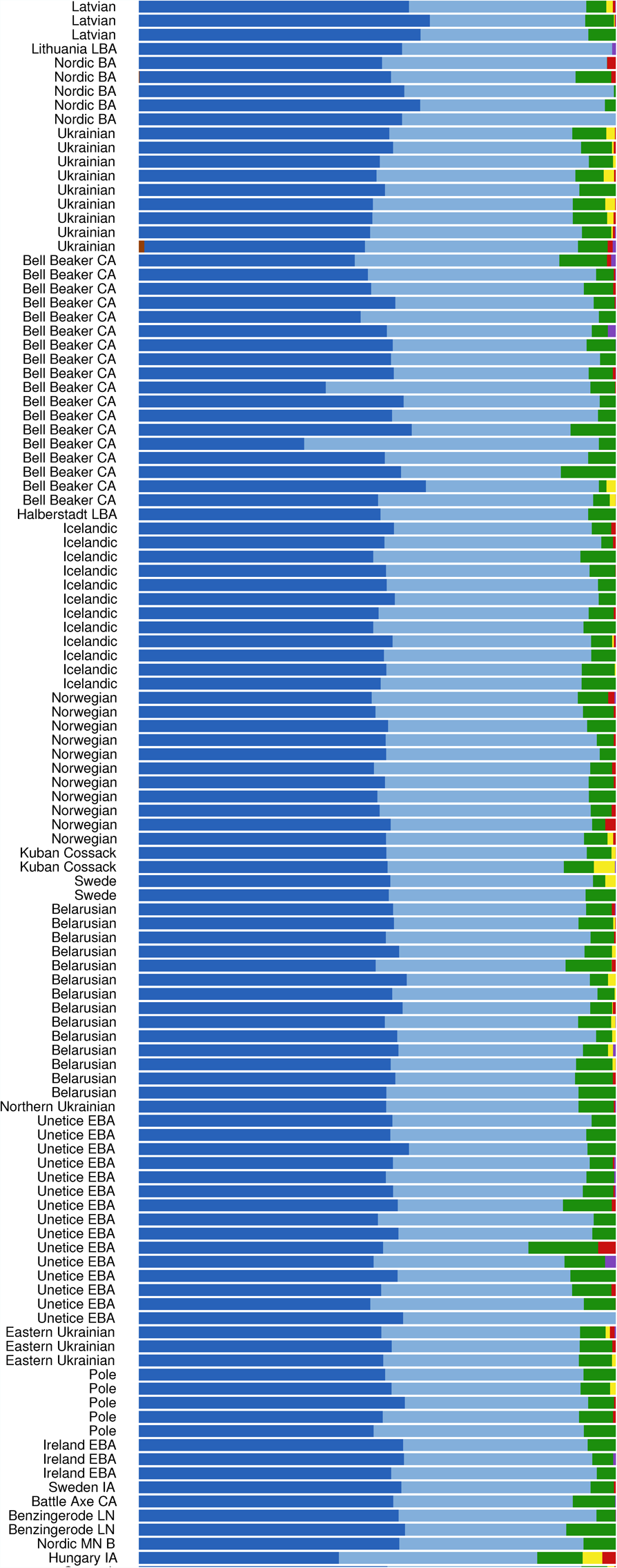
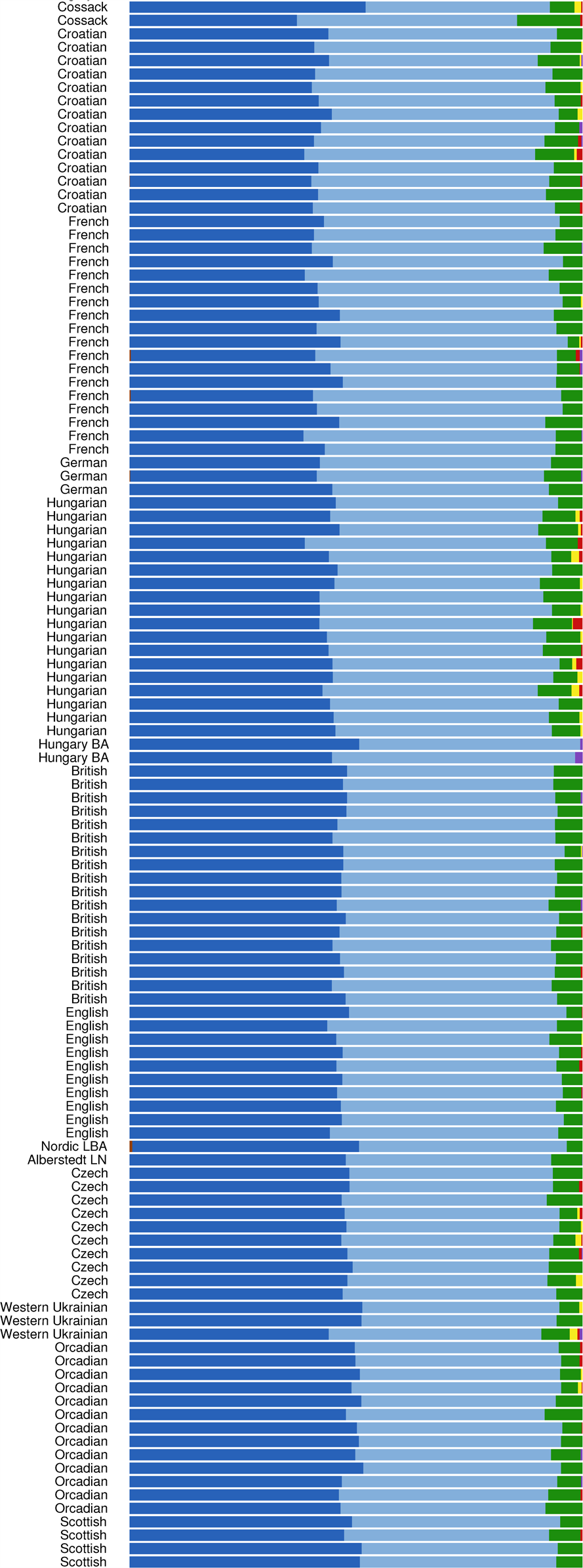
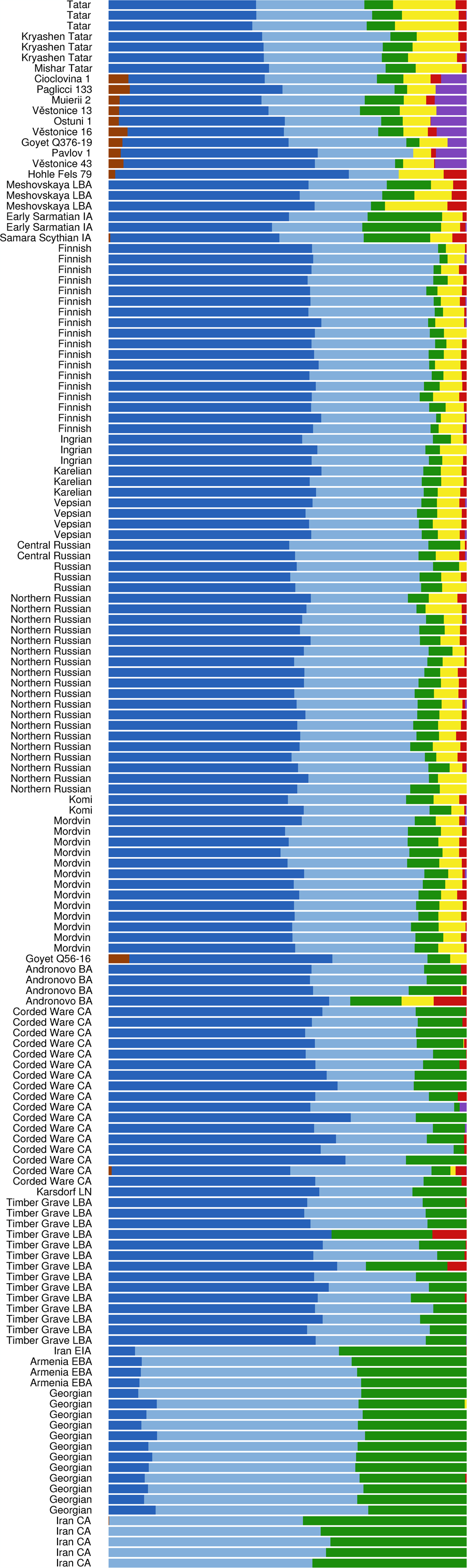
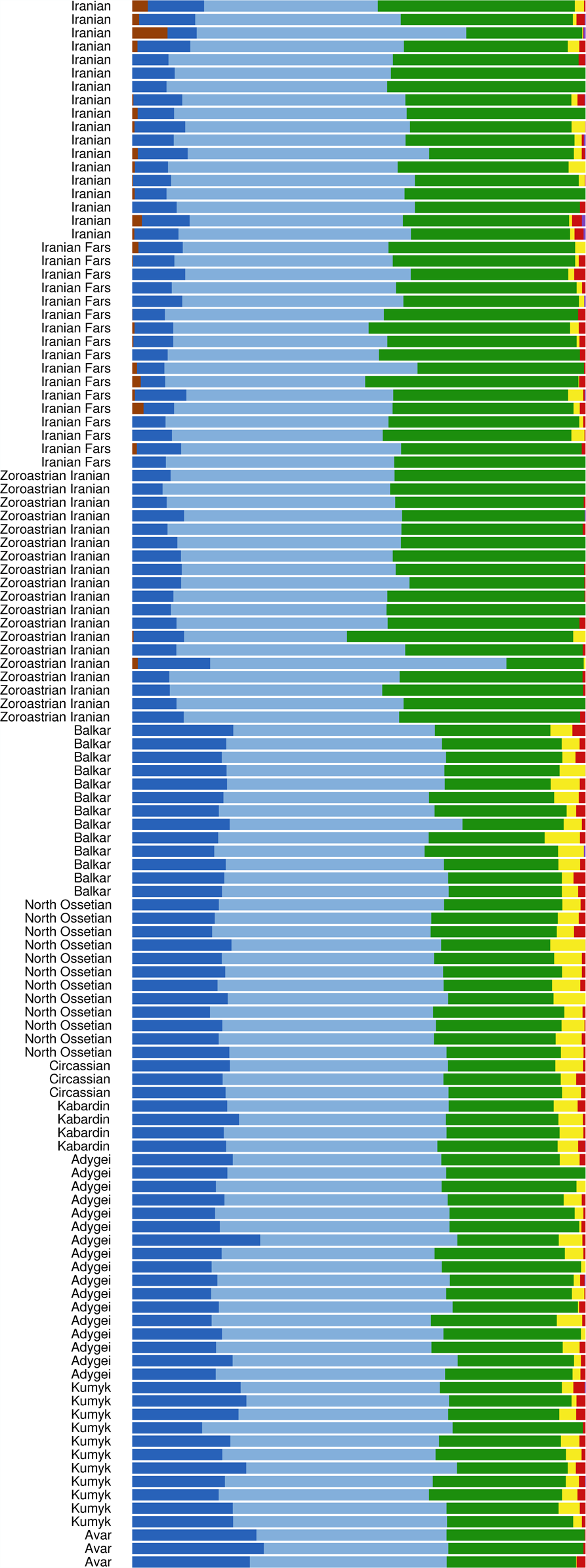
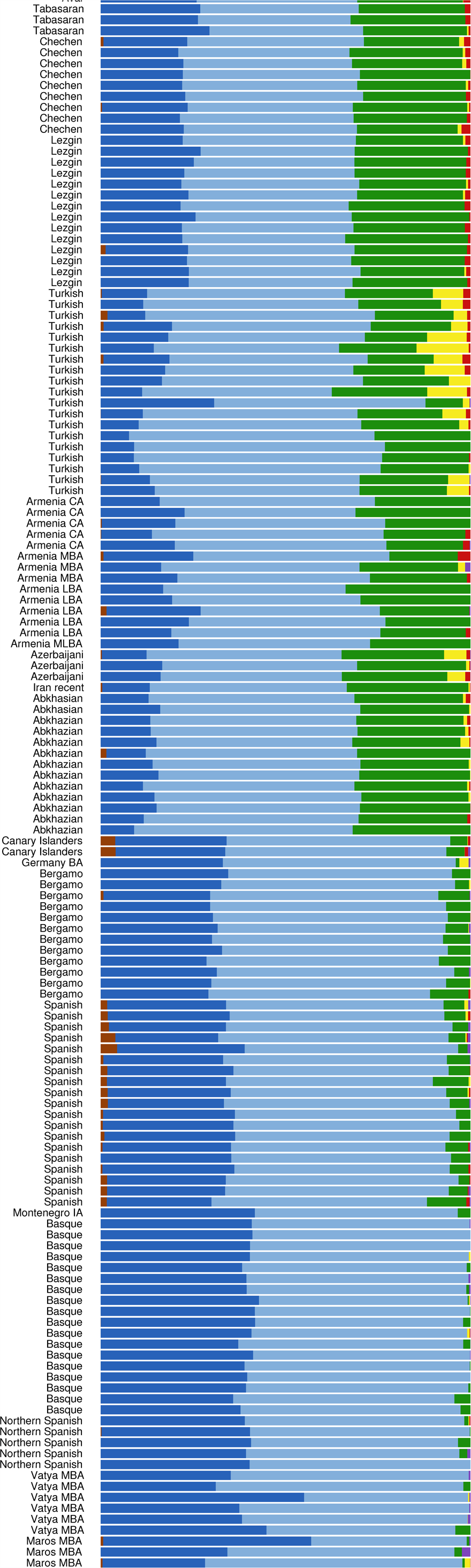
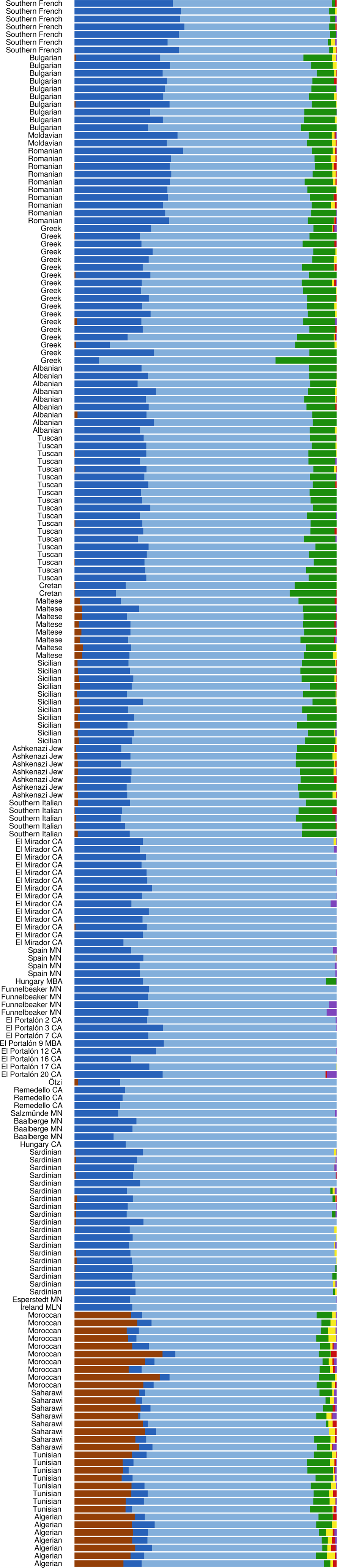
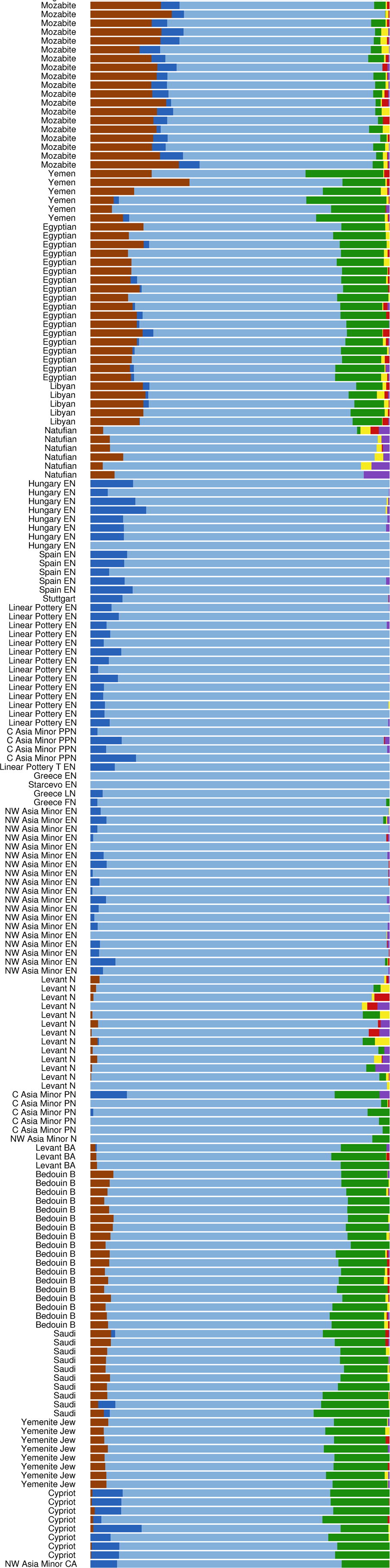
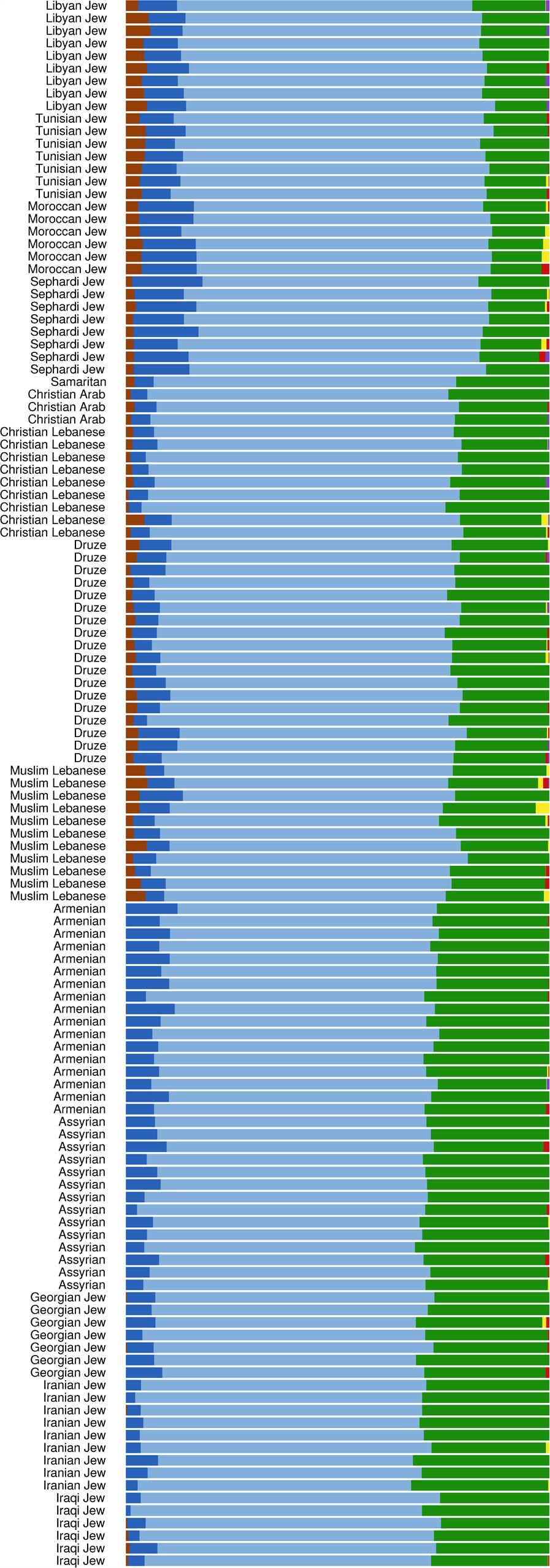
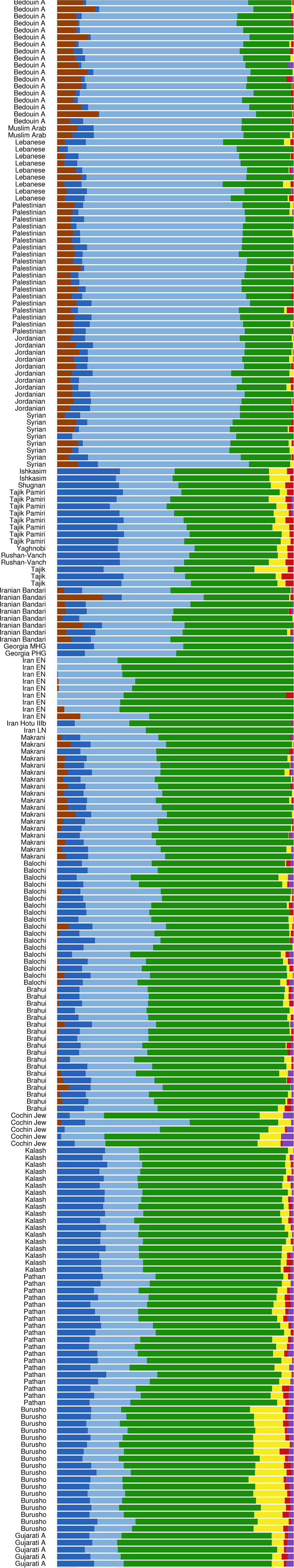
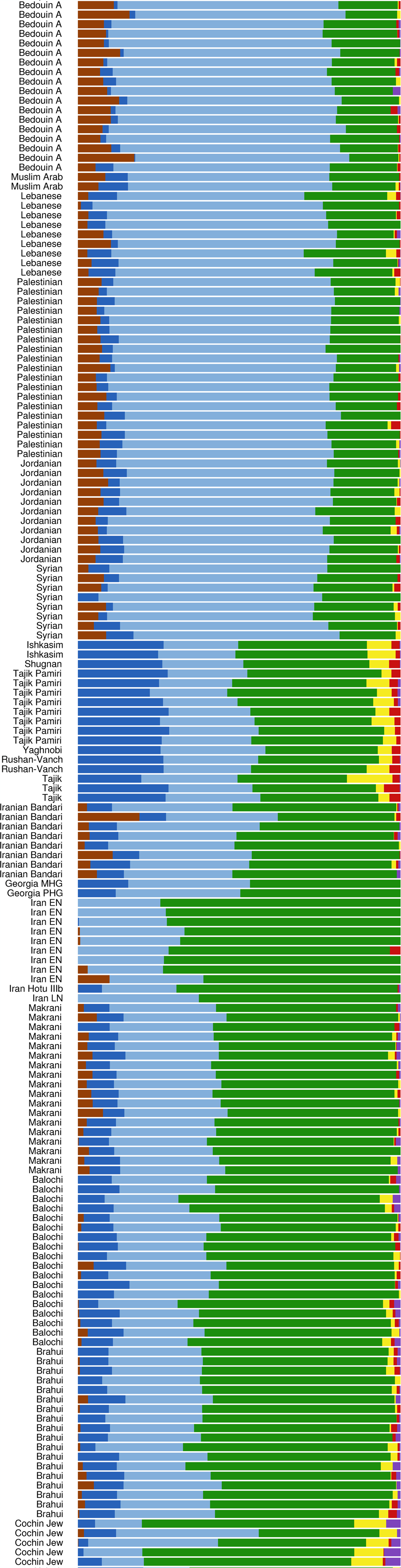
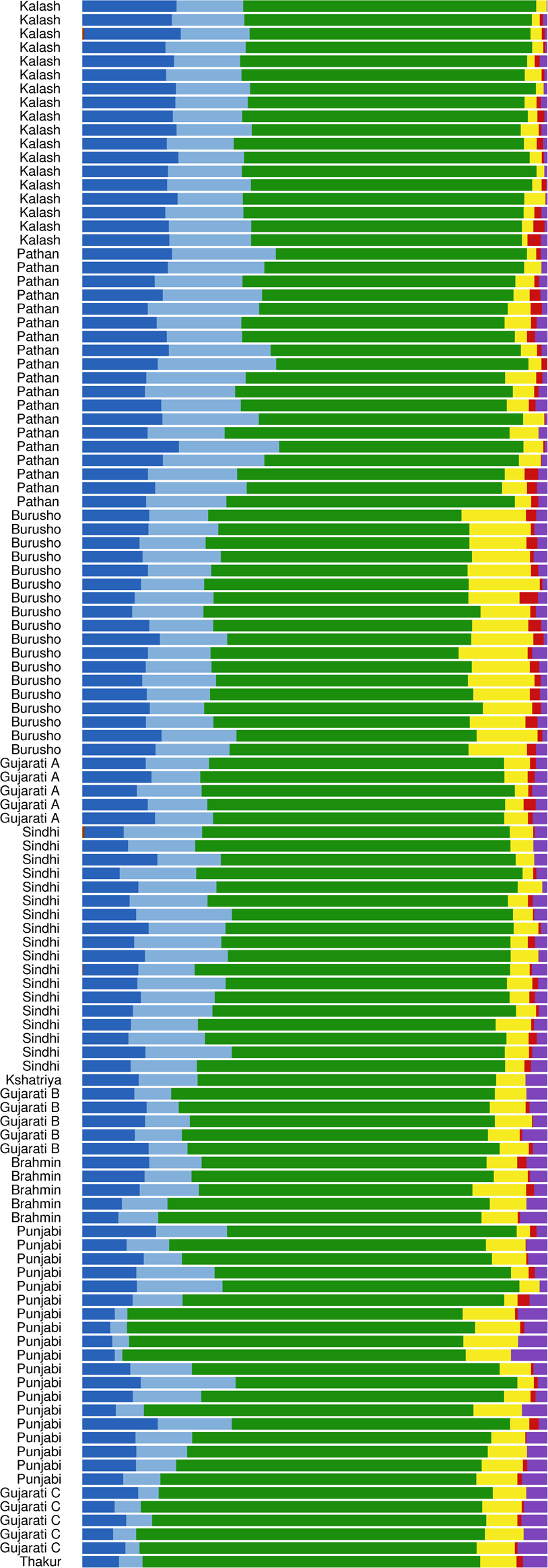
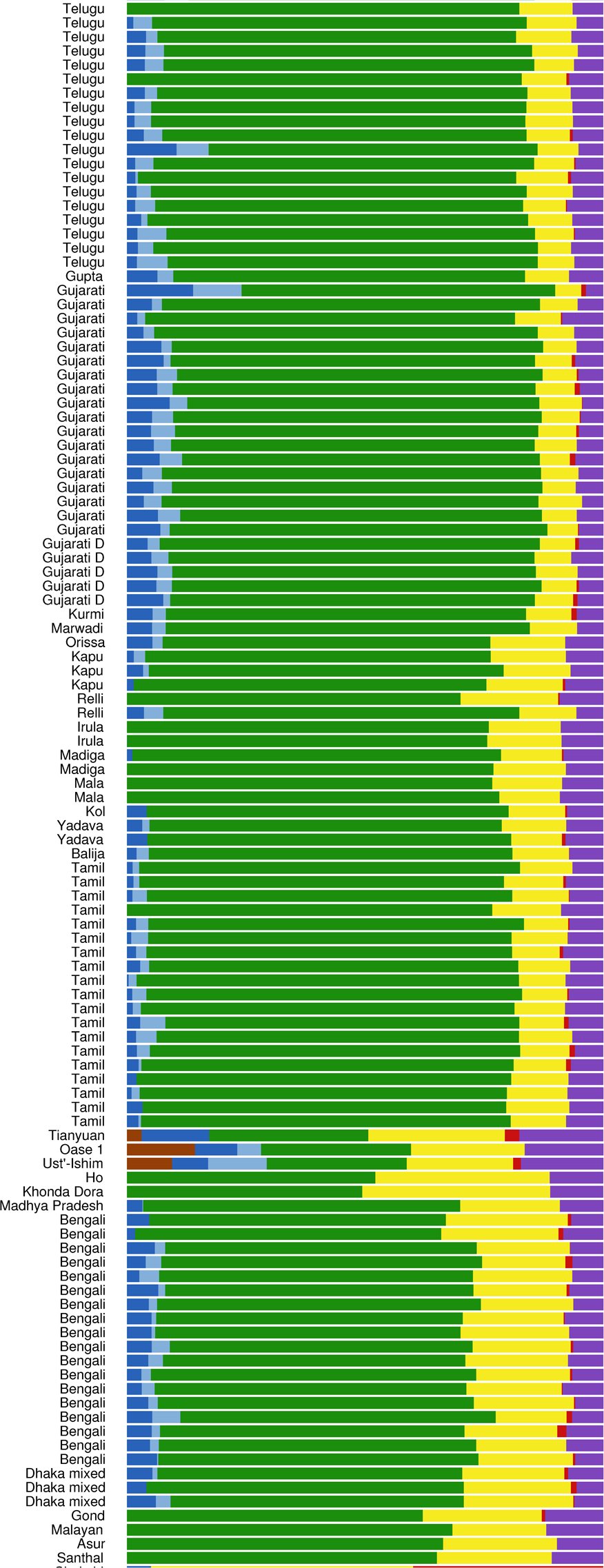
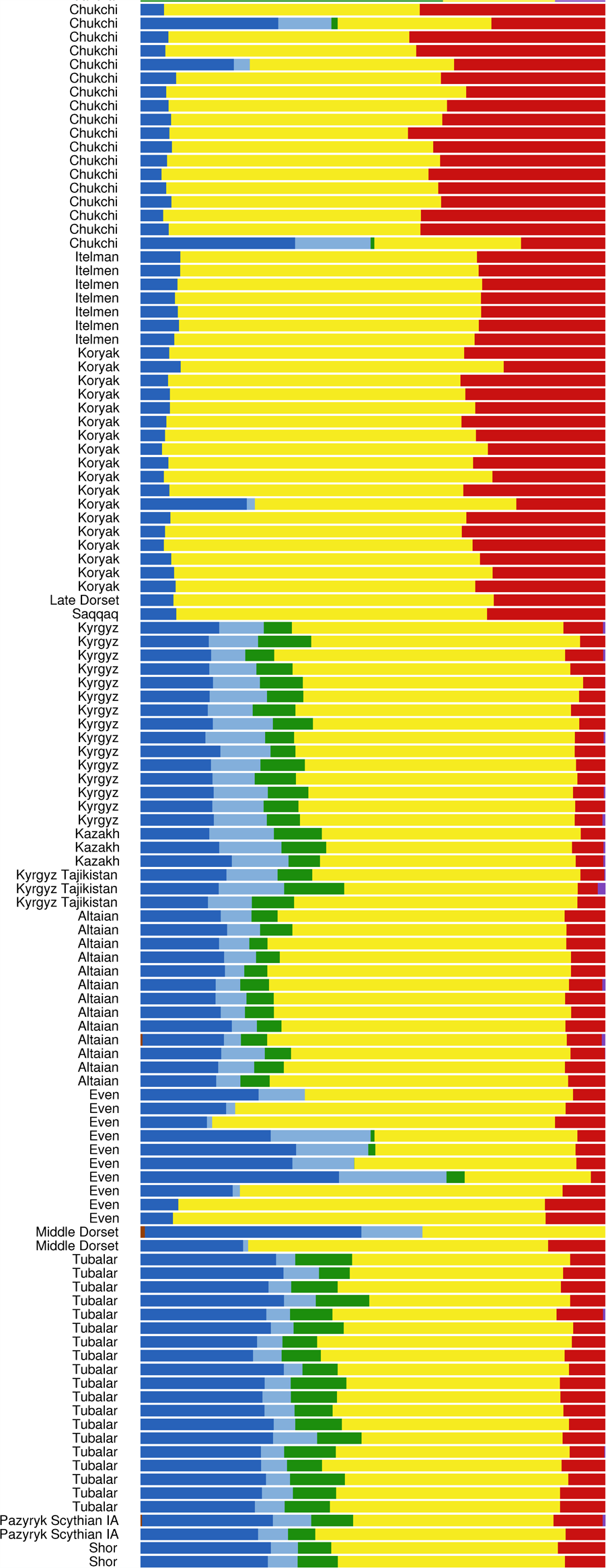
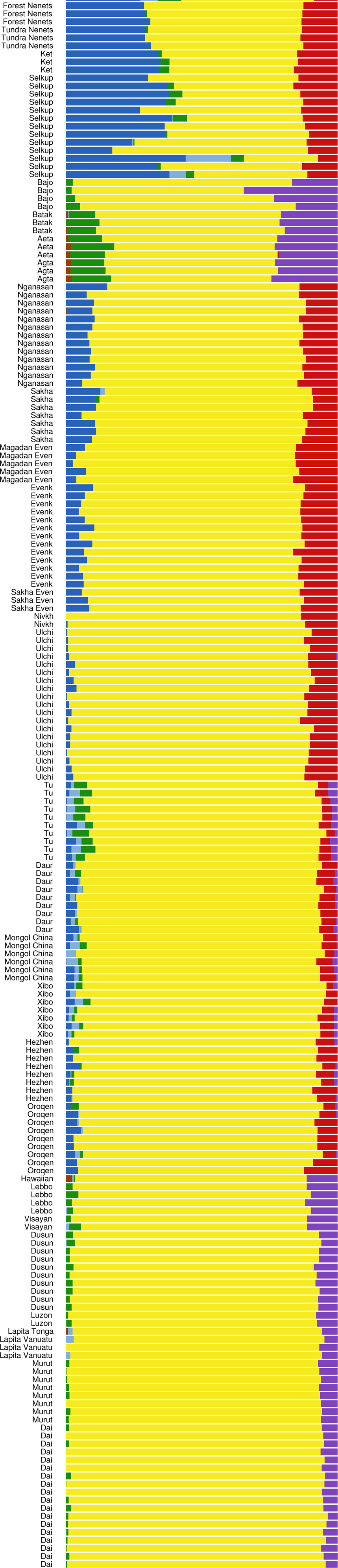
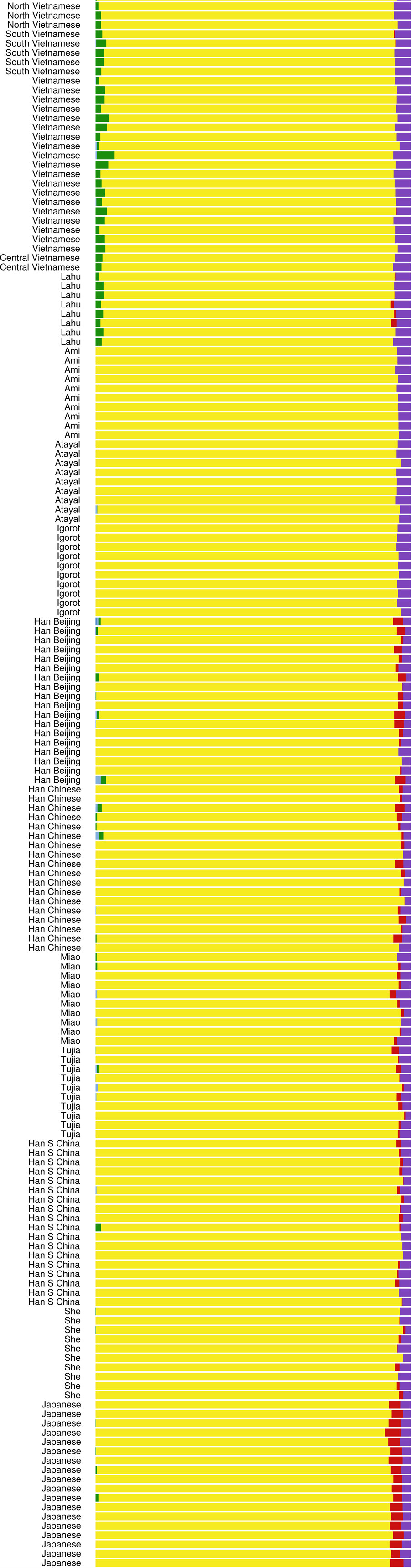
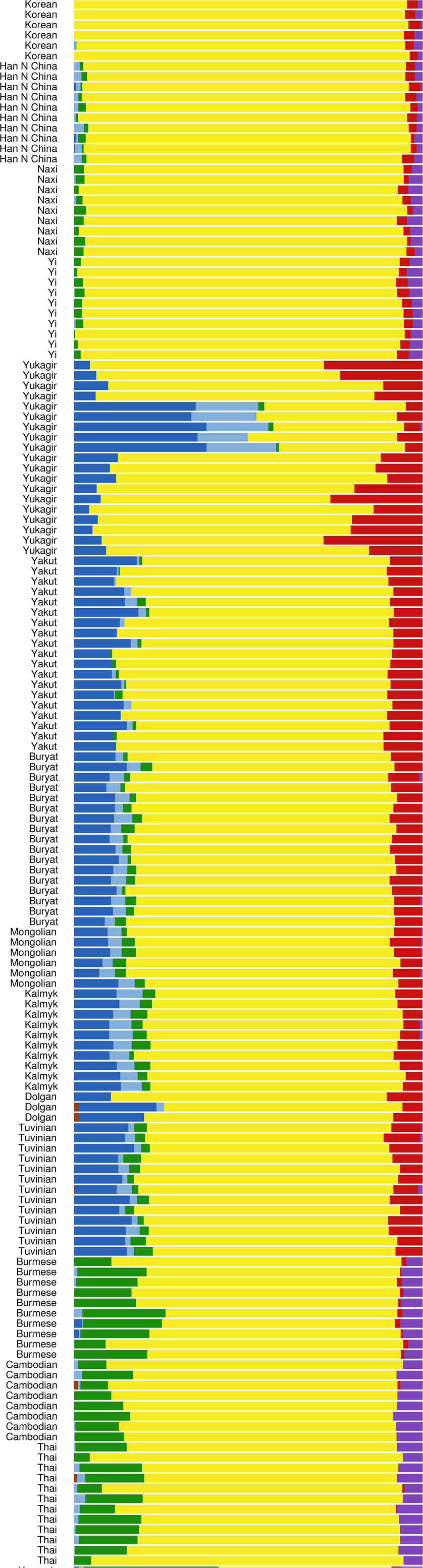
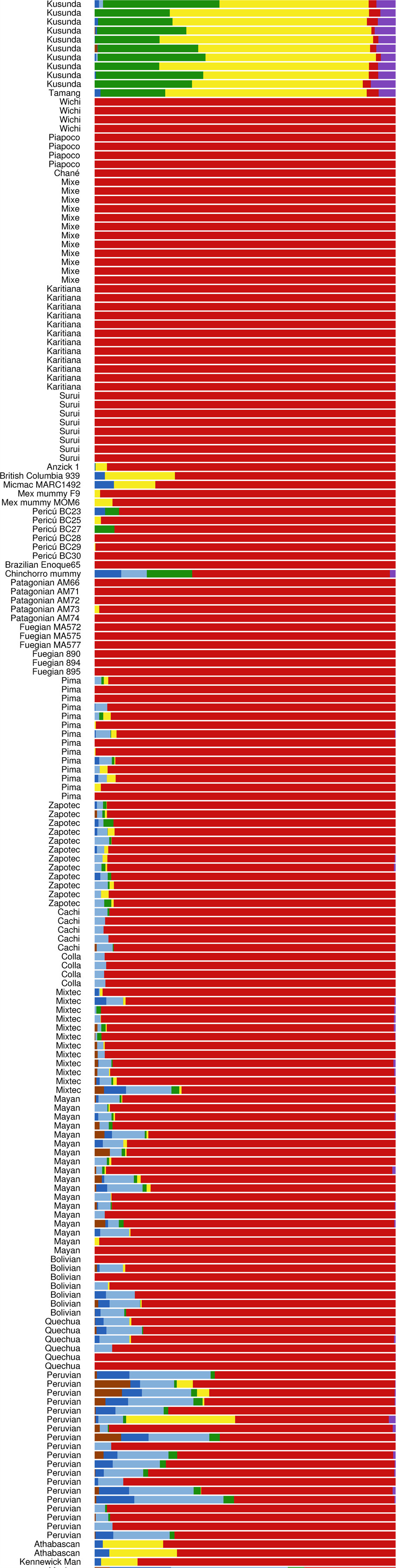
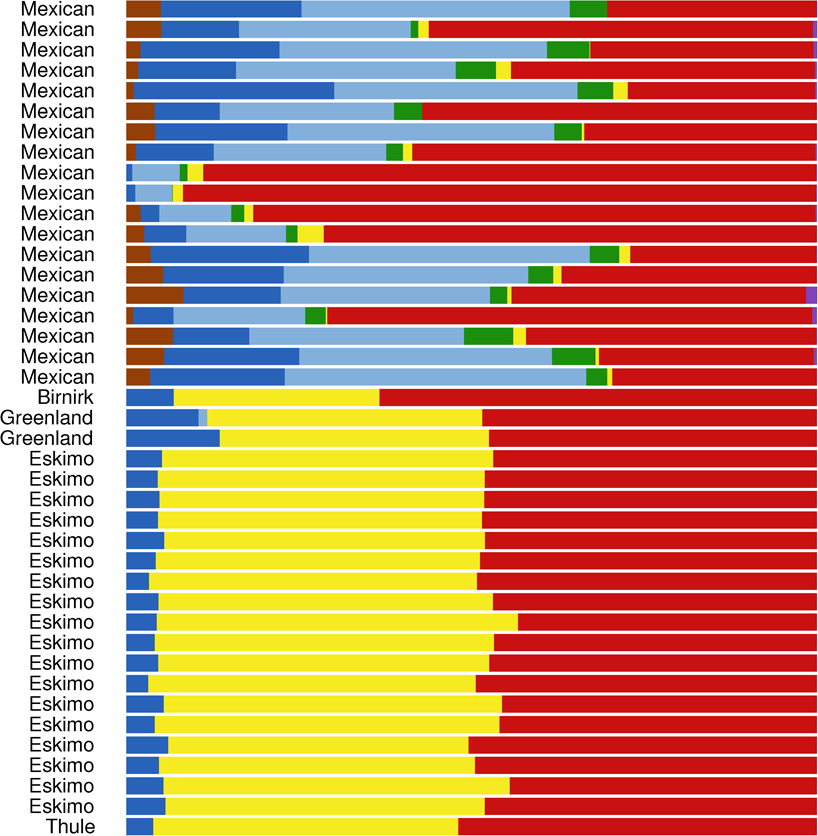
**K = 7 ADMIXTURE analysis**

**Figure 7:**
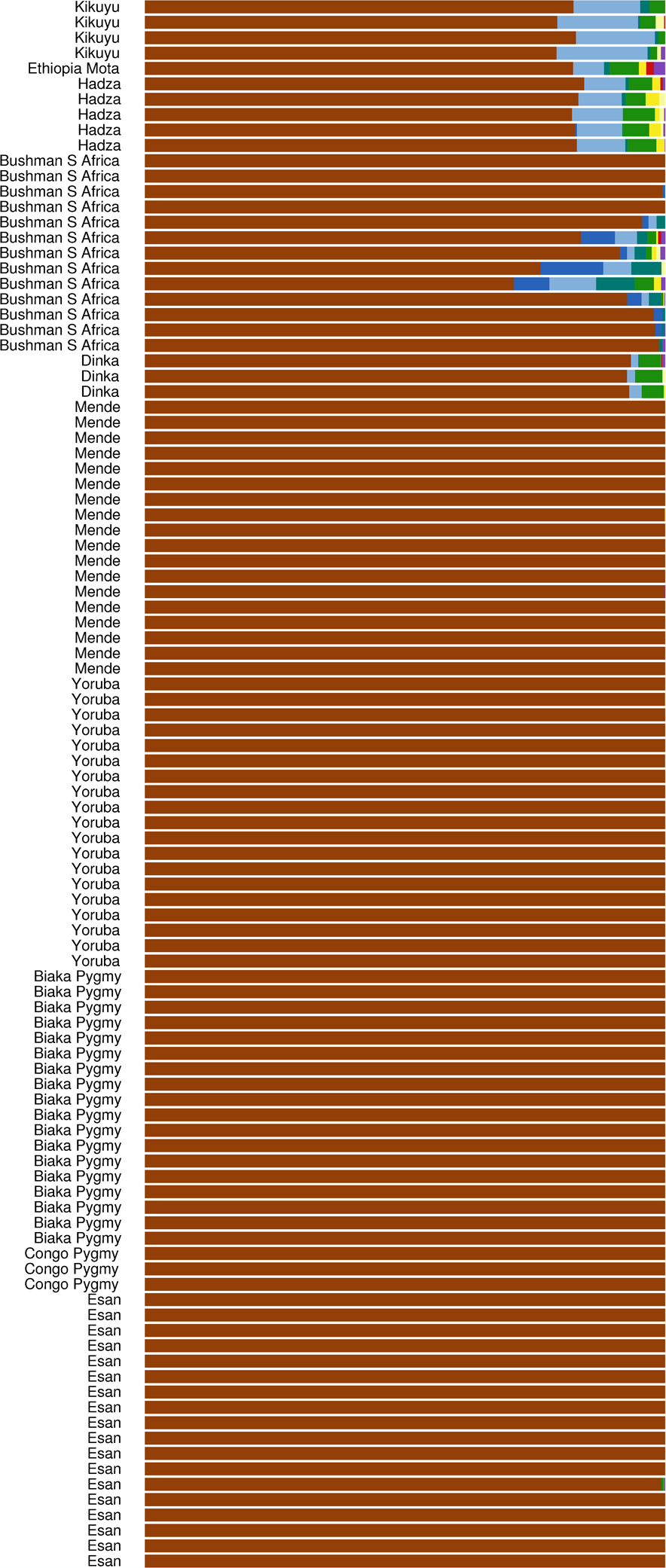
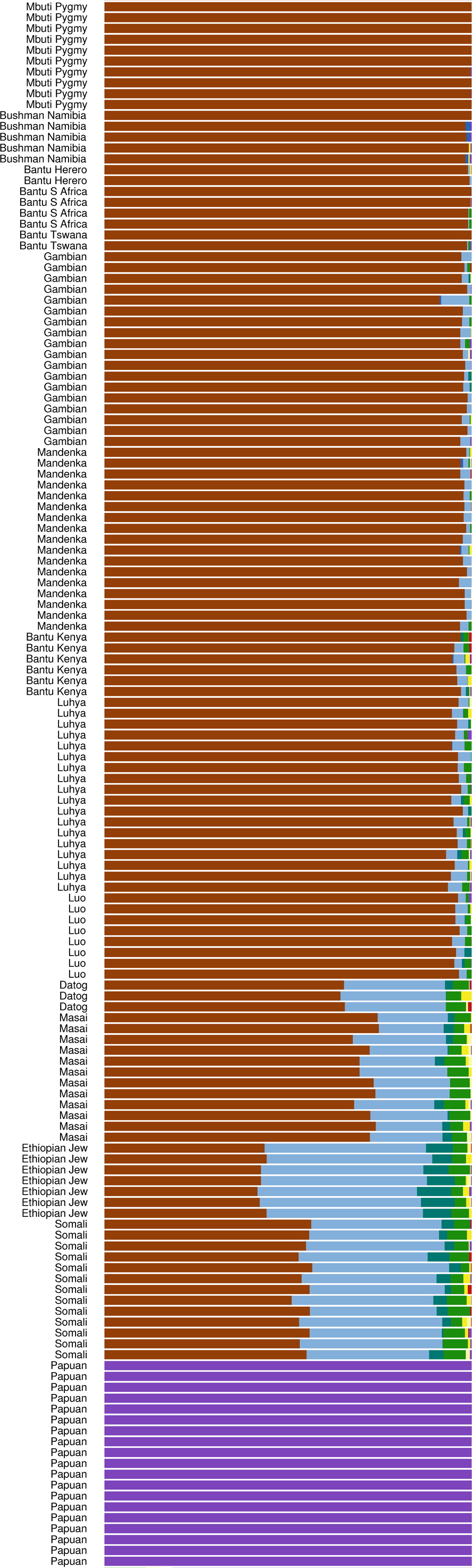
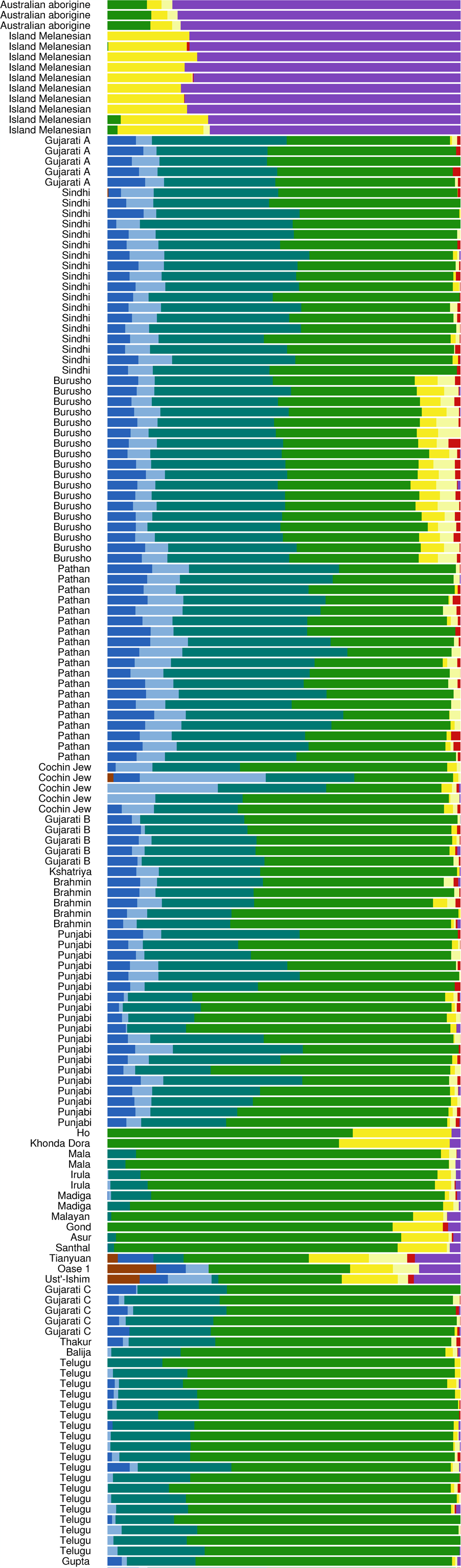
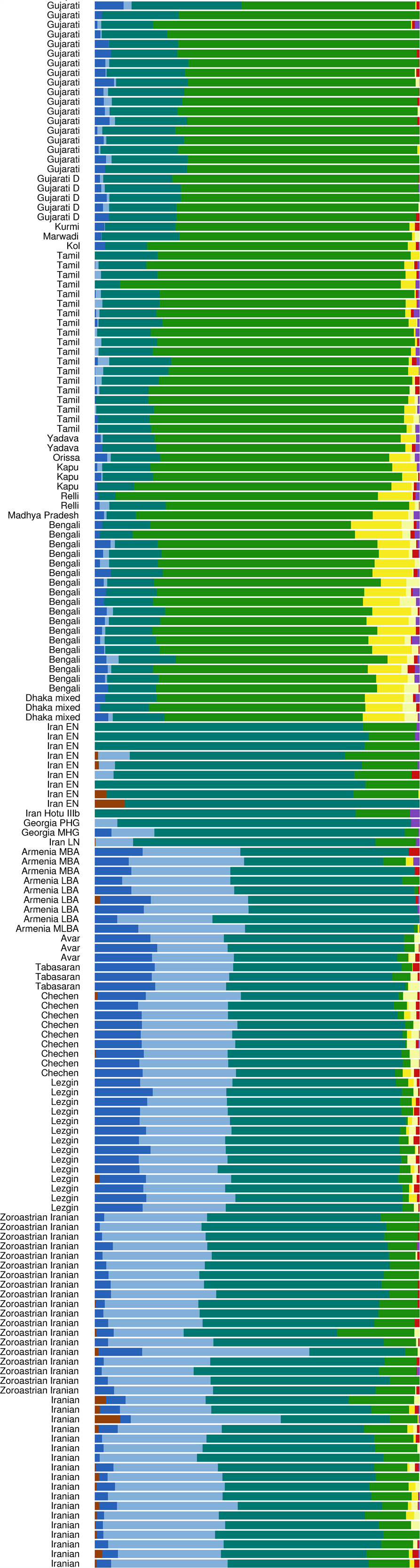
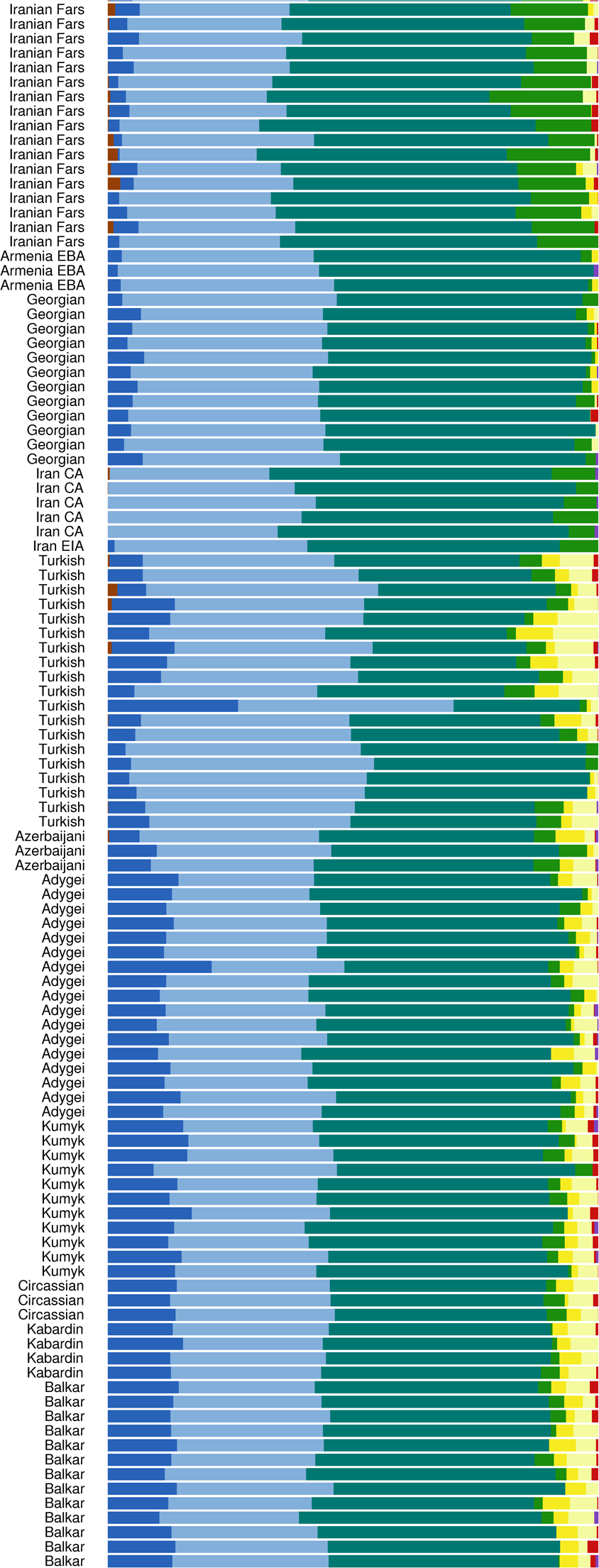
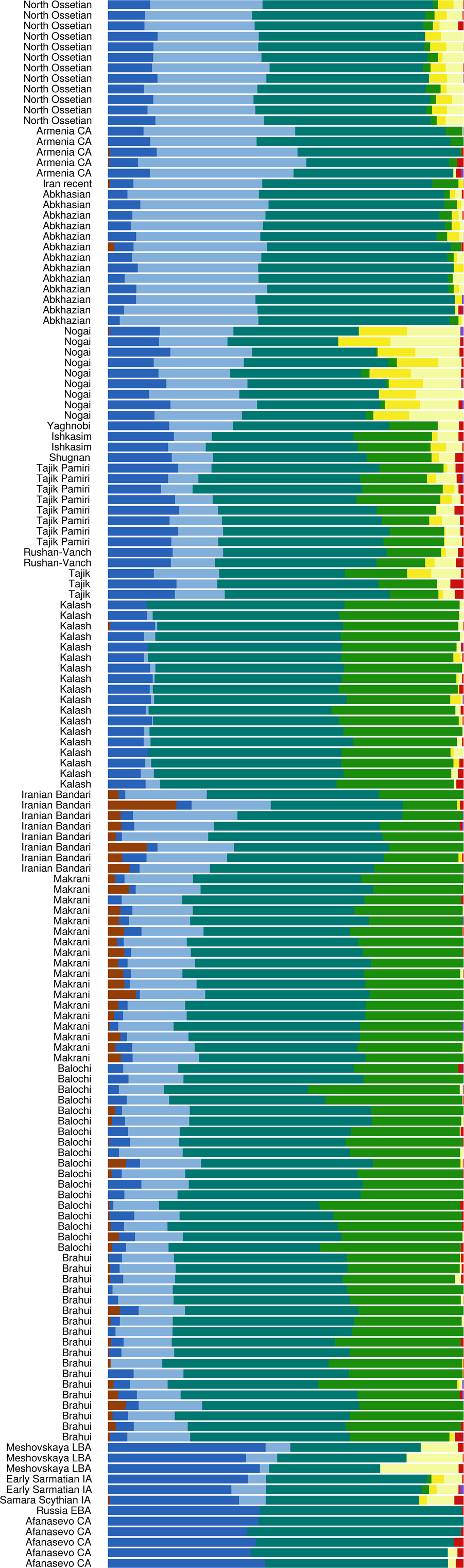
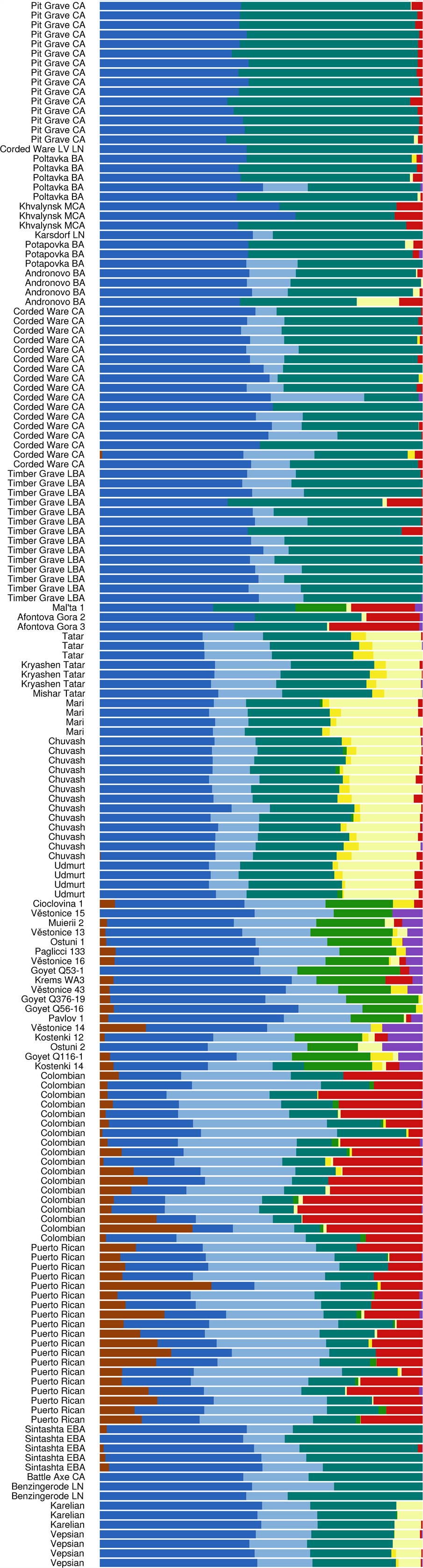
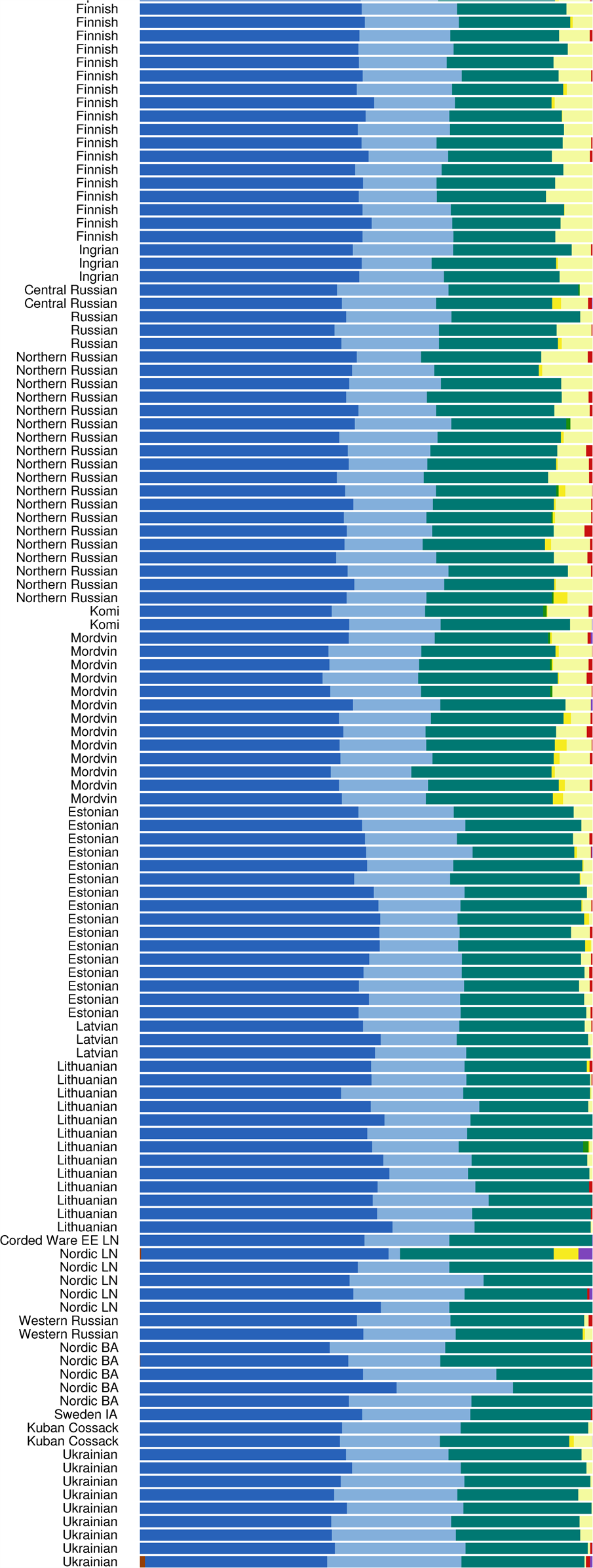
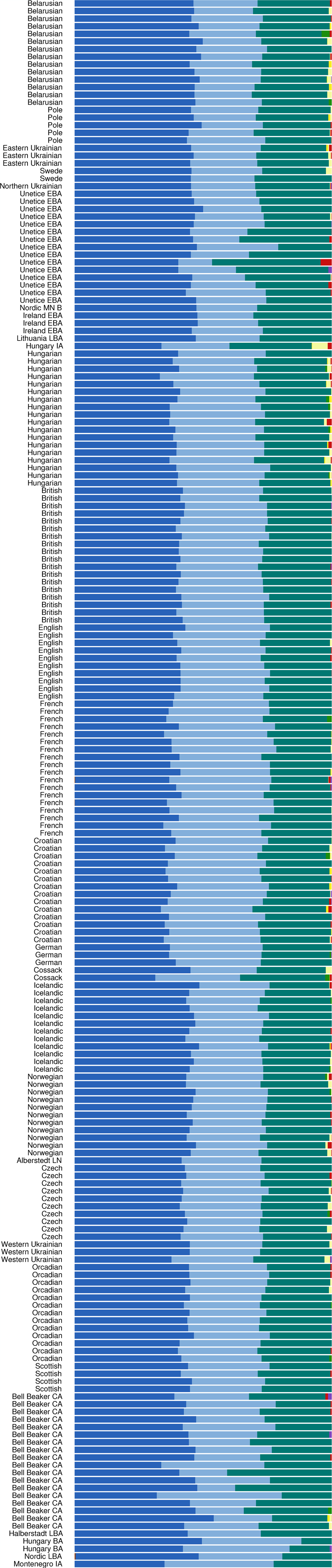
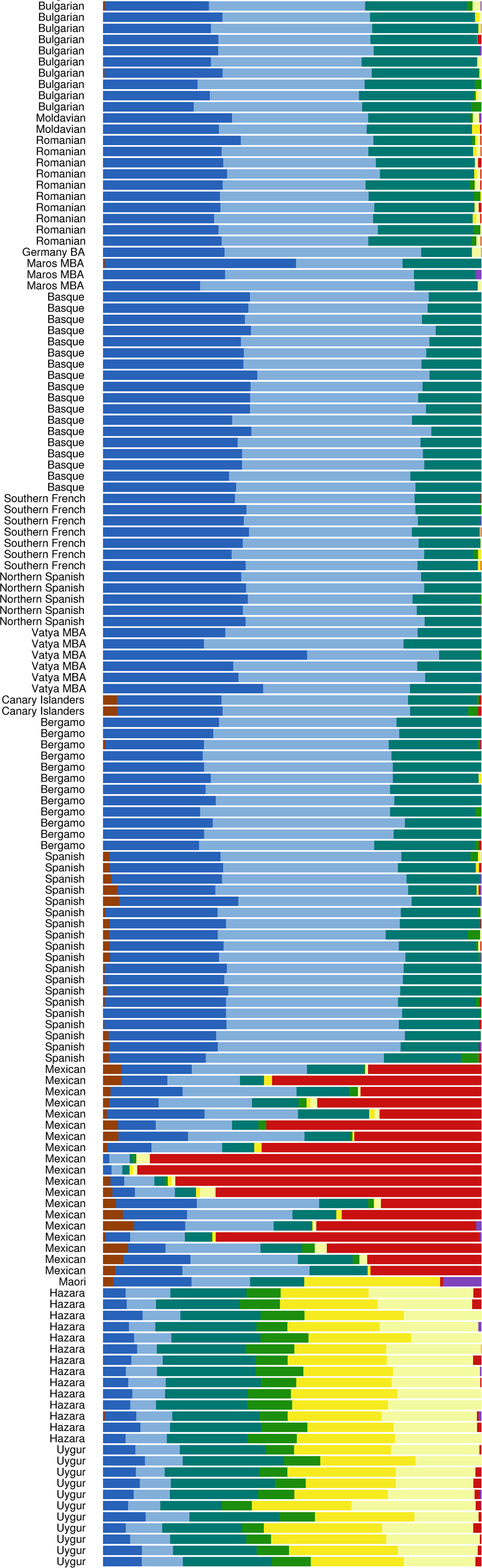
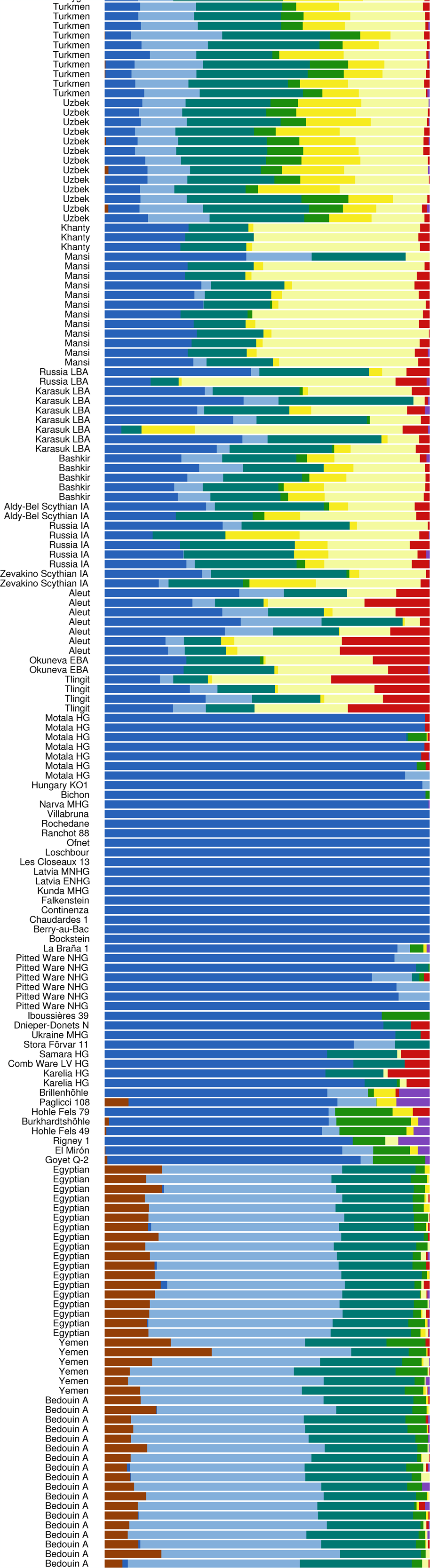
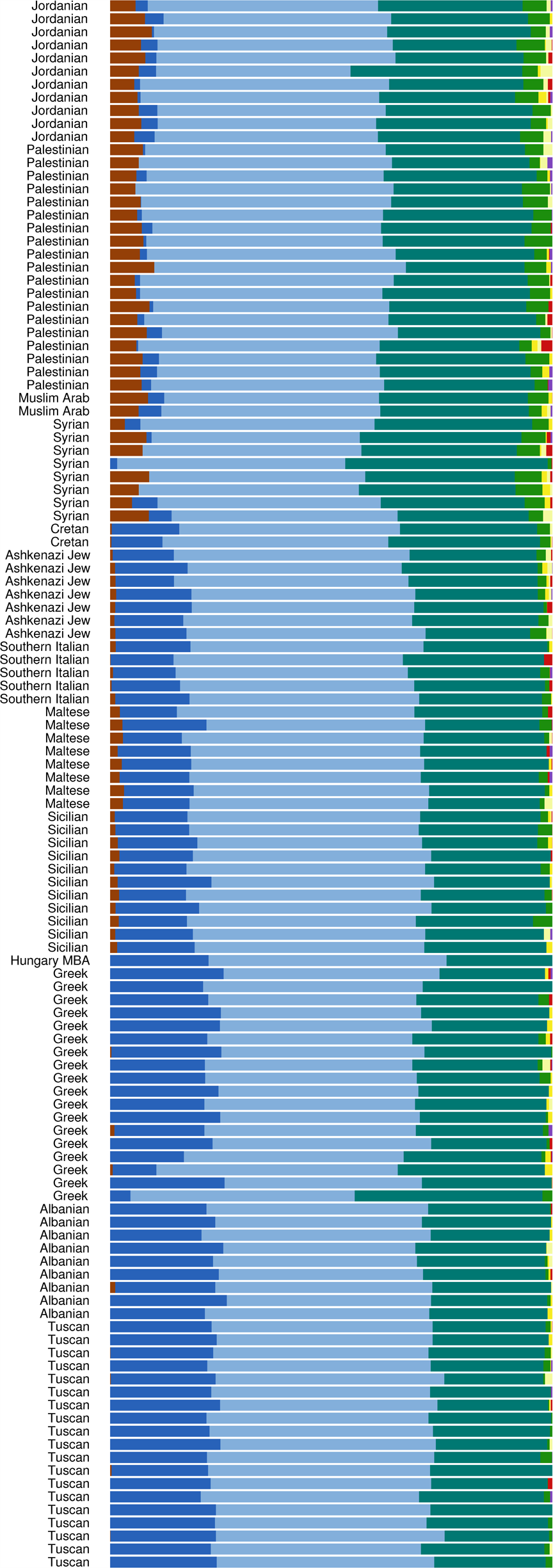
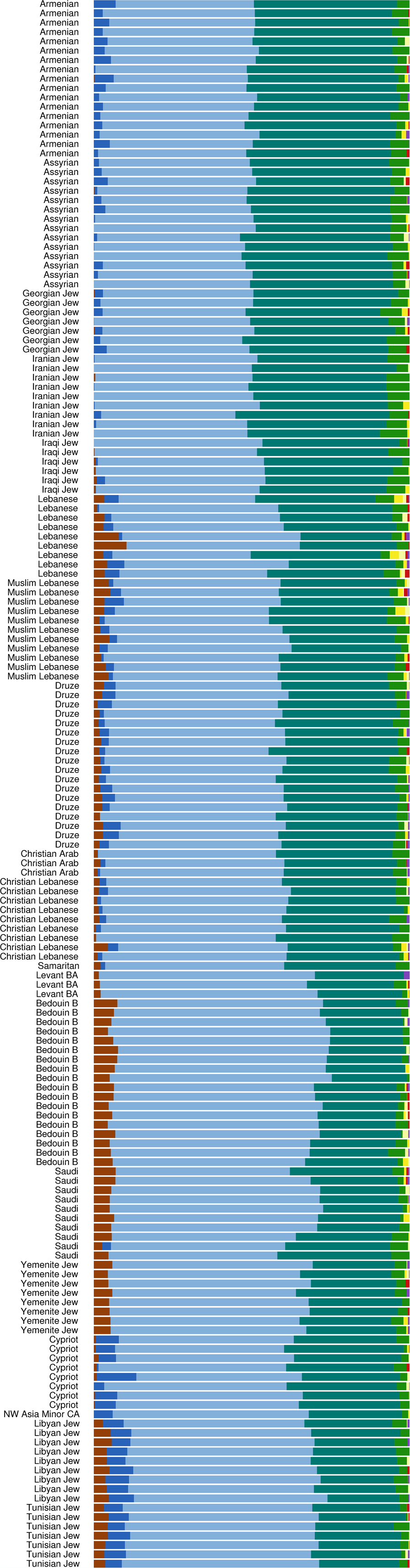
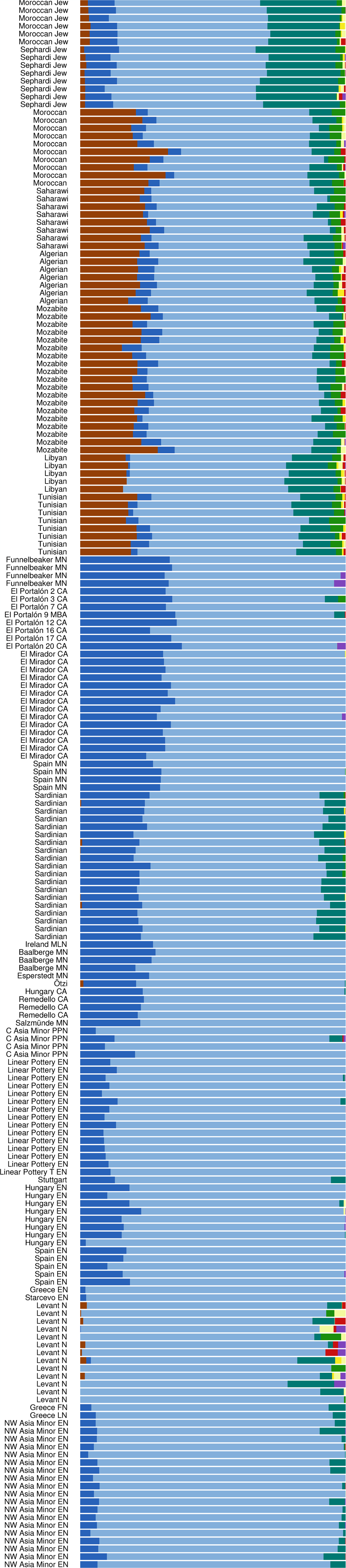
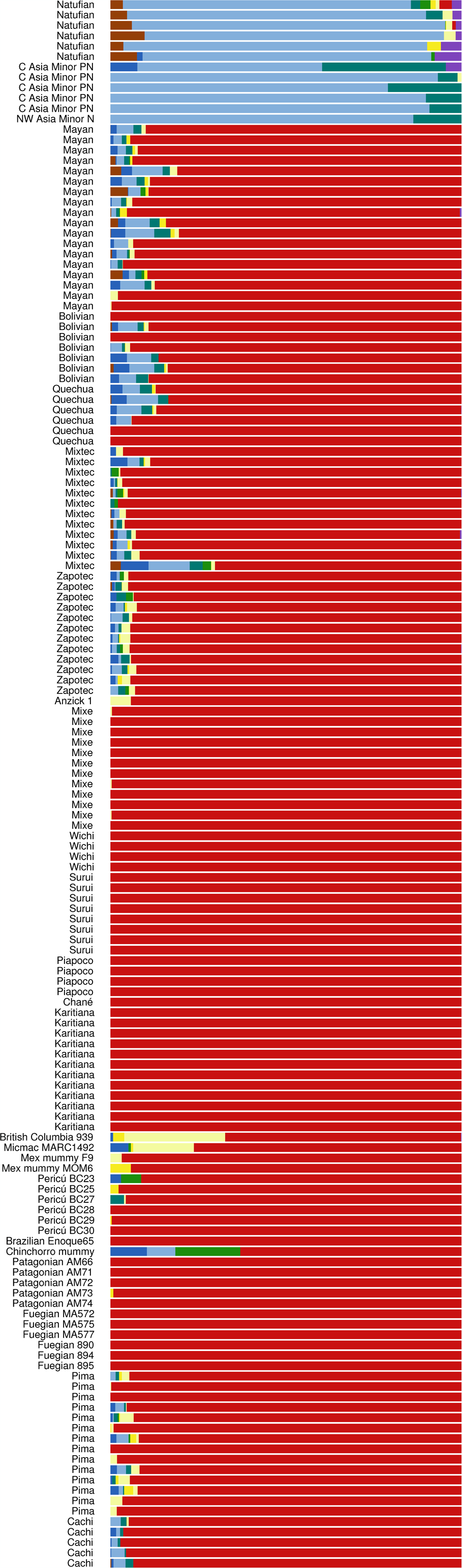
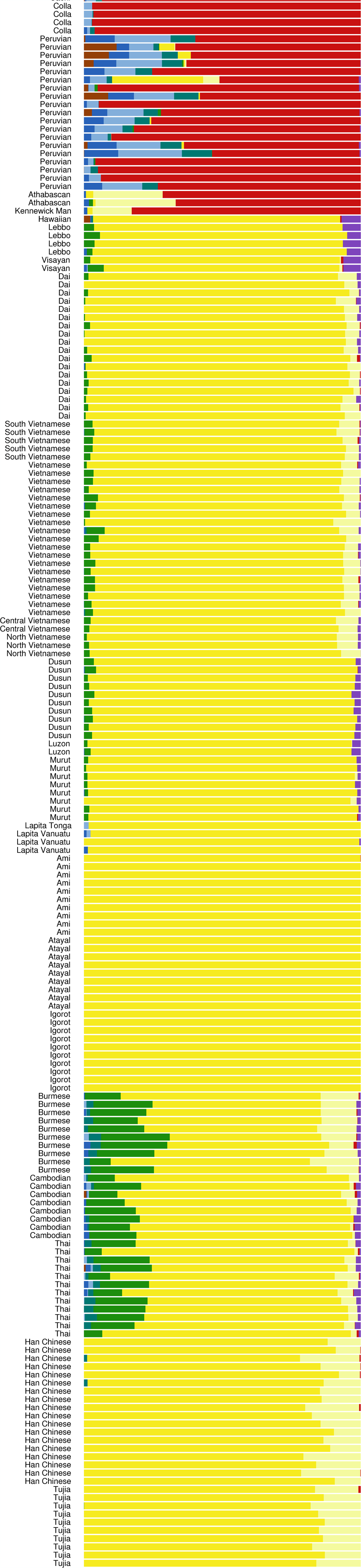
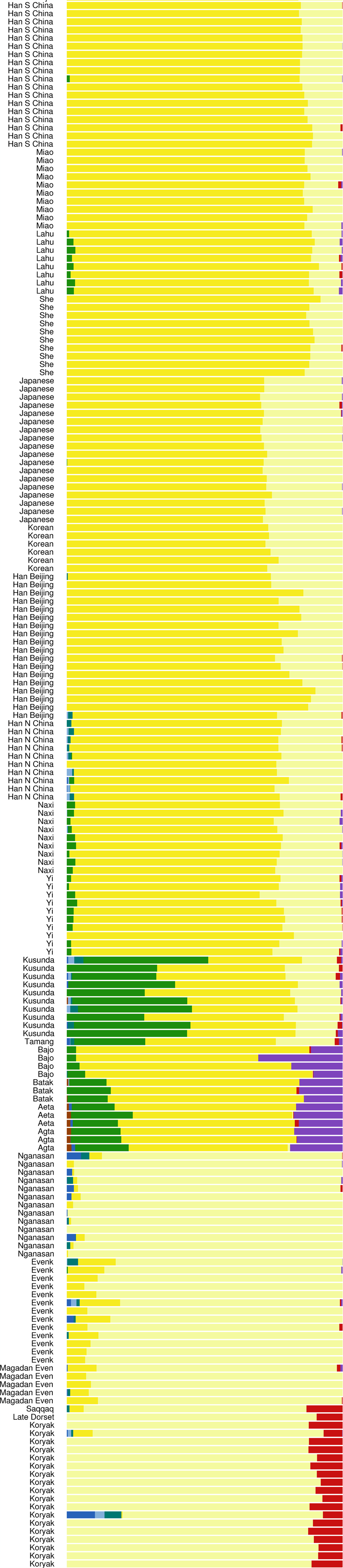
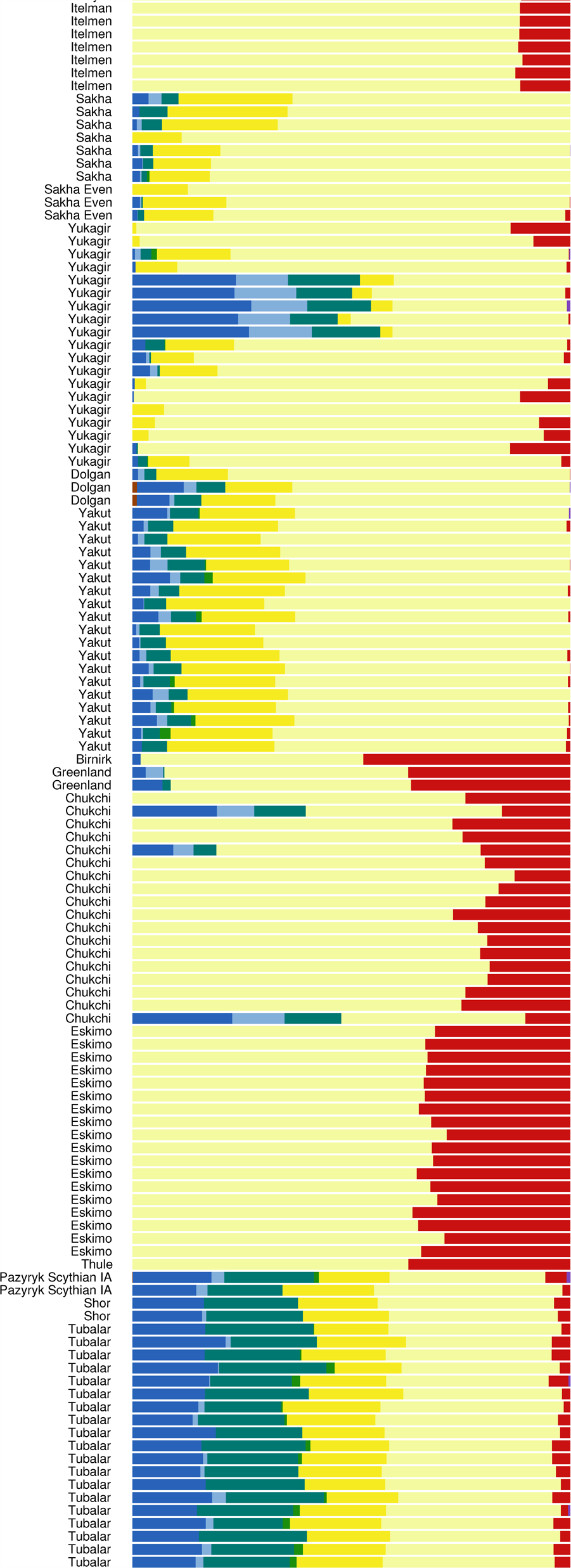
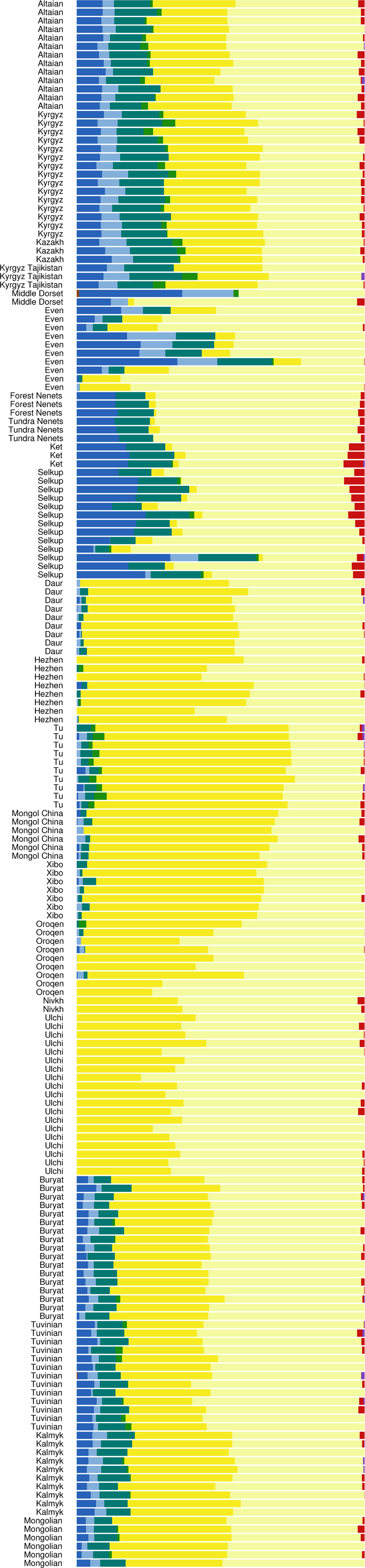
**K = 9 ADMIXTURE analysis**

**Figure 8:**
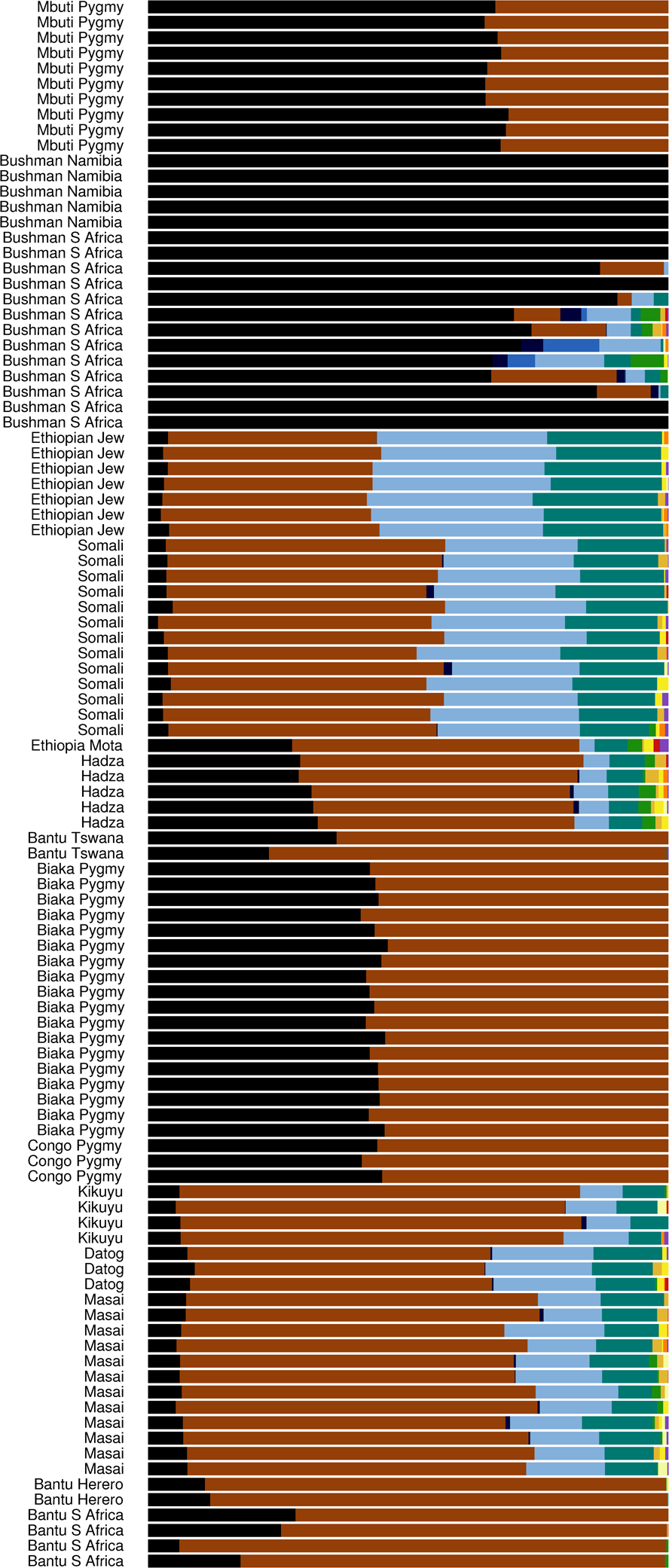
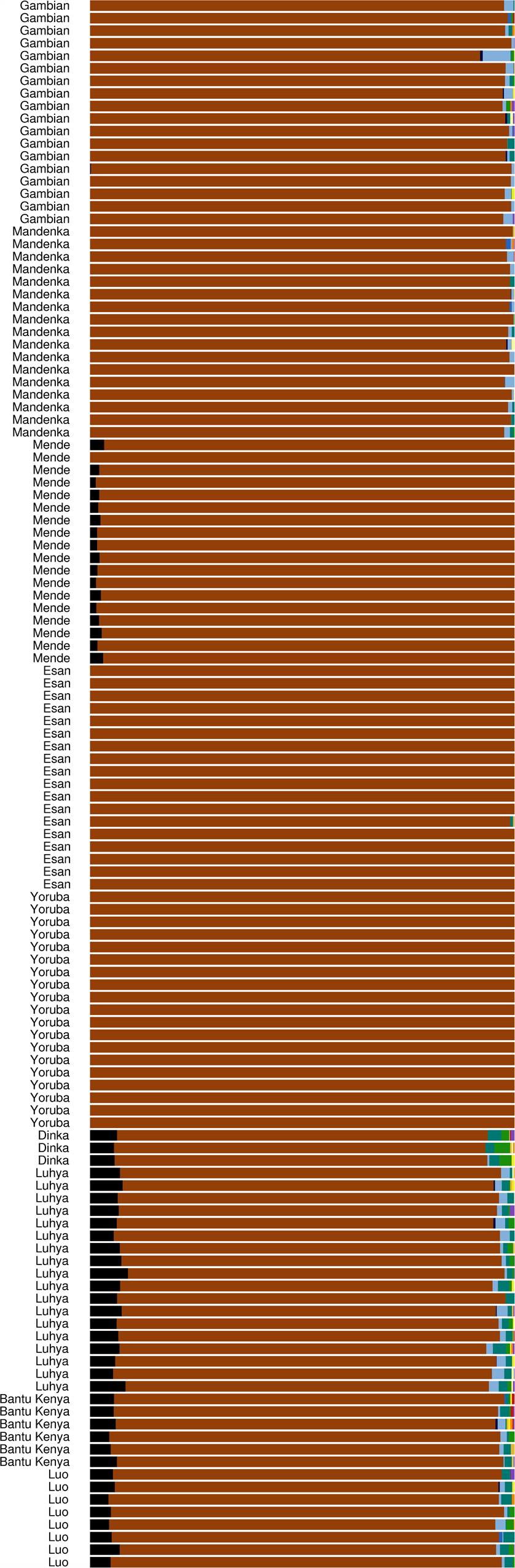
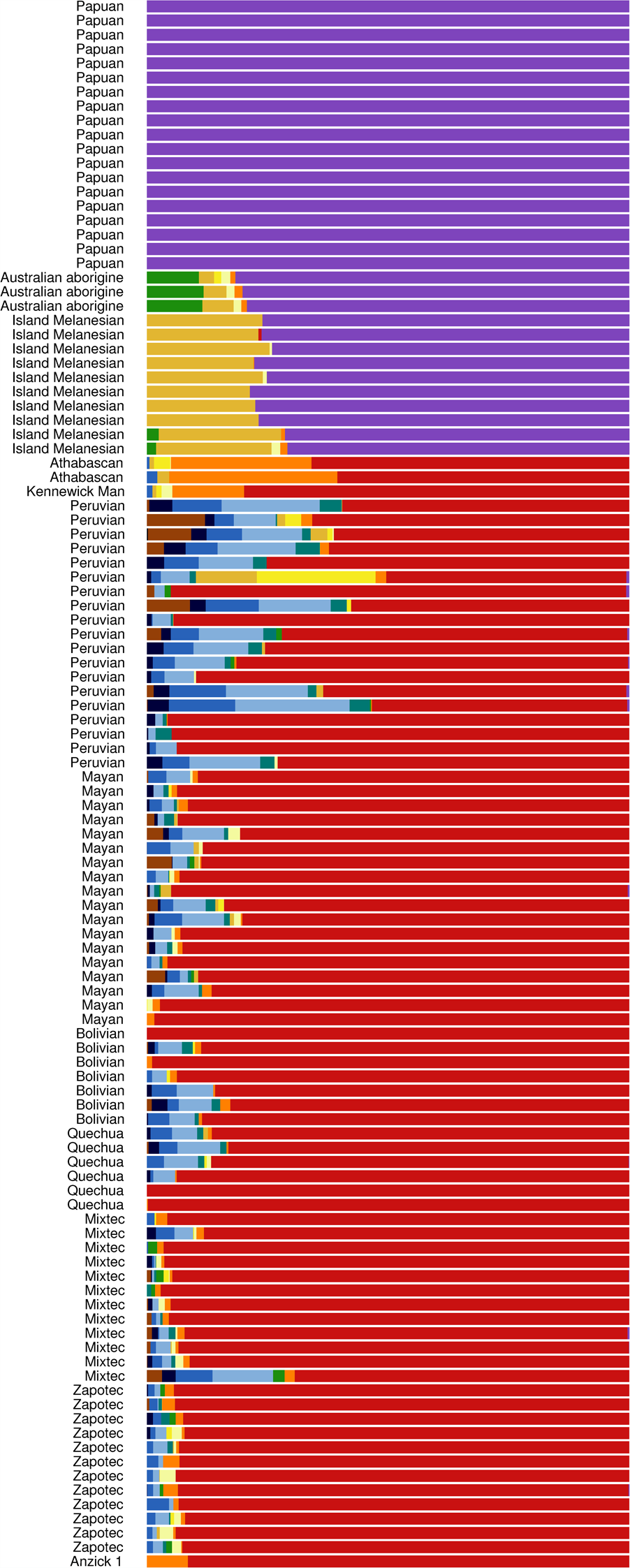
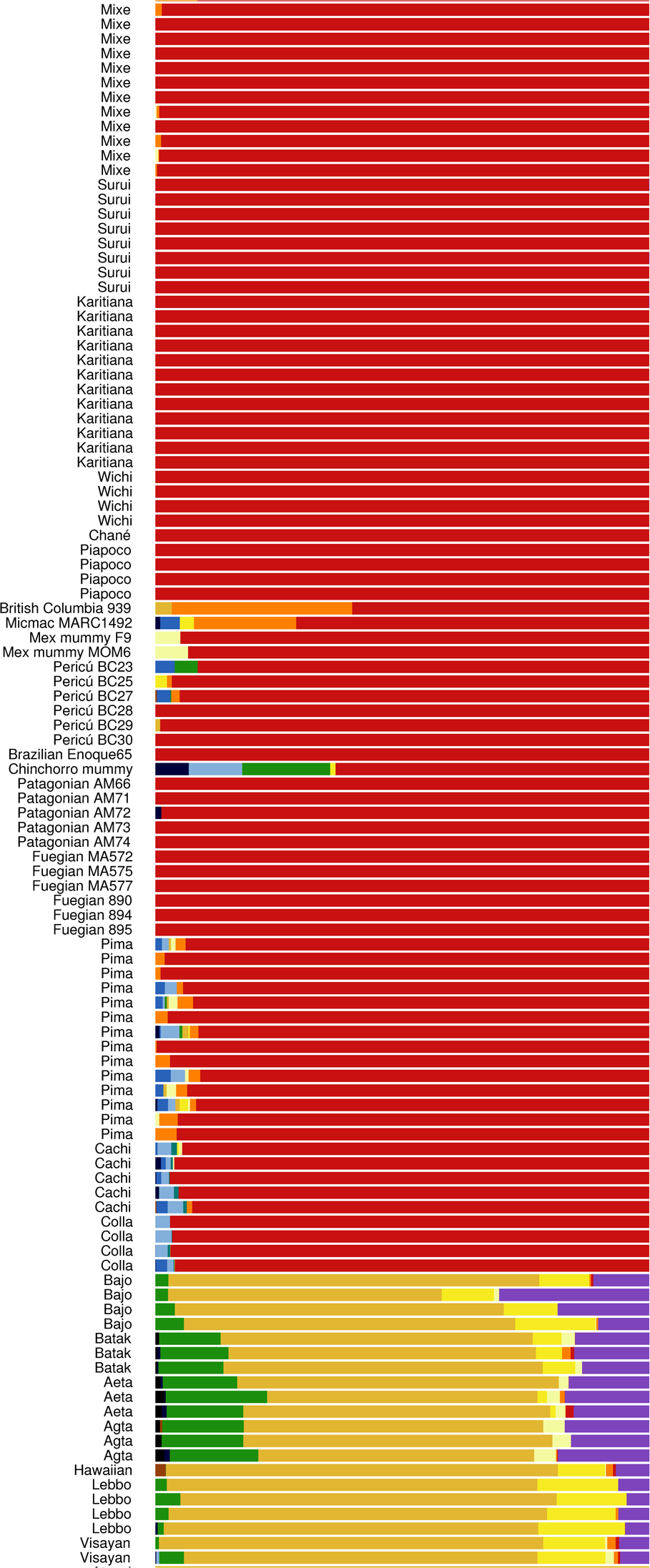
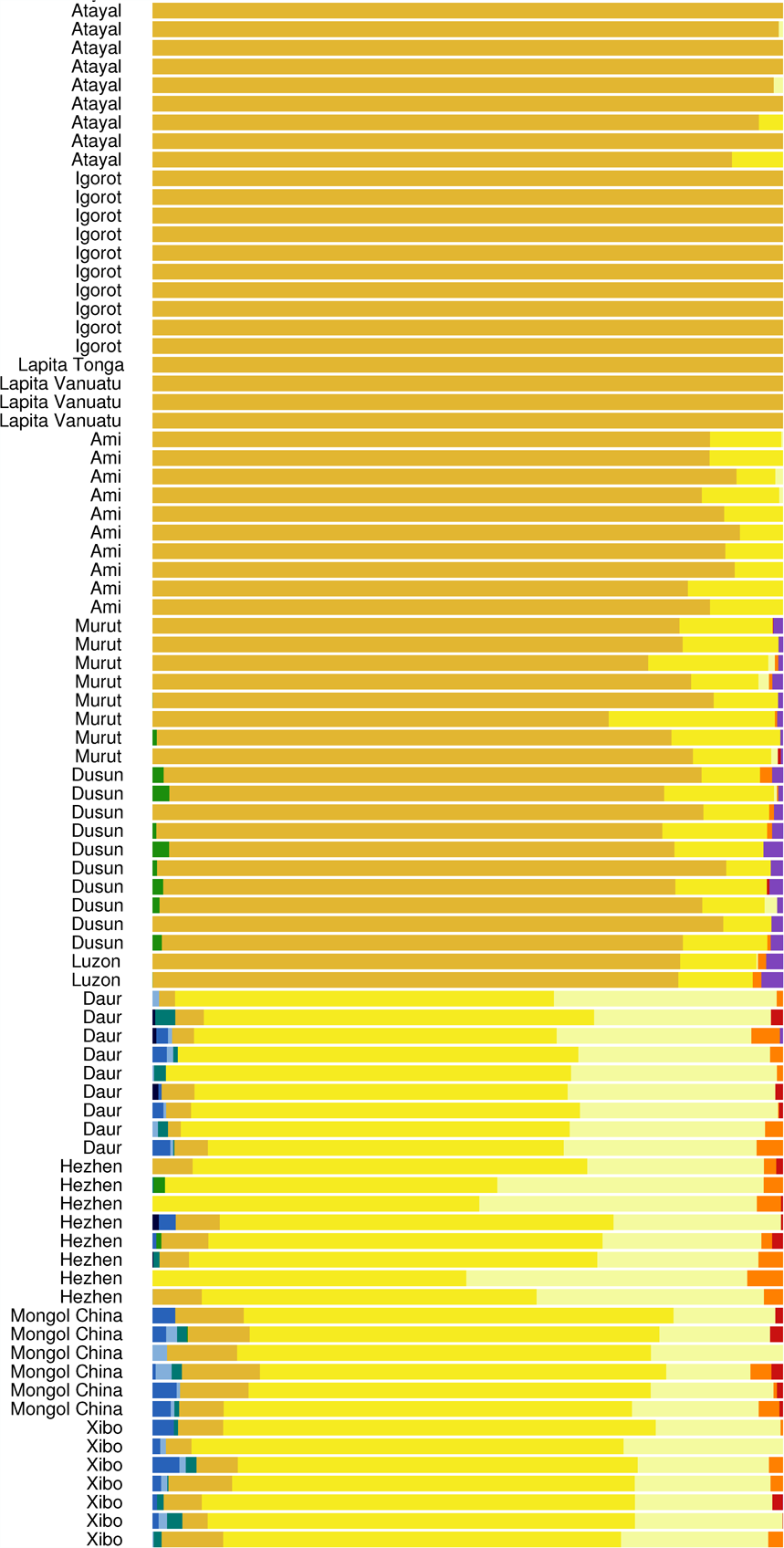
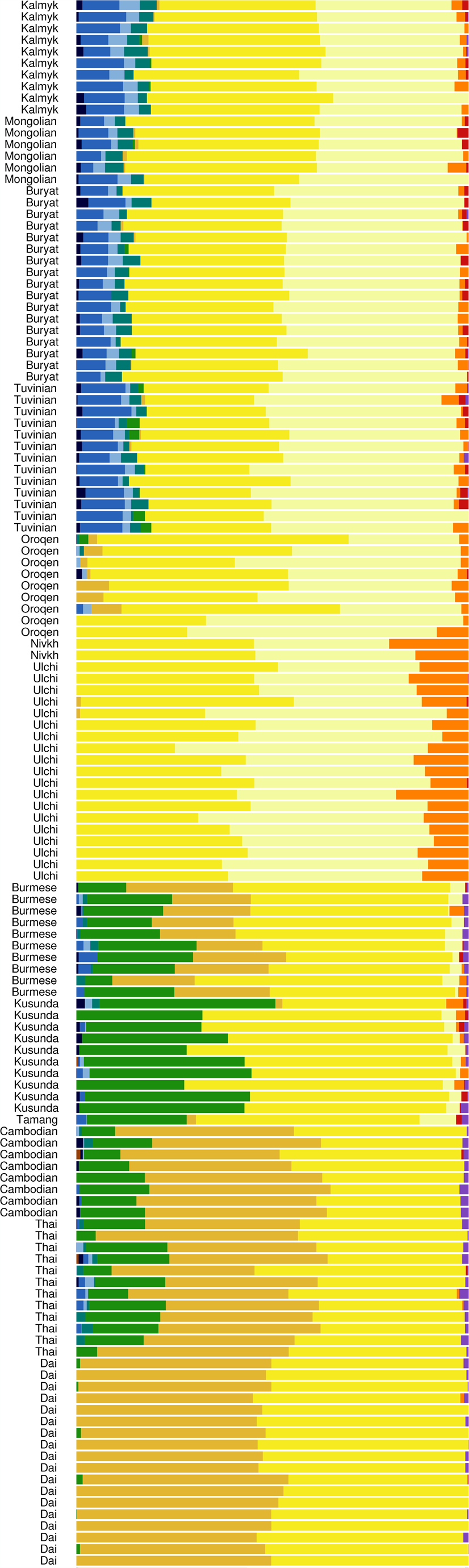
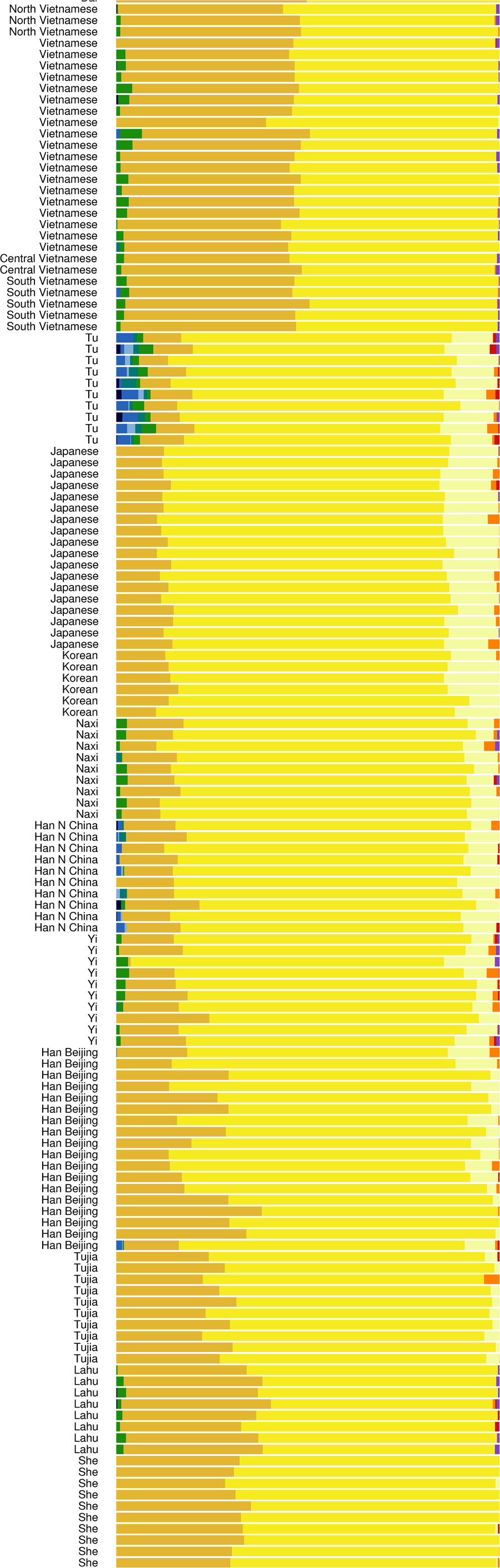
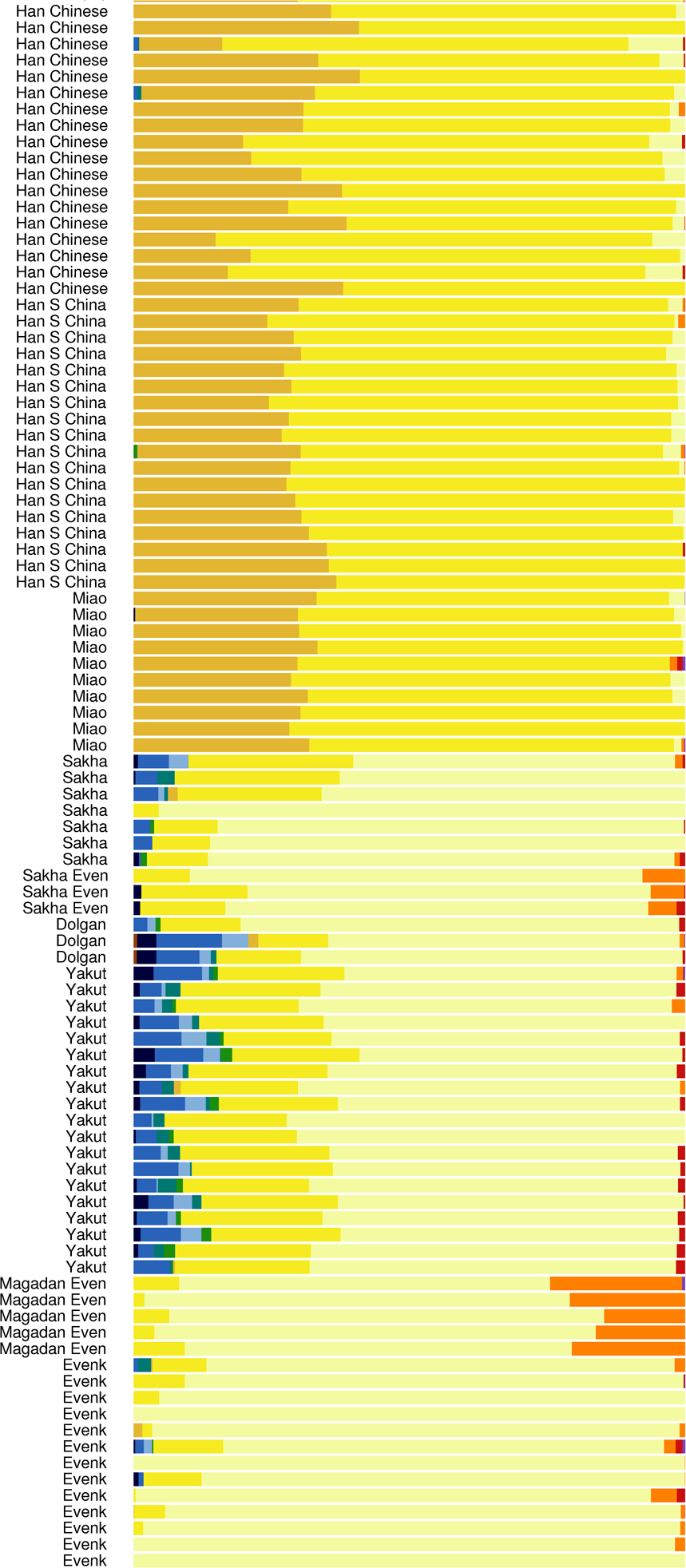
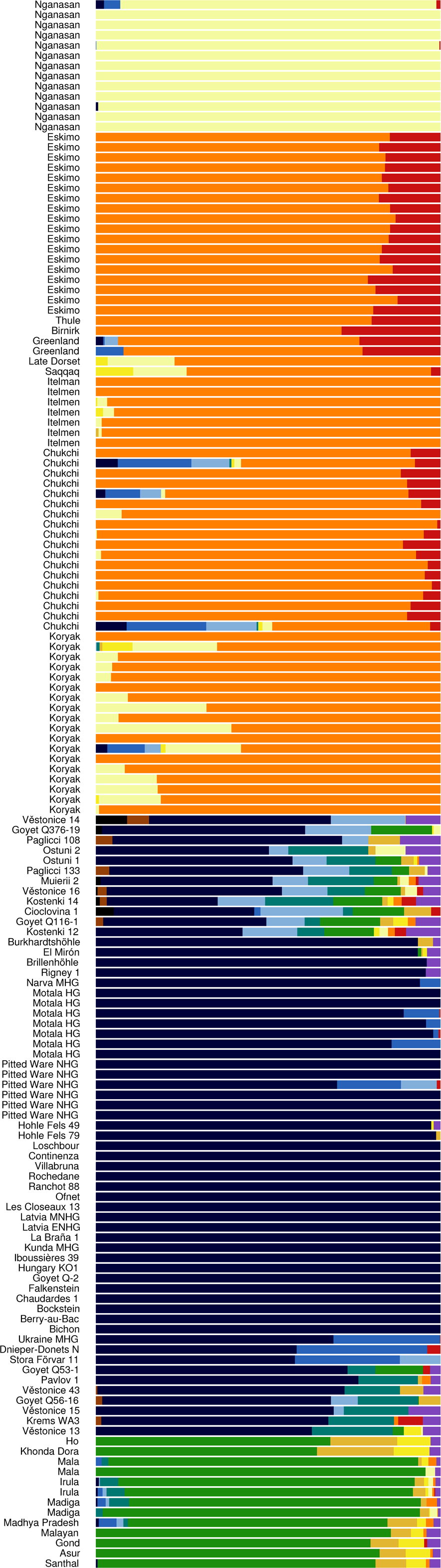
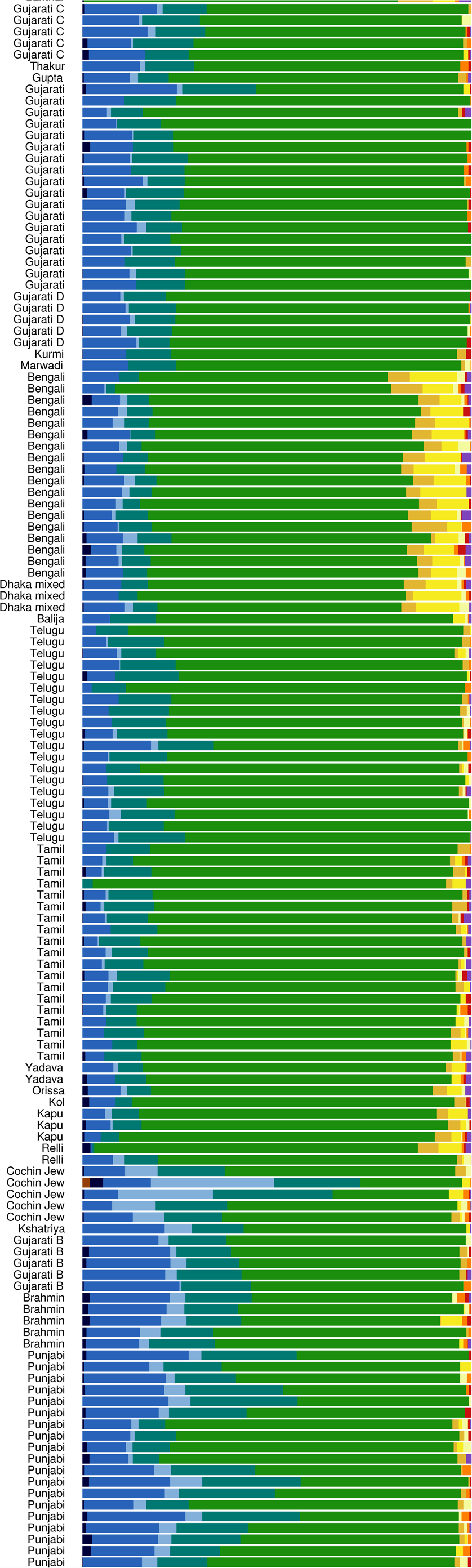
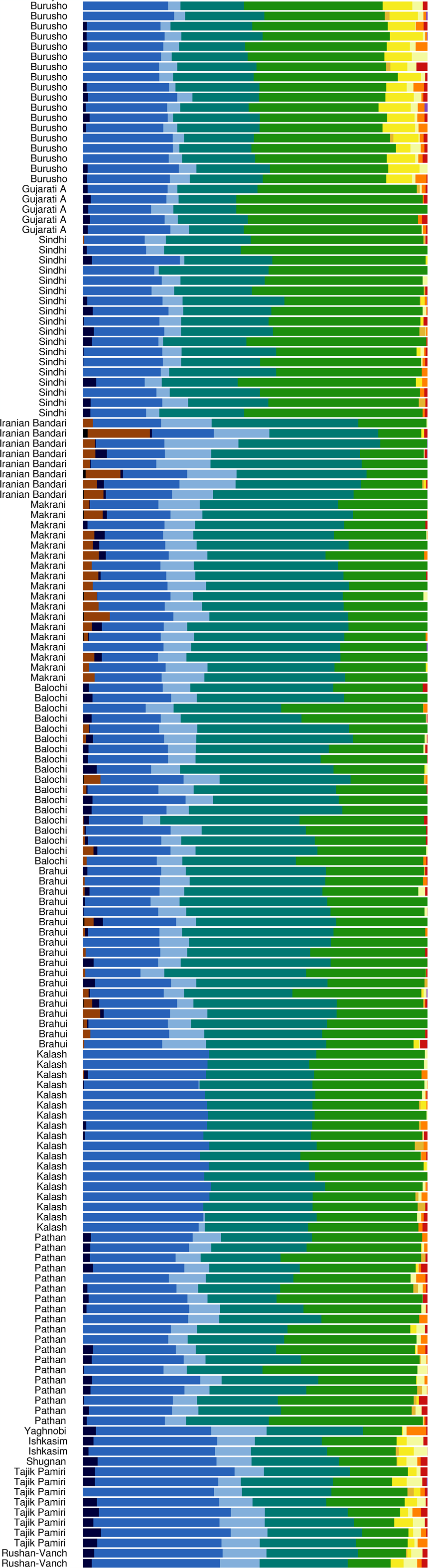
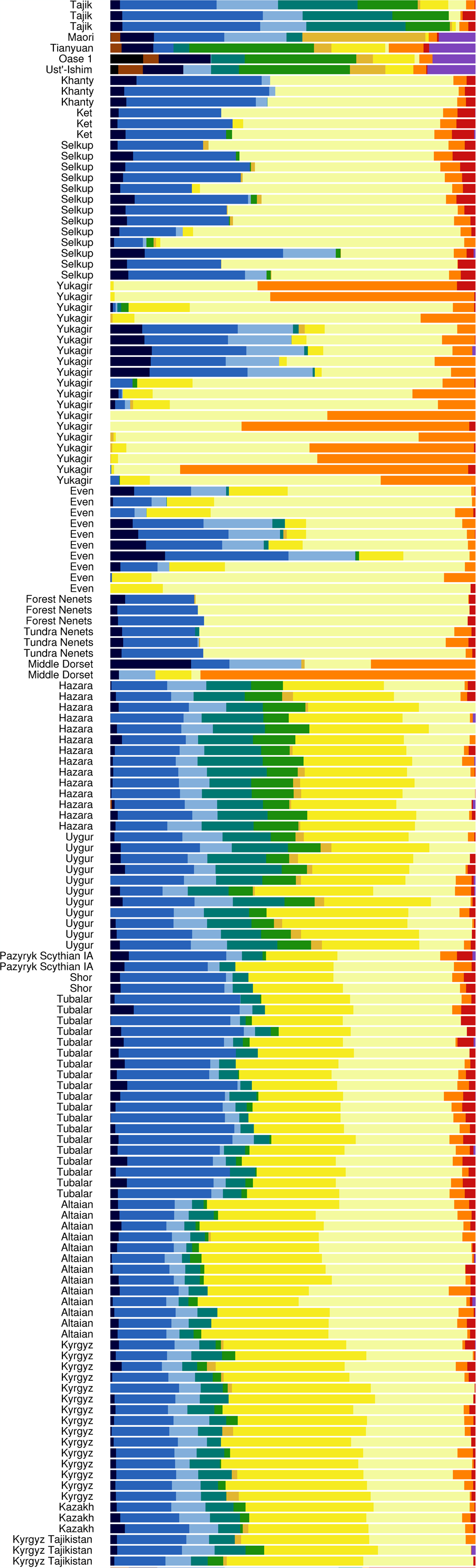
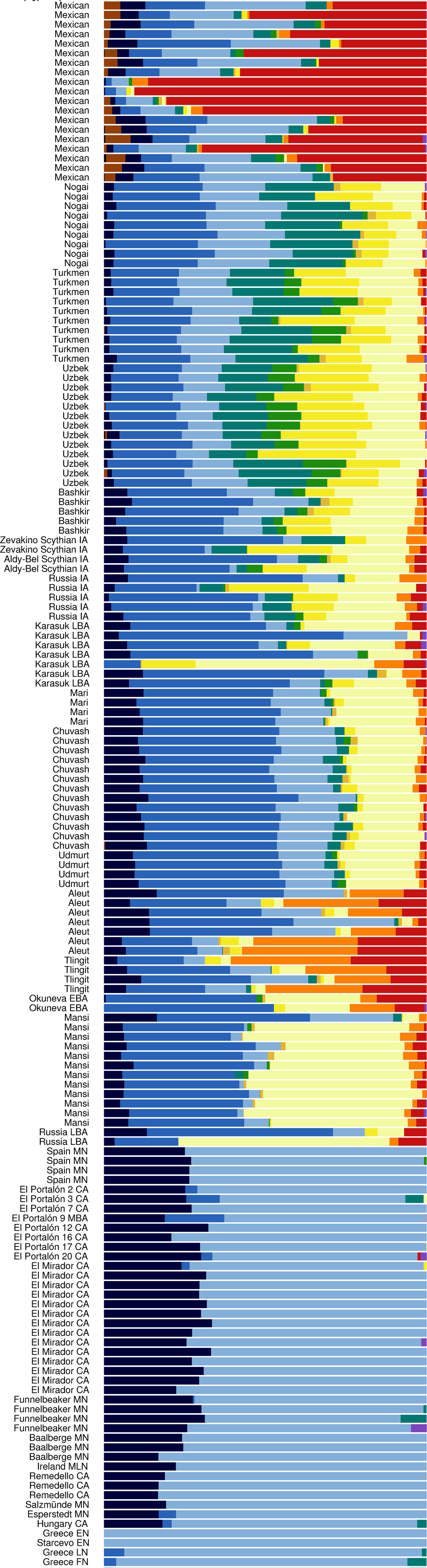
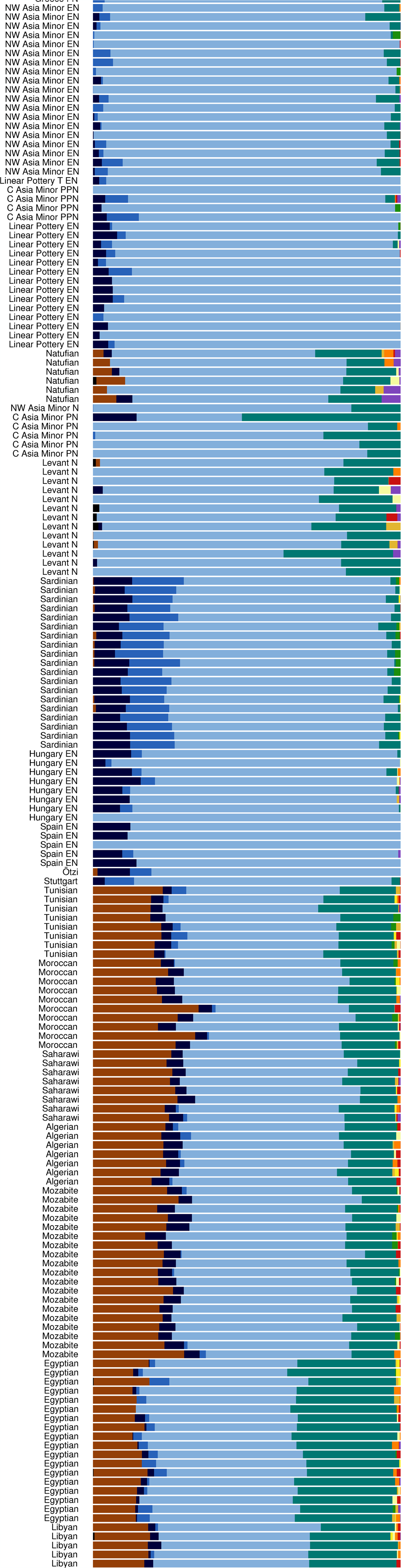
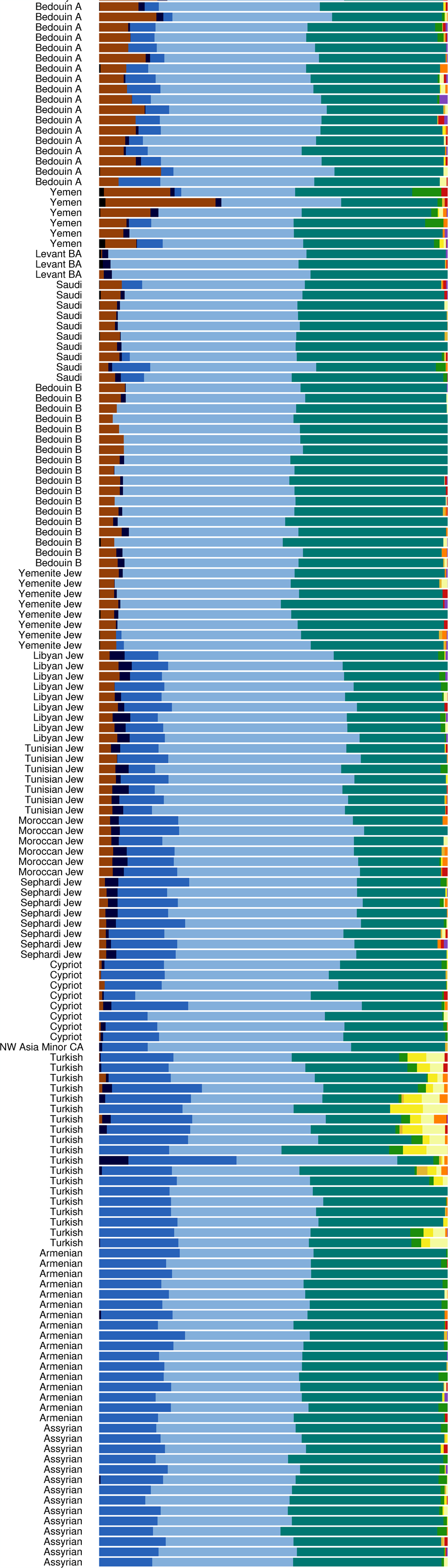
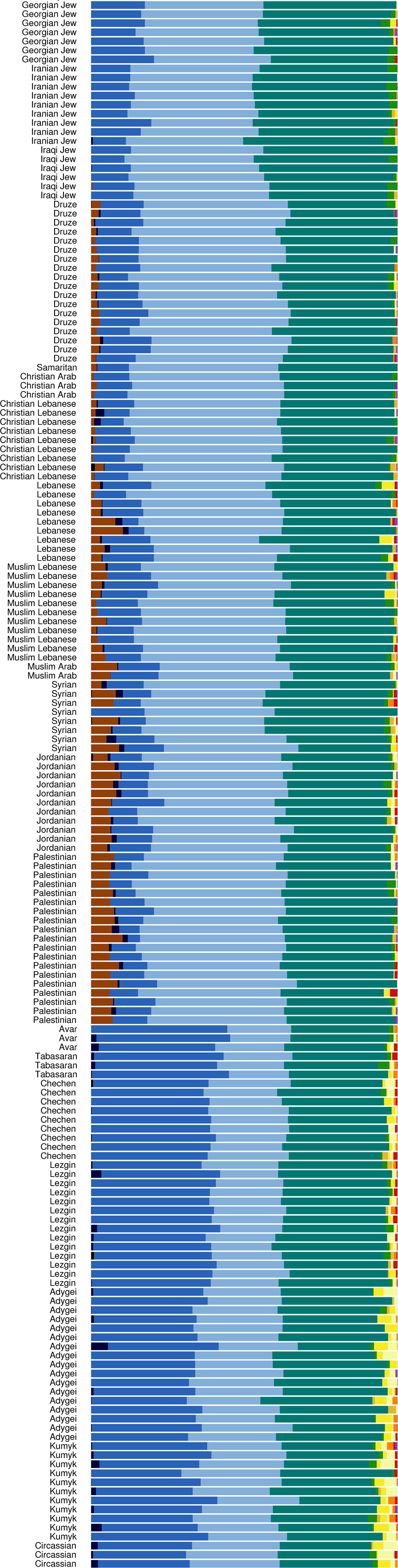
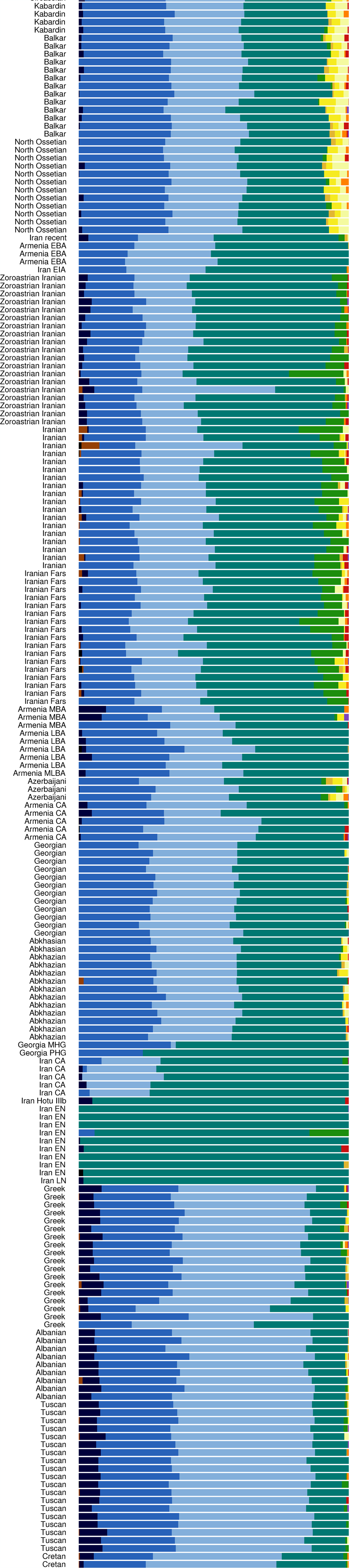
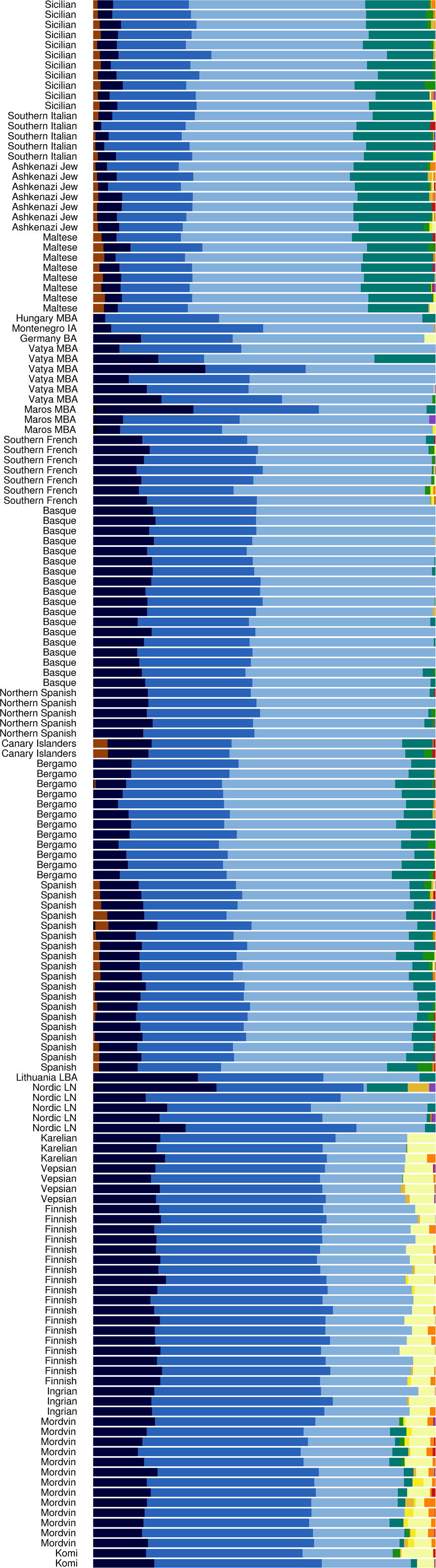
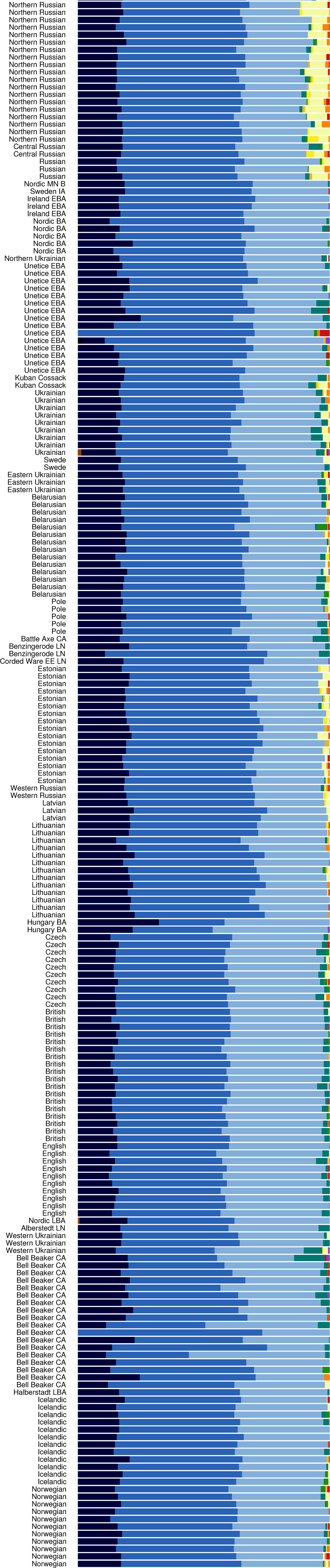
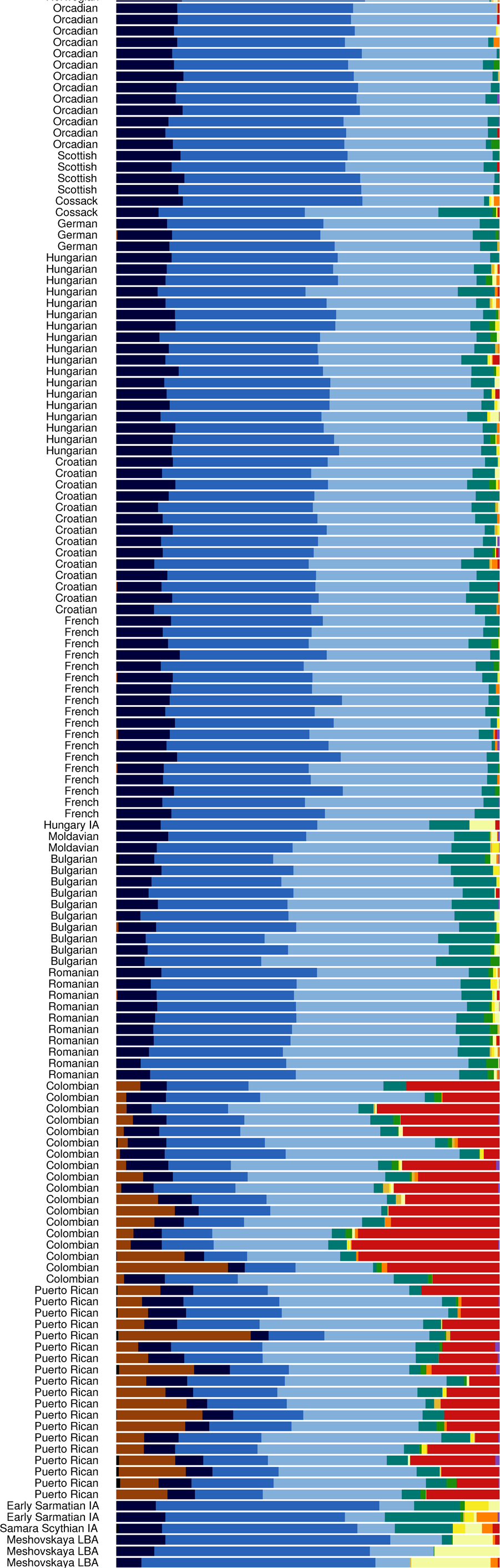
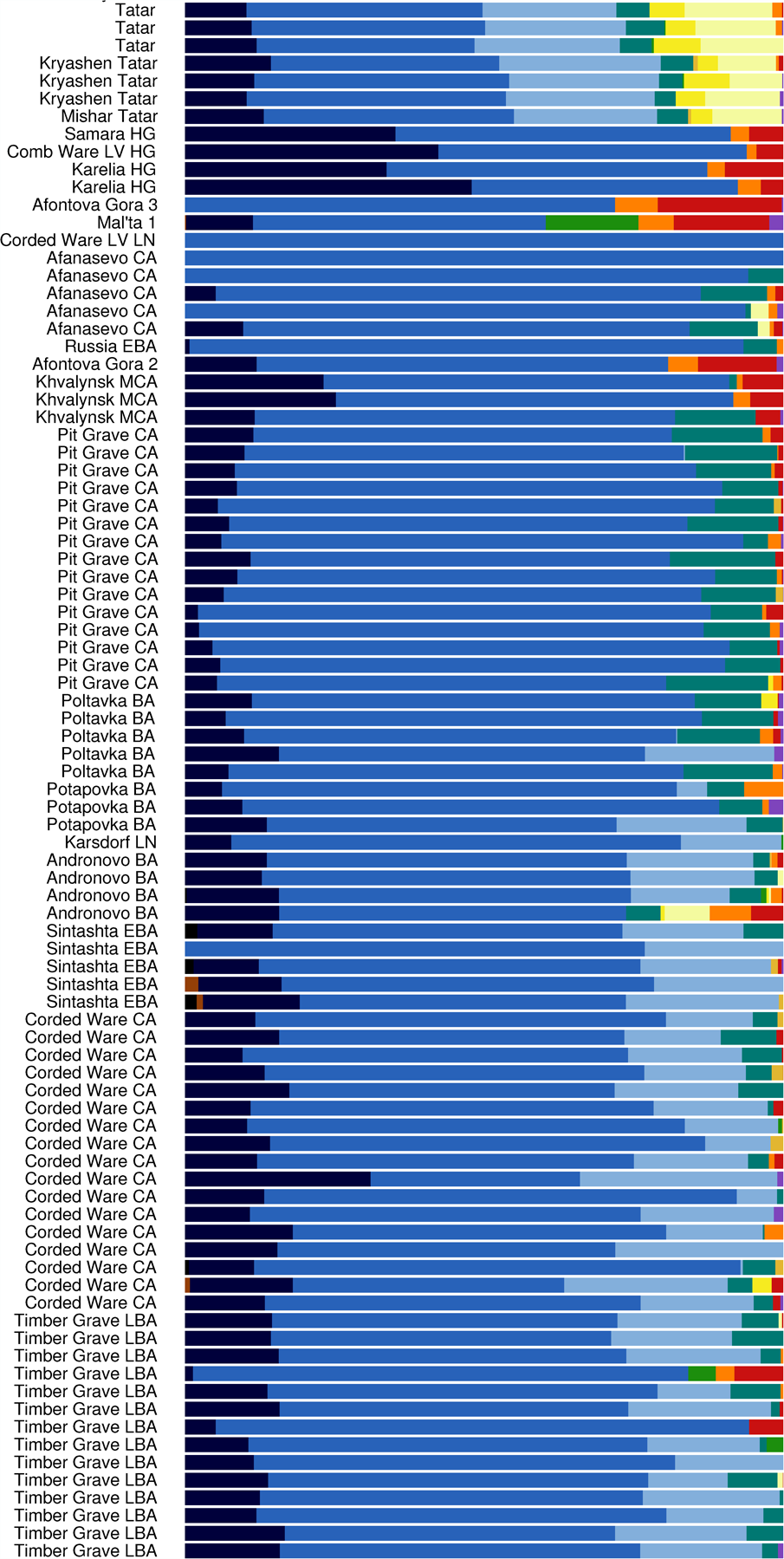
**K = 13 ADMIXTURE analysis**

## Discussion

### *qpAdm* analysis

In Table 2, the 11 analyses in which the Chinchorro mummy sample was the target population have the 11 largest European, Middle Eastern, or South Asian mixture coefficient lower bounds. The mean coefficients for the Chinchorro sample tend to be around 0.45, with the lower bounds mostly around 0.30.

### Principal component analysis

In Figure 1, some of the Mayan, Bolivian, and Quechua samples, which ADMIXTURE analyses show to contain up to 17% European admixture, are shifted to the right of the more pure Amerindian samples, toward the European and Middle Eastern samples. To the right of them is a Mixtec sample which is 23.5% European, and to the right of that Mixtec sample is the 30–40% European Chinchorro mummy sample, and also the Pericú sample BC23, which ADMIXTURE analyses also show to have a significant amount of European admixture. Note that in Figure 1 the Chinchorro mummy sample has a positive value of the second principal component, making it shifted in the direction of Europeans rather than Middle Easterners. In Figure 2 the positions of the Amerindian samples are similar, but BC23 is horizontally between the Mixtec sample and the Chinchorro mummy sample.

### ADMIXTURE analysis

In Figures 3 through 8, for all of the different values of *K*, the Chinchorro mummy sample consistently shows between 30% and 40% non-Amerindian admixture, and the only other samples that show a pattern of non-Amerindian components similar to that seen in the Chinchorro sample are the European samples from before the Last Glacial Maximum (LGM):
- *K* = 4: The Chinchorro sample is 31.66% non-Amerindian. It has a large amount of the blue component, and a small amount of the yellow component. The pre-LGM European samples also have large amounts of the blue component, and smaller amounts of the yellow component.
- *K* = 5: The Chinchorro sample is 32.13% non-Amerindian. It has a large amount of the blue component, a very small amount of the yellow component, and some of the purple component. The pre-LGM European samples also have large amounts of the blue component, and smaller amounts of the yellow and purple components.
- *K* = 6: The Chinchorro sample is 32.67% non-Amerindian. It has large amounts of the blue and green components, and small amounts of the yellow and purple components. The pre-LGM European samples also have large amounts of the blue component, and smaller amounts of the yellow and purple components, and some of them, particularly Kostenki 14 and the Gravettian sample Goyet Q53-1 from Belgium, have some of the green component.
- *K* = 7: The Chinchorro sample is 34.30% non-Amerindian. It has significant amounts of the plain blue, light blue, and green components, and a small amount of the purple component. The pre-LGM European samples also have significant amounts of the plain blue, light blue, green, and purple components.
- *K* = 9: The Chinchorro sample is 37.00% non-Amerindian. It has significant amounts of the plain blue, light blue, and plain green components. The pre-LGM European samples also have significant amounts of the plain blue, light blue, and plain green components. The pine green component is completely absent in the Chinchorro sample, and, with the exception of Kostenki 14, it is also completely absent in the pre-LGM European samples. The plain green component is absent in post-Magdalenian Europeans, and the pine green component is present in significant amounts in Europeans from the Copper Age on, which eliminates modern contamination as a possible source of the European admixture in the Chinchorro sample.
- *K* = 13: The Chinchorro sample is 36.49% non-Amerindian. The Chinchorro sample has significant amounts of the dark blue, light blue, and plain green components, and a small amount of the plain yellow component. The pre-LGM European samples also have significant amounts of the dark blue, light blue, and plain green components, and small amounts of the plain yellow component. The medium blue component is completely absent in the Chinchorro sample, and since the Eastern European and Western Siberian hunter-gatherers are made up mostly of that component, they are excluded as sources of the European admixture in the Chinchorro sample.

## Conclusions

The above *qpAdm*, principal component, and ADMIXTURE analyses of 23 ancient American samples reveal that one of them, taken from a Chinchorro mummy of northern Chile dated to 3972–3806 BC, contains 30–40% European admixture. That the non-Amerindian admixture present in the Chinchorro sample is more closely related to Europeans than to Middle Easterners is demonstrated by both the principal component and ADMIXTURE analyses. The ADMIXTURE analyses further shed light on exactly which European population the non-Amerindian admixture in the Chinchorro sample might be from: only the pre-LGM Europeans show a pattern of non-Amerindian components similar to that seen in the Chinchorro sample, which strongly suggests that the pre-LGM Aurignacians or Gravettians, or possibly the LGM Solutreans, were the source of the admixture. The Solutreans seem like a particularly likely source population, in light of the ample archeological evidence for their presence in the Americas. A complicating factor in identifying the exact source of the European admixture in the Chinchorro sample is the amount of divergent genetic drift that would have occurred between the arrival of the European source population in the Americas and the time of the Chinchorro individual analyzed. If the Solutreans were the source, and if they arrived in the Americas around 26,000 years ago, then around 20,000 years of divergent genetic drift would have accumulated in their American descendants by the time of the Chinchorro individual. This drift might account for the differences in the exact proportions of the non-Amerindian components in the Chinchorro and pre-LGM European samples, but it could also conceivably result in a somewhat later European population, such as the Magdalenians, not being correctly identified as the true source population. Regardless of the exact source of the European admixture in the Chinchorro sample, the fact that such admixture exists is the first ancient DNA proof of pre-Norse transatlantic contact.

